# Adaptation to climate change through dense planting for sustainable agriculture

**DOI:** 10.1101/2022.06.14.496193

**Authors:** Zhixiong Huang, Xue He, Xueqiang Zhao, Wan Teng, Mengyun Hu, Hui Li, Yijing Zhang, Yiping Tong

**Affiliations:** State Key Laboratory of Plant Cell and Chromosome Engineering, Institute of Genetics and Developmental Biology, Chinese Academy of Sciences, Beijing 100101, China; College of Life Sciences, University of Chinese Academy of Sciences, Beijing 100049, China; Hebei Laboratory of Crop Genetics and Breeding, Institute of Cereal and Oil Crops, Hebei Academy of Agriculture and Forestry Sciences, Shijiazhuang, China; State Key Laboratory of Genetic Engineering, Collaborative Innovation Center of Genetics and Development, Department of Biochemistry, Institute of Plant Biology, School of Life Sciences, Fudan University, Shanghai 200438, China; The Innovative Academy of Seed Design, Chinese Academy of Sciences, Beijing, China; College of Advanced Agricultural Sciences, University of Chinese Academy of Sciences, Beijing 100049, China

## Abstract

Overuse of fertilizers increased greenhouse gases emissions, induced global climate changes and extreme weather and made future agriculture unsustainable. Engineering crops to adapt to stressed conditions is crucial. Here, we cloned a transcription factor *TabZIP45* (basic region zipper), controlled by a microRNA binding site polymorphism, conferring adaptation to both nitrogen deficiency and dense planting. TabZIP45 interacted with TaFTL43 (Flowering locus T like43) to change gene expression regulation. TabZIP45 coordinated phosphatidylinositol diphosphate (PIP2) metabolism and calcium (Ca^2+^) signaling to adapt to environmental stresses. Knockout of *TabZIP45-4B* by genome editing rescued grain yield loss caused by nitrogen deficiency by modulation of *TaDwarf4* under dense planting through Ca^2+^ signaling disruption. Thus, *TabZIP45-4B* edited wheat warranted a sustainable and environmentally friendly way to enhance grain yield under adverse conditions.

**One-Sentence Summary:** Calcium and lipids integrated adverse environmental signaling to modulate plant growth

---

The ever-increasing global population demands more cereal grains. However, climate change and recently extreme weather decreased crops yield and endangered human race survival. Global climate change resulted in temperature disturbance and drought, and over use of fertilizer after green revolution deteriorated the climate by greenhouse gases emissions mainly in form of nitrous oxide (N_2_O), which has 1000 times greenhouse effects than that of carbon dioxide (CO_2_) (*1*). To cope with diverse stresses, plants employ complex signaling pathways through interaction among several phytohormones (*2*). Dense planting integrates light, nutrients, phytohormones [such as Brassinosteroids (BRs)] and water signaling to regulated crops growth and ultimately affects grain yield (*3*, *4*). However, the molecular mechanism of how plant integrates environmental signals to modulate growth remains unknown. Interestingly, different stresses signals converge on calcium signaling that plays indispensable roles in regulation plant growth. Calcium signaling is inextricably linked with phospholipids signaling in both animal and plant cell (*5*). Phospholipids play a key role in signal transduction, besides being a major component of cell membranes (*6*). Here, we showed that *TabZIP45-4B*, a *basic region zipper (bZIP*) transcription factor coordinated plant growth in response to environmental stresses by modulation of (Phosphatidylinositol diphosphate) PIP2 - Ca^2+^ signaling. We further demonstrated that dense planting of *TabZIP45-4B* mutant attenuated grain yield loss caused by nitrogen input limitation through crosstalk with BRs signaling.

## Stresses diminished plant fitness

Plants are sessile and have to adapt to local environment to survive which contributed to evolution. Stressed conditions shorten plant life-cycle and shrink plant architecture. Moreover, adverse conditions in agriculture cultivation decrease yield. Nitrogen (N) is one of the essential macronutrients for crops. Wheat spike number (Fig. 1A) and grain yield (Fig. 1B) increases with nitrogen input. Since 1960s, the green revolution increased grain yield under high input of fertilizers. However, overusing of nitrogen fertilizer led to serious environmental pollutions and made agriculture not sustainable. Thus, finding an environmentally friendly and sustainable way to increase grain yield is emergent. Yield increase in modern agriculture was partially due to dense planting (*7*). For example, as the planting density increased, the yield of maize grains increased and then decreased after reaching the optimum value (*7*). In wheat, we observed an interaction between nitrogen supply level and sowing density on spike number and grain yield per unit area. The two sowing densities (D145 with 145 seeds m^-2^ and D72.5 with 72.5 seeds m^-2^) did not differ in spike number and grain yield at N270 (270 kg N m^-2^) level, but displayed significantly difference when N supply level was reduced to N90 or to N0 (Fig. 1A and B). As such, sowing density is required to be optimized for maximizing yield according to N availability.

**Fig. 1.**
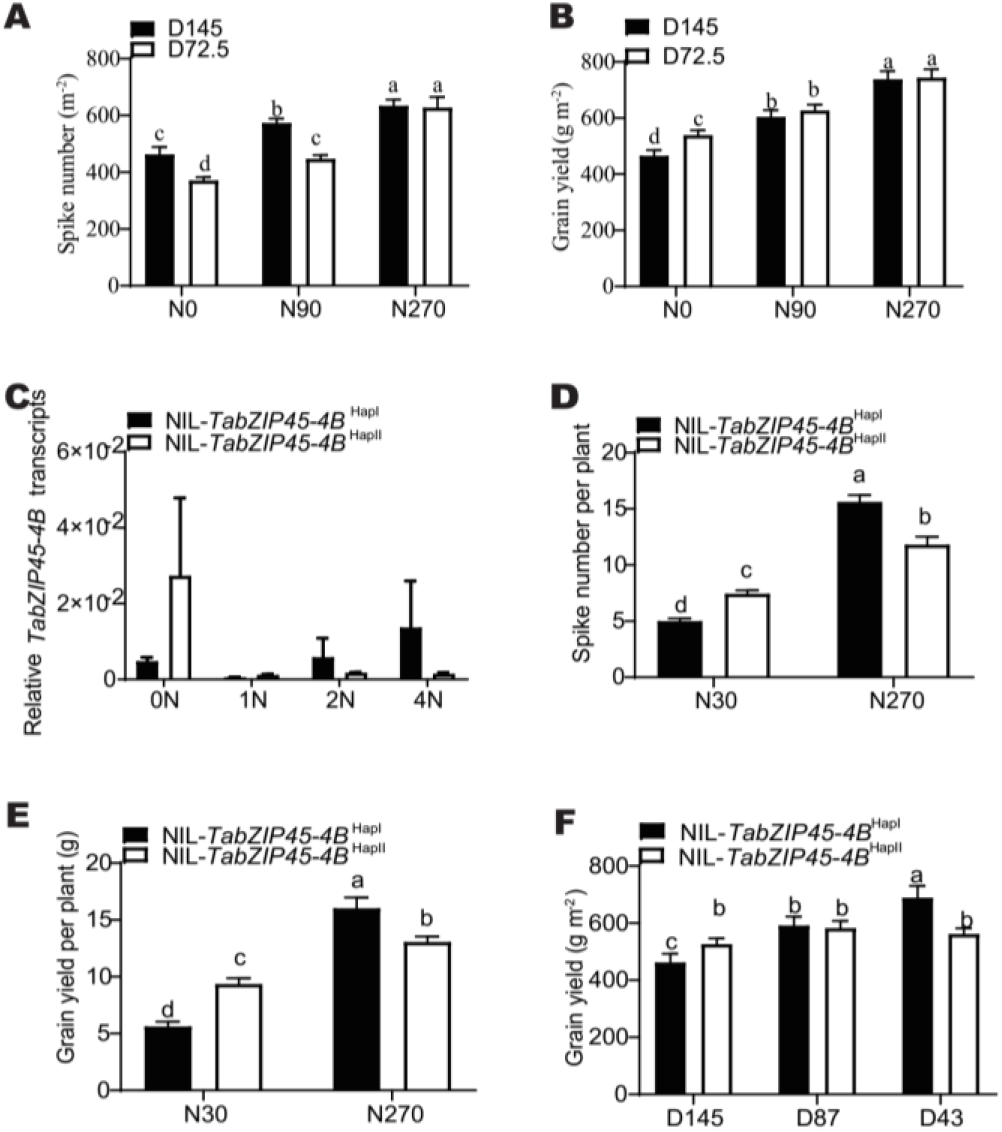
*TabZIP45-4B^HapII^* promoted stresses adaptation. (**A**) Spike number and (**B**) Grain yield per plant of KN199WT (KN199 wild type) under different nitrogen supply levels and sowing densities. Data in A and B are mean ± S. E. (n ≥ 34) (**C**) Relative abundance of *TabZIP45-4B* transcripts in roots. Seedlings were grown at increasing N supply (0.00 mM NH_4_NO_3_, 0N; 0.50 mM NH_4_NO_3_, 1N; 1.0 mM NH_4_NO_3_, 2N; 2.0 mM NH_4_NO_3_, 4N). Isogenic lines *TabZIP45-4B*^HapI^ and *TabZIP45-4B^HapII^* are generated by introducing *TabZIP45-4B*^HapII^ from J411 into XY54 (BC5F6). Data in C are mean ± S. E. (n = 3) (**D**) Spike number and (**E**) Grain yield per plant of *NIL-TabZIP45-4B*^HapI^ and NIL-*TabZIP45-4B*^HapII^ under different nitrogen supply. Data in D and E are mean ± S. E. (n ≥ 15) (**F**) Grain yield of NIL-*TabZIP45-4B*^HapI^ and NIL-*TabZIP45-4B*^HapII^ under different planting densities. Data in F are mean ± S. E. (n ≥ 15). *P* values are calculated by two tailed unpaired *t* test. In (A-B), (D-E), different letters denote statistically significant differences (*P* < 0.05) by two-way ANOVA multiple comparisons. N0 (0 kg N/ha), N30 (30 kg N/ha), N90 (90 kg N/ha), N270 (270 kg N/ha). D43 (43 seeds/m^2^), D72.5 (72.5 seeds/m^2^), D87 (87 seeds/m^2^), D145 (145 seeds/m^2^). All data are from at least three independent experiments.

To isolate genes that response to nitrogen availability and planting density, we constructed a BC_5_F_5_-6 population derived from backcross hybrid between the winter wheat varieties Xiaoyan 54 (XY54) and Jing411 (J411) which were different in nitrogen use efficiency (*8*). The genomic region neighboring centromere in chromosome 4B was significantly associated with response to nitrogen supply by genetics-spike number (per plant) linkage mapping analysis under nitrogen deficient conditions (fig. S1A). We isolated a pair of near-isogenic lines (NILs) from this backcross population that was heterogenous in the region. We further narrowed down the genetic region in the heterogenous offsprings and next sequenced genomic sequences in the region between near-isogenic lines. We found a single-nucleotide polymorphism (SNP) in a gene encoding a basic region zipper (bZIP) transcription factor (fig. S2). The SNP from G (*NIL-TabZIP45-4B^HapI^*) to T (*NIL-TabZIP45-4B^HapII^*) led to a missense mutation from glycine to valine (fig. S1–S2). This glycine is conserved among different plant species (fig. S2). The SNP in haplotype II disrupted the binding sites of a predicted miR5181c-p (fig. S1B). Indeed, we found that *TabZIP45-4B* differentially responded to nitrogen availability between NIL*-TabZIP45-4B*^HapI^ and NIL-*TabZIP45-4B*^HapII^ in shoots of seedlings (Fig. 1C). In addition, *TabZIP45* (also known as *TGA2*) was associated with diverse roles in nutrient sensing, stresses adaptation and growth development (fig. S3). Thus, we supposed that *TabZIP45* regulated nitrogen deficiency adaptation probably by growth modulation.

Indeed, three independently paired NILs showed that NIL-*TabZIP45-4B*^HapII^ significantly increased spike number, grain yield per plant and photosynthesis rate compared to NIL-*TabZIP45-4B*^HapI^ under nitrogen deficient conditions (fig. S4A to D). *NIL-TabZIP45-4B^HapII^* slightly increased plant height (fig. S4E) and grain number per spike (fig. S4 F) than *NIL-TabZIP45-4B^HapI^* without affecting 1000-grain weight (fig. S4 G). The decrease in grain yield and spike number per plant caused by low nitrogen input was greater in *NIL-TabZIP45-4B^HapI^* than NIL-*TabZIP45-4B*^HapII^ (Fig. 1D and E). Thus *NIL-TabZIP45-4B^HapI^* was more sensitive to nitrogen availability compared with *NIL-TabZIP45-4B^HapII^*.

We next explored whether the nitrogen competition among plants was similar to deficient nitrogen supply. The decline of sowing densities from D145 (145 seeds m^-2^) to D43 (43 seeds m^-2^) significantly increased grain yield in NIL-*TabZIP45-4B^HapI^* (Fig. 1F), whereas grain number of NIL-*TabZIP45-4B*^HapII^ was relatively stable in different sowing densities (Fig. 1F). In addition, the grain yield of NIL-*TabZIP45-4B^HapII^* was higher than that of NIL-*TabZIP45-4B*^HapI^ at D145 (Fig. 1F), but was lower than that of NIL-*TabZIP45-4B*^HapI^ at D43 (Fig. 1F). Compared with NIL-*TabZIP45-4B*^HapI^, NIL-*TabZIP45-4B^HapII^* had a decrease in spike number and 1000-grain weight under all sowing densities investigated, but an increase in grain number per spike (fig. S5 A to C) under relative denser sowing conditions (D145 and D87). In addition, high sowing densities (D145) reduced more grain number (per spike) in NIL-*TabZIP45-4B*^HapI^ compared with NIL-*TabZIP45-4B*^HapII^ (fig. S5B). As such, that NIL-*TabZIP45-4B*^HapII^was less sensitive to planting density than NIL-*TabZIP45-4B*^HapI^. Together, these two NILs differed in the yield traits in response to nitrogen availability and planting densities, and NIL-*TabZIP45-4B*^HapII^ was more sensitive to nitrogen availability and planting densities than NIL-*TabZIP45-4B*^HapI^.

### *TabZIP45-4B* regulated nitrogen response

*TabZIP45-4B* encodes a transcription factor closely related to TGACG DNA-binding factors (TGA) family (fig. S6A). TabZIP45-4B differs with TabZIP45-4A and TabZIP45-4D in amino acids residue site at 178 (fig. S6B and S7A). TabZIP45-4B was localized in nucleus in wheat protoplast (Fig. 2A). We next examined the *TabZIP45* expression in response to different nitrogen availability. The expression of *TabZIP45-4A* and *TabZIP45-4D* was induced by nitrogen deprivation (fig. S8); however, *TabZIP45-4B* transcripts increased under both nitrogen deprivation (0N) and nitrogen over-supply concentration (4N) when compared with 2N treatment (fig. S8). To confirm the role of *TabZIP45-4B* in adaptation to nitrogen deficiency and dense planting, we created the knockout mutants by clustered regularly interspaced short palindromic repeats (CRISPR) cas9 genome editing (fig. S7). During our screening for mutants after sowing, we obtained a heterozygous mutant line of *TabZIP45-4B*. We sequenced *TabZIP45-4B* and found a stop gain mutation in the mutated *tabzip45-4b* (fig. S7). A total of 157 plants from this heterozygous mutant displayed segregation ratio of 38 : 84 : 35 (approximately 1: 2: 1) for null segregant (*TabZIP45-WT*) : heterozygous mutation (*tabzip45*-*Bb*) : homozygous mutation (*tabzip45*-*bb*) (fig. S9A). Compared with *TabZIP45*-WT and *tabzip45*-*Bb*, the increase of grain yield (per plant), spike number (per plant) and grain number (per spike) in *tabzip45*-*bb* (fig. S9A to C) was significant without affecting thousand grain weight (fig. S9D). Furthermore, the two independent lines with *TabZIP45-4B* single gene mutation (*tabzip45*-*bb*) had more spike number and grain yield than wild type control (Fig. 2B and C). We next investigated whether tillering was affected by *TabZIP45-4B* mutation. The *tabzip45-bb* mutant had more tiller buds during early shoot development than negative control *TabZIP45-WT* (fig. S10), indicating that *TabZIP45-4B* influenced shoot bud formation rather than shoot elongation. We then characterized the role of *TabZIP45-4B* in response to different nitrogen supply level. The spike number and grain yield (both *tabzíp45-bb* and *TabZIP45-WT*) were decreased under N90 (90 kg N ha^-1^) conditions compared with N270 (270 kg N ha^-1^) conditions (Fig. 2D to F). Nevertheless, the mutation of *TabZIP45-4B* increased grain yield, spike number and dry biomass under both N90and N270 conditions (Fig. 2D to F, fig. S11A) without significantly affecting on plant height, the grain number (per spike) and 1000-grain weight (fig. S11 B to D). These results suggested that *TabZIP45-4B* was involved in nitrogen response by growth modulation.

**Fig. 2.**
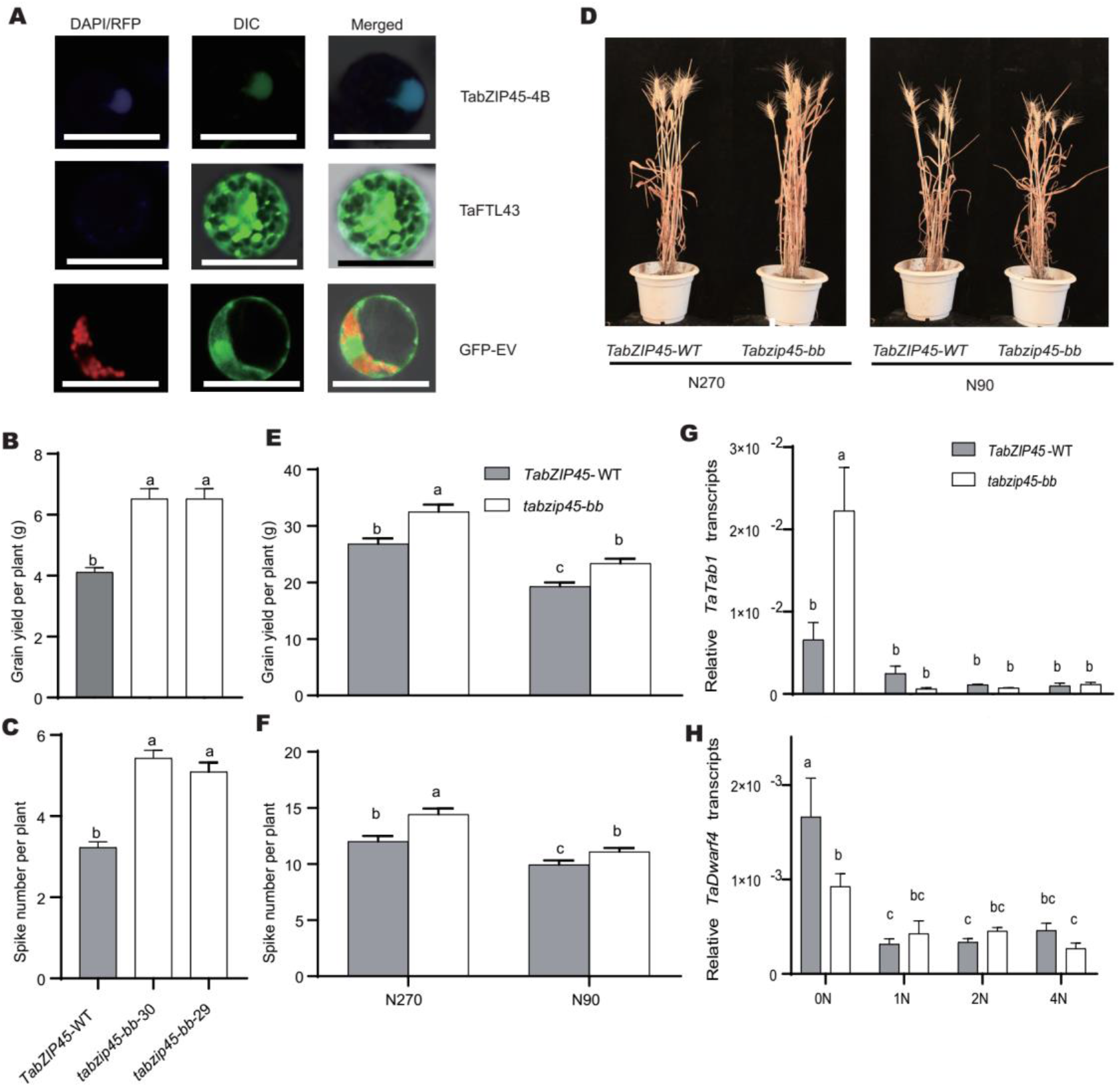
*TabZIP45-4B* regulated spike number in response to nitrogen supply. (**A**) TabZIP45-4B and subcellular localization. Scale bars, 20 μm. (**B**) Grain yield and (**C**) Spike number of *TabZIP45-4B* mutant *tabzip45-bb* and control *TabZIP45-WT* under 30 kg/ha nitrogen supply level. Data in B and C are mean ± S. E. (n ≥ 49). (**D**) mature plants under 270 kg/ha and 90 kg/ha nitrogen supply level. Scale bars, 20 cm. (**E**) Grain yield and (**F**) Spike number of *TabZIP45-4B* mutant *tabzip45-bb* and control *TabZIP45-WT* under 270 kg/ha and 90 kg/ha nitrogen supply level. (**G**) Relative abundance of *TabZIP45-4B* transcripts in roots. Seedlings were grown at increasing N supply (0.00 mM NH_4_NO_3_, 0N; 0.50 mM NH_4_NO_3_, 1N; 1.0 mM NH_4_NO_3_, 2N; 2.0 mM NH_4_NO_3_, 4N). Isogenic lines *TabZIP45-4B^HapI^* and *TabZIP45-4B^HapII^* are generated by introducing *TabZIP45-4B^HapII^* from J411 into XY54 (BC5F6). *P* values are calculated by two tailed unpaired *t* test. Data in G are mean ± S. E. (n ≥ 6) and different letters denote statistically significant differences (*P* < 0.05) by two-way ANOVA multiple comparisons. All data are from at least three independent experiments. In (D-F), N90 (90 kg N/ha), N270 (270 kg N/ha).

To identify the downstream genes expression affected by nitrogen supply level, we analyzed the gene binding sites by chromatin immunoprecipitation (ChIP)-sequencing under nitrogen deficient conditions. The *bZIP* gene family binding motif TGCA box was enriched (fig. S12). Furthermore, *TaTab1*, a *WUSCHEL* ortholog *tab1 (TILLERS ABSENT 1*, responsible for both tiller formation and elongation) (*54*) and *TaDwarf4* [a speed-limiting enzyme in BRs (*TraesCS4B03G0638000*) synthesis] were enriched in sequencing signals. The binding affinity of TabZIP45 to the *TaTab1* and *TaDwarf4* promoter was corroborated by yeast one hybrid and DNA-TabZIP45 pulldown binding assay (fig. S13). We further assessed whether *TabZIP45-4B* regulated *TaTab1* and *TaDwarf4* gene expression. Although no significant *TabZIP45 (TabZIP45-4A, TabZIP45-4B*, and *TabZIP45-4D*) expression difference between *TabZIP45-4B* and *tabzip45-bb* were observed (fig. S14), the expression *TaTab1* in *tabzip45-bb* was significantly increased compared with *TabZIP45-4B* in nitrogen deprivation conditions (0N) (Fig. 2G), whereas *TaDwarf4* in *tabzip45-bb* was significantly decreased compared with *TabZIP45-4B* under 0N (Fig. 2H). Moreover, the difference of both *TaTab1* and *TaDwarf4* gene expression between the two genotypes vanished with increasing nitrogen supply level (Fig. 2G to H). Notably, the *TaTab1* expression in the *TabZIP45-WT* plants and *TaDwarf4* expression in the *tabzip45-bb* plants were relatively stable in different nitrogen supply (Fig. 2G to H). These results indicated that *TaTab1* and *TaDwarf4* may be coordinated by *TabZIP45* to regulate tiller development. Indeed, we found that the heading date between *TabZIP45-4B* near isogenic lines (NILs) under nitrogen deprivation condition was significantly different (approximately one week), which may lead to spike number difference (Fig. 1D, fig. S4C, fig. S5A). Thus, our results indicated that *TabZIP45-4B* regulated tillering in adaption to different nitrogen supply.

## TabZIP45 was targeted by TaFTL43

The life cycle of common wheat is the annual cycle of sowing in autumn, vernalizing in winter and growing to harvest in summer. Different with *Arabidopsis thaliana*, several vernalization factors in common wheat had been investigated (*9*–*11*). One of them was reported to be important in sense nitrogen (*12*). We reasoned that too short time of low temperature duration was not enough for vernalization, but too longtime duration led to freezing stress, both of which diminished spike number. To reveal molecular mechanism that how *TabZIP45* regulate tillering in response to environmental stimulations, we collected tiller samples at tillering stage after sowing under nitrogen deficient conditions. We carried out quantitative real-time polymerase chain reaction (qRT-PCR) assay to examine all *FLOWERING LOCUS T (FT*) family genes expression level (fig. S15). *TaFTL-43 (TaPEBP30*, TraesCS3B02G015200) was the one with the highest transcripts abundance in shoots (fig. S15A). We further assessed the transcriptional level of *TaFTL-43* in leaves and shoot basal parts of the *tabzip45-bb* and *TabZIP45-WT* plants, respectively. Compared with *TabZIP45-WT, TaFTL-43* was downregulated in *tabzip45-bb* shoot basal parts but not in leaves (fig. S15B). Thus, these results indicated that *TaFTL-43* acted with *TabZIP45-4B* to regulate tiller development under nitrogen deficient condition.

Both nitrogen deficiency and high planting density resulted in spike number decrease (Fig. 1A). In addition to the effects of nutrient competition, planting density also affected light intensity and quality of light (the ratio of far-red/red), which regulated FT activity (*13*, *14*) and shoot branching (*15*). We therefore addressed whether *TaFTL-43* responded to planting density. Indeed, compared with D43, *TaFTL-43* was downregulated under D87 in tiller basal parts but not in leaves (fig. S15C). These results indicated that *TaFTL-43* might also was involved in the tiller formation in response to planting density.

*bZIP* transcription factor were the only regulators known so far to work under far-red, red and blue light conditions and to regulate the light signaling pathway (*16*). In *Arabidopsis* and common wheat, FT protein interacted with *bZIP* transcription factor FLOWERING LOCUS D (FD) protein to regulate flowering (*12*, *17*, *18*). We therefore investigated whether TaFTL-43 interacted with TabZIP45-4B. Indeed, bimolecular fluorescence complementation (BIFC) assay indicated that TabZIP45-4B interacted with TaFTL-43 *in vivo* in tobacco (Fig. 3A). The glutathione S-transferase (GST) pulldown assay validated direct interaction between TaFTL-43 and TabZIP45-4B (Fig. 3B). We next examined whether the interaction between TaFTL-43 and TabZIP45-4B is necessary for TabZIP45-4B to regulate downstream genes. At the presence of TaFTL-43, knockout of *TabZIP45-4B* (premature stop gain mutation) increased *TaTab1* expression level (fig. S16), which indicated that disruption the interaction between TaFTL-43 and TabZIP45-4B led to activate *TaTab1* expression. We next determined whether different haplotype of *TabZIP45-4B* transcription factor function was affected by the interaction with TaFTL-43. Unexpectedly, downstream gene *TaTab1* transcription was not significantly changed between the two haplotypes (fig. S16) in all tested condition, which suggested that the variant in the two haplotypes did not affect the interaction between TaFTL-43 and TabZIP45-4B. Indeed, the amino acid residue sites at 196 was far away from interaction Ser-Ala-Pro (SAP) motif (*19*, *20*). These results indicated that TaFTL-43 and TabZIP45-4B were jointly involved in the regulation of downstream genes.

**Fig. 3.**
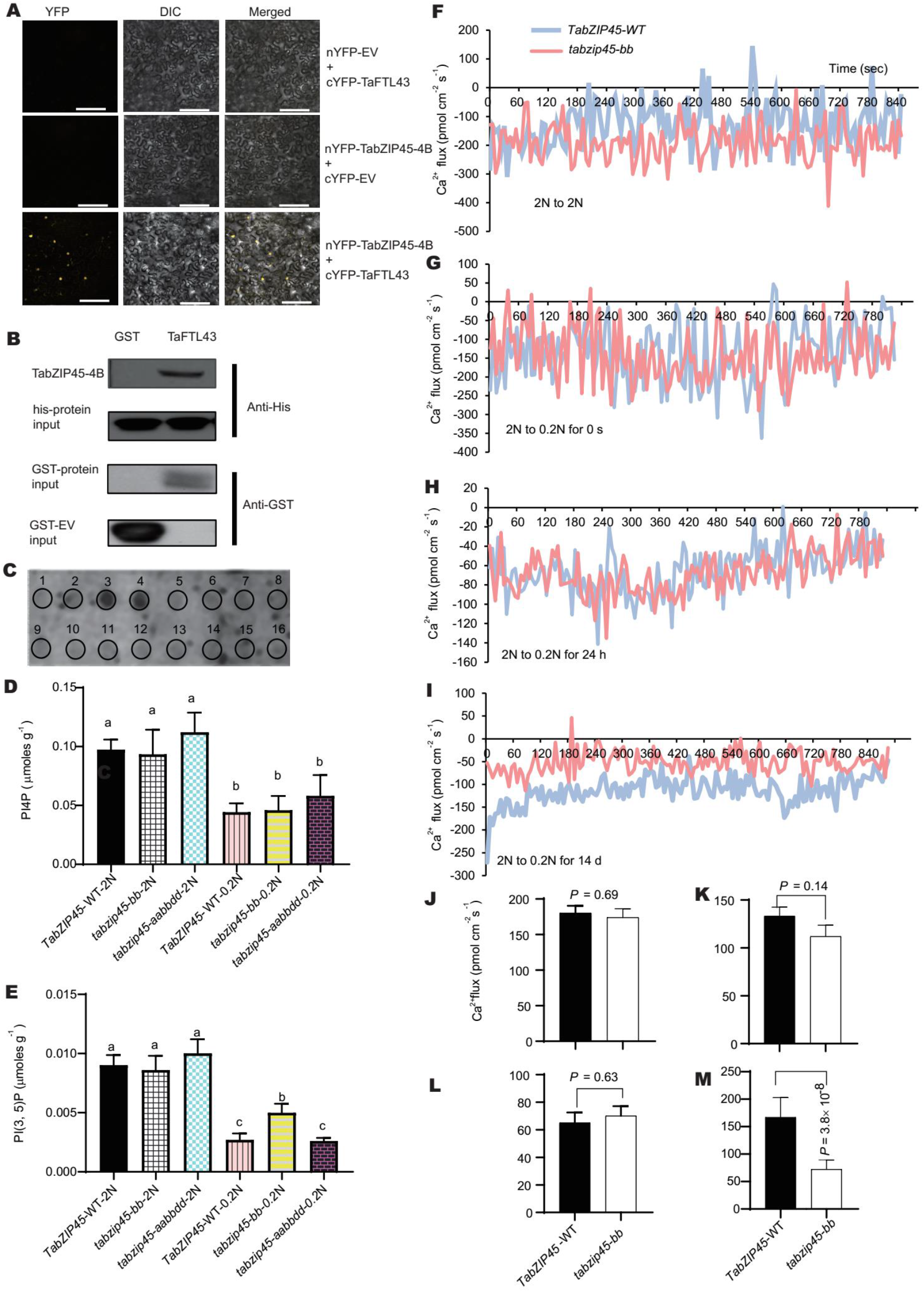
TaFTL43 targeted *TabZIP45-4B* to coordinated PIP2 metabolism and calcium signaling. (**A**) BiFC experiment. Scale bar, 100 μm. (**B**) GST pull down assays. The *in vitro* interaction of TabZIP45-4B and TaFTL-43. (**C**) Dot blot assay. lipids were dotted as indicated (1, phosphatidylinositol; 2, Phosphatidylinositol-4-phosphate [PtdIns(4)P]; 3, Phosphatidylinositol-4,5-bisphosphate [PtdIns(4,5)P2]; 4, Phosphatidylinositol-3,4,5-trisphosphate [PtdIns(3,4,5)P3]; 5, Cholesterol; 6, Sphingomyelin; 7, 3-sulfogalactosylceramide (Sulfatide); 8, Blank; 9, Triglyceride; 10, Diacylglycerol; 11, Phosphatidic acid (PA); 12, Phosphatidylserine (PS); 13, Phosphatidylethanolamine (PE); 14, Phosphatidylcholine; 15, Phosphatidylglycerol (PG); 16, Cardiolipin. (**D**) PtdIns(4)P and (**E**) PtdIns(4,5)P2 concentration in the tiller node of wild type *TabZIP45-4B-WT, TabZIP45-4B* mutant *tabzip45-bb, TabZIP45-4ABD* triple mutant *tabzip45-aabbdd*. Seedlings were grown under 1.0 mM NH_4_NO_3_ (2N) or 0.1 mM NH_4_NO_3_ (0.2N) for two weeks and samples were collected. Data are mean ± S. E. (n ≥ 3). Different letters denote statistically significant differences (P < 0.05) from Fisher’s LSD ANOVA multiple comparisons. **(F)**Seedlings were grown under 2N for two weeks, then were transferred into the 2N test buffer (test buffer with 1 mM NH_4_NO_3_) for 48 hours before Ca^2+^ flux measurement at 100 μm from root tips. **(G)** The seedlings were transferred into a 0.1N test buffer (test buffer with 0.05mM NH_4_NO_3_) and tested immediately. **(H)** After another 24 hours in 0.1N test buffer, the seedlings were tested. **(I)** Seedlings were grown under 0.1N for two weeks, then were transferred into the 0.1N test buffer for 48 hours before Ca^2+^ flux measurement at 100 μm from root tips. **(J, K, L, M)** Statistical comparisons between maximal Ca^2+^ flux amplitude in *TabZIP45-WT* and *tabzip45-bb* of F, G, H, I respectively. Data are mean ± S. E. (n = 6) and *P* values in E-H were from paired Student’s *t*-test. *TabZIP45-WT*, homozygous wild type offsprings. *tabzip45-bb*, homozygous mutant offsprings.

### TabZIP45 triggered PIP2 mediated calcium signaling

Given the role of interaction of TaFTL-43 and TabZIP45-4B in gene expression regulation, we wondered whether TaFTL-43 might be regulated by other molecules. FT proteins were predicted to bind lipids such as phosphatidylethanolamine (*21*, *22*). Previous study revealed that *Arabidopsis* florigen FT bound with phosphatidylcholine (*23*). To identify lipids which TaFTL-43 interacts with, we carried out dot blot assay. TaFTL-43 specifically bound Phosphatidylinositol-4,5-bisphosphate [PtdIns(4,5)P2] and Phosphatidylinositol-3,4,5-trisphosphate [PtdIns(3,4,5)P3] (Fig. 3C). To bolster evidence that the crosstalk between TaFTL-43 and phospholipid was important in nitrogen deficiency adaptation, we analyzed the phospholipid concentration in the shoot basal parts. Compared with the high-nitrogen treatment (2N), The low-nitrogen treatment (0.2N) significantly reduced Phosphatidylinositol-4-phosphate (PtdIns4P) and Phosphatidylinositol-4,5-bisphosphate [PtdIns(4,5)P2] rather than other types of lipids concentrations in *TabZIP45-WT, tabzip45-bb* and *tabzip45-aabbdd* (Fig. 3D and E and fig. S17). These results indicated that either the hydrolysis or synthesis of PtdIns4P and PtdIns(4,5)P2 was disturbed under nitrogen deficient conditions. Furthermore, the PtdIns(4,5)P2 concentration in shoot basal parts of the *tabzip45-bb* plants was higher than that in those of the *TabZIP45-4B* and *tabzip45-aabbdd* plants under 0.2N (Fig. 3E). These results indicated that *TabZIP45-4B* was important in Phosphatidylinositol (PI) metabolism regulation.

PtdIns(4,5)P2 was critical in calcium signaling transduction (*24*). We therefore investigated the calcium signaling under different nitrogen conditions. In normal nitrogen supply (2N), the roots of *TabZIP45-* WT and *tabzip45-bb* plants exhibited a similar level of net Ca^2+^ influx (Fig. 3F and J). We next set up experiment in parallel to access whether the transfer from 2N to deficient nitrogen supply (0.1N) triggered different calcium signaling pattern. Unexpectedly, the calcium influx amplitude between wild type and mutant was not significantly different (Fig. 3G and K), indicating that *TabZIP45-4B* may not be involved in transient response to nitrogen fluctuation. Therefore, we extended the time duration after transfer from 2N to 0.1N for short time (24 hours, ST) and for long time (14 days, LT) in another two independent trails respectively. The significant difference in Ca^2+^ influx between the *TabZIP45-* WT and *tabzip45-bb* plants was observed in LT (Fig. 3I and M), but not in ST (Fig. 3H and L). These data suggested that *TabZIP45-4B* was involved in calcium signaling only during long time nitrogen deficiency response.

In addition to nutrient sensing, calcium signaling plays diverse roles in stress response, and phytohormone crosstalk (*25*). We next characterized whether bZIP45 was responsible for adaptation to other type of environment stresses. Drought conditions limited nitrogen availability. We therefore investigated whether the *tabzip45-bb* was more tolerated to drought stress. As expected, *tabzip45-bb* had more spike number and grain yield under drought conditions (fig. S18). In addition, spike development, as previously mentioned, was closely related to freezing. We therefore determined whether low temperature influenced seedlings survival. Indeed, the survival ratio of *tabzip45-bb* was higher than that of *TabZIP45-4B* and KN199WT under freezing stress conditions, indicating that *TabZIP45-4B* possibly regulated tiller development by freezing resistance (fig. S19). These results indicated that *TabZIP45* regulated tiller development in response to various environmental stresses in form of plants morphogenesis modulation. BRs were closely related to plants morphogenesis regulation and nitrogen response in common wheat and *Arabidopsis* (*26*–*28*) by regulating lipids transfer and calcium signaling (*29*, *30*).

### The mutation of TabZIP45-4B increased grain yield under dense planting by modulating *TaDwarf4*

Environmental stress triggers calcium signaling and limits normal plant growth and development (*25*). We postulated that dense planting will increase grain yield under environmental stresses. We therefore determined whether *TabZIP45-4A, −4B*, and −*4D* expression are response to planting densities. Indeed, we found that *TabZIP45-4B* rather than *TabZIP45-4A* and *TabZIP45-4D* expression was significantly enhanced by the increased planting density (fig. S20). We then evaluated the modulating of *TabZIP45-4B* on yield traits in response to planting densities with or without extra nitrogen fertilizer. The grain yield and spike number of *tabzip45-bb* and *TabZIP45-* WT plants were differentially affected (Fig. 4A to D). The grain yield and spike number were increased in *tabzip45-bb* but slightly affected in *TabZIP45-WT* from D43-D145 under both nitrogen supply levels (Fig. 4A to D). Moreover, The *tabzip45-bb* plants had an advantage over the *TabZIP45-* WT plants in grain yield and spike number under the sowing density of D72.5 (72.5 seeds m^-2^) and D145 (145 seeds m^-2^) with the N270 (270 kg N ha^-1^) treatment (Fig. 4A and D), and under D145 with the N0 (0 kg N ha^-1^) treatment (Fig. 4B and E).

**Fig. 4.**
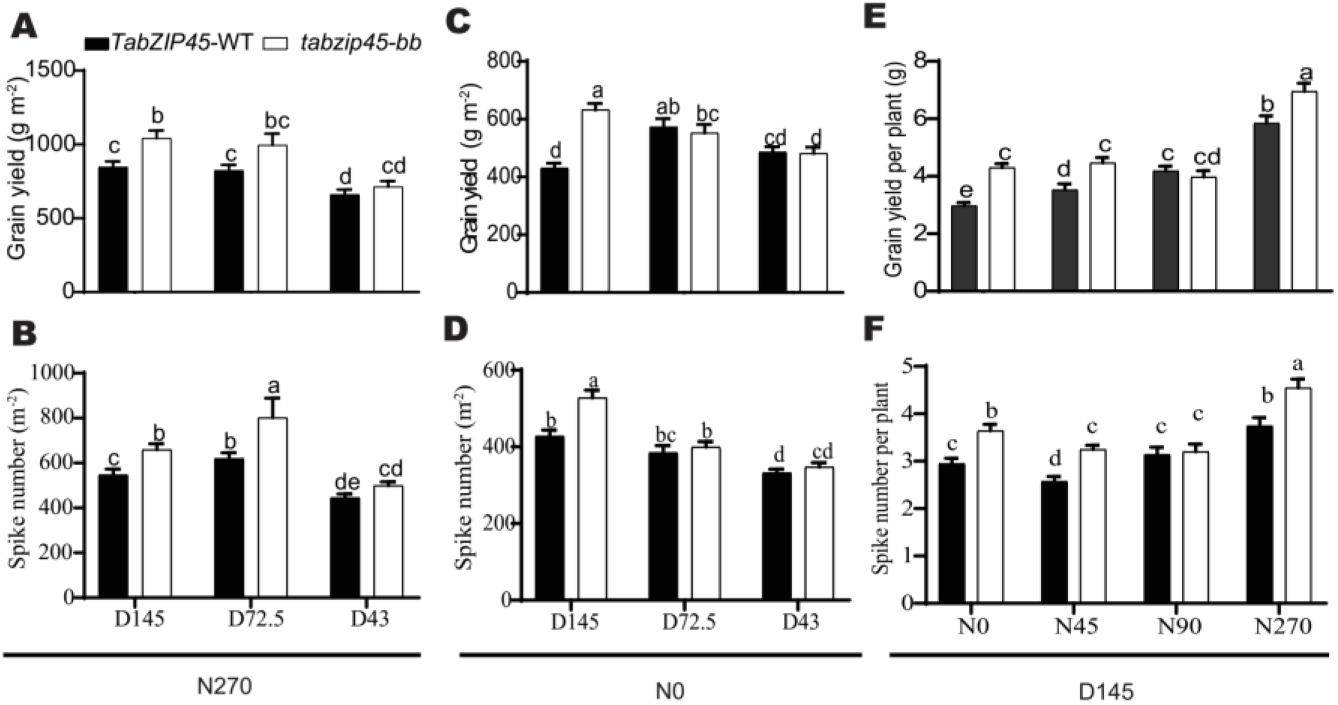
*TabZIP45-4B* edited allele increased grain yield and spike number under dense planting condition. **(A)** Grain yield and **(B)** Spike number under different sowing densities with 270 kg/ha nitrogen supply level. **(C)** Grain yield and **(D)** Spike number under different sowing densities without nitrogen supply. **(E)** Grain yield (per plant) and **(F)** Spike number (per plant) under different sowing densities with different nitrogen supply levels. *TabZIP45-WT*, homozygous wild type offsprings. *tabzip45-bb*, homozygous mutant offsprings. Data are mean ± S. E. of representative plants (n = 40) from more than three independent experiments. Different letters denote statistically significant differences (*P* < 0.05) from two-way ANOVA multiple comparisons. In (A-F), N0 (0 kg N/ha), N45 (45 kg N/ha), N90 (90 kg N/ha), N270 (270 kg N/ha). D43 (43 seeds/m^2^), D72.5 (72.5 seeds/m^2^), D145 (145 seeds/m^2^).

We further accessed the yield advantage of *tabzip45-bb* over *TabZIP45-WT* at the high sowing density (D145) under a wide regimes of nitrogen supply levels, and observed grain yield and spike number advantage at N0, N45 and N270 (Fig. 4E and F). Moreover, the *TabZIP45-* WT plants showed a decline trend in grain yield and spike number when nitrogen supply level was reduced from N270 to N0; however, the *tabzip45-bb* plants maintained a relative stable value of grain yield and spike number when nitrogen supply level was reduced from N90 to N0 (Fig. 4E and F). Furthermore, *tabzip45-bb* had relatively more stable grain number per spike (fig. S21A) particularly at deficient nitrogen supply levels (N0, N45 and N90) than *TabZIP45-* WT. Compared *TabZIP45-* WT, *tabzip45-bb* performed better in biomass and plant height at different nitrogen supply levels, though the decrease of 1000-grain weight in both *TabZIP45-WT* and *tabzip45-bb* (fig. S21B to D). Thus, the decline of spike number and grain yield from N270 to N0 was attenuated by dense planting in *tabzip45-bb* rather than *TabZIP45-* WT.

The rice *OsDwarf4* functions in BRs biosynthesis, and its loss of function mutant was associated with enhanced grain yields under conditions of dense planting and low nitrogen input (*7*, *31*). Furthermore, BR was also associated with root foraging under low nitrogen(*26*). Calcium signaling were essential for both plant response to nitrogen deficient conditions (Fig. 3F-M) and BRs response to environmental stimulations (*30*). We reasoned that *TabZIP45-4B* regulated *TaDwarf4* to adapt to dense planting under different nitrogen conditions in plants. Indeed, we found that TabZIP45-4B could bind to the promoter of *TaDwarf4* (fig. S13B), and knockout of *TabZIP45-4B* reduced *TaDwarf4* expression at 0N (Fig. 2H), suggesting that *TaDwarf4* works downstream of *TabZIP45-4B*.

Low ratio of red light to far red light (R:FR) under dense planting the conditions induced PIP2 hydrolysis to inhibit plants branching (*32*–*35*). Furthermore, the touch among leaves of neighboring plants under dense planting conditions triggered calcium signaling (*36*). *TabZIP45-4B* regulated downstream events by calcium signaling (Fig. 3F-M). We speculated that *TabZIP45-4B* regulated dense planting under nitrogen deficient conditions through *TaDwaf4*-PIP2-Ca^2+^ axis. We therefore mutated *TaDwarf4* by genome editing and obtained triple mutant *TaDwarf4-aabbdd* in three sub-genomes (*TaDwarf4-4A, TaDwarf4-4B* and *TaDwarf4-4D*) to characterize whether *TaDwarf4-aabbdd* response to different planting densities without extra nitrogen supply (N0). Compared with D72.5, relatively denser planting (D145) decreased grain yield of wild type control KN199 (KN199WT), but increased grain yield and spike number of *TaDwarf4-aabbdd* (Fig. 5 A and B) under N0, which was consistent with previous results in rice (*31*). We further characterized whether *TaDwarf4-aabbdd* response to different planting densities under normal nitrogen supply (N270). Similarly, compared with D72.5, D145 increased grain yield and spike number in *TaDwarf4-aabbdd* but only slightly increased grain yield in KN199WT (Fig. 5 A and B). The nearly doubled increase in grain yield and spike number of *TaDwarf4-aabbdd* from D72.5 to D145, which was not observed in KN199WT, indicated that *TaDwarf4-aabbdd* was less sensitive to planting densities than KN199WT possibly owing to BRs mediated PIP2-Ca^2+^ signaling in planting density adaptation was inhibited in *TaDwarf4-aabbdd* (*37*–*39*).

**Fig. 5.**
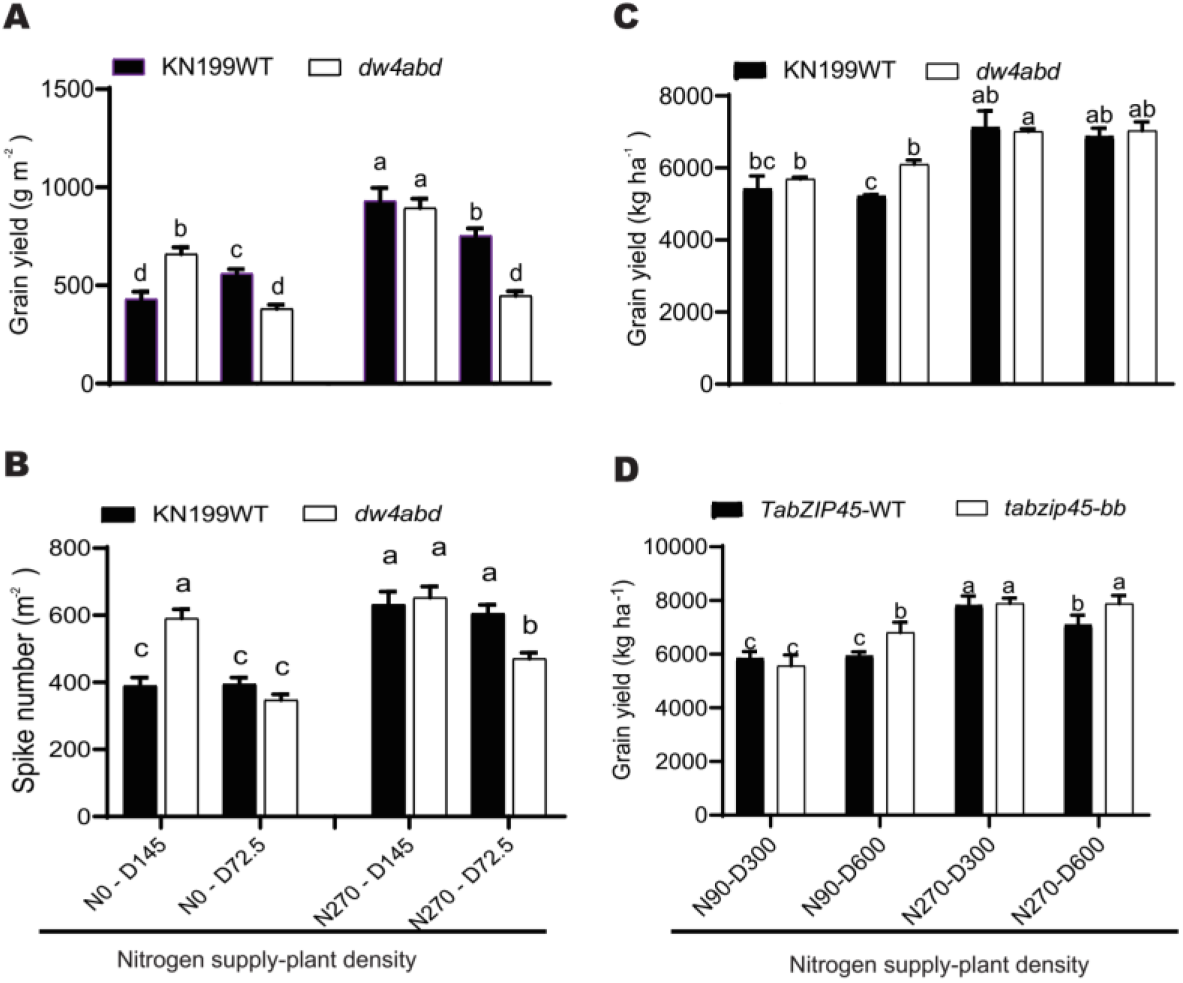
*TabZIP45-4B* rescued nitrogen deficiency induced grain yield loss by regulation of *TaDwarf4*. (**A**) Grain yield. (**B**) Spike number. Data in A-B are mean ± S. E. (n = 40) and different letters denote statistically significant differences (*P* < 0.05) from two-way ANOVA multiple comparisons. KN199WT, KN199 wild type. *dw4-abd*, homozygous mutant offsprings of *TaDwarf-aabbdd*. Nitrogen supply level (270 kg/ha or 0 kg/ha). (**C** and **D**) grain yield (per plot) under different nitrogen supply levels and plant densities. *TabZIP45-WT*, homozygous wild type offsprings. *tabzip45-bb*, homozygous mutant offsprings. Data are mean ± S. E. of representative plants (n = 40) from more than three independent experiments. Different letters denote statistically significant differences (*P* < 0.05) from two-way ANOVA multiple comparisons. In (A-D), N0 (0 kg N/ha), N90 (90 kg N/ha), N270 (270 kg N/ha). D72.5 (72.5 seeds/m^2^), D145 (145 seeds/m^2^), D300 (300 seeds/m^2^), D600 (600 seeds/m^2^).

We then investigated the role of *TaDwarf4* in regulation of the decline of spike number and grain yield from N270 to N0 under different planting densities. Compared with N270, N0 treatment significantly reduced spike number and grain yield of KN199WT regardless of sowing density (Fig. 5 A and B). However, spike number of the *tadwarf4-aabbdd* mutant under D72.5 and D145 only slightly affected by N0 treatment compared with the N270 (Fig. 5B). The results showed that the mutation of *TaDwarf4* inhibited the decrease of grain yield and spike number of KN199WT from N270 to N0 also possibly owing to that BRs mediated PIP2-Ca^2+^ signaling in nitrogen adaptation was blocked in *TaDwarf4-aabbdd* (*37*–*39*).

We further characterized the grain yield of KN199WT, *TaDwarf4-aabbdd*,TabZIP45-4B and *tabzip45-bb* in different nitrogen supply and plant density combinations in a plot experiment. Compared N270 treatment, the N90 treatment reduced grain yield and spike number in both D300 and D600 planting densities in wild type controls of both KN199WT and *TabZIP45-WT* (Fig. 5C and D). Nevertheless, grain yield and spike number loss from N270 to N90 was slightly in *TaDwarf4-aabbdd* and *tabzip45-bb* under D600 though a larger loss under D300 (Fig. 5C and D). Both of the *TaDwarf4-aabbdd* and *tabzip45-bb* mutants had significantly higher grain yield than their corresponding wild type control under D600-N90 (Fig. 5C and D), and the *tabzip45-bb* mutant also had significantly higher grain yield than the *TabZIP45-* WT at D600-N270 (Fig. 5D). Taken together, dense planting (D600) of *tabzip45-bb* mutant reduced nitrogen-deficiency induced grain yield loss by regulating *TaDwarf4*.

## Materials and method

### Pant material and growth condition

The winter wheat varieties Xiaoyan 54 (XY54), Jing 411 (J411), and BC4F5-10 and BC5F1-5 near-isogenic lines of J411//XY54 were planted in the long-term low-nitrogen trial area (2015-2020 growing season) at the Beijing Changping Experimental Base of the Institute of Genetics and Developmental Biology, Chinese Academy of Sciences, and the long-term low-nitrogen trial area (2018-2020 growing season) at the Zhao County Modern Agricultural Park Base in Shijiazhuang, Hebei, China. (2018-2020 growing season). Konong 199 (KN199WT) and each transgenic line: *TabZIP45* knockout lines *tabzip45-dd*, *tabzip45-bb*, *tabzip45-aabbdd*, *TabZIP45-AABBDD*, were grown at the Experimental Station of Cereal and Oil Crop Research Institute of Hebei Provincial Academy of Agricultural and Forestry Sciences, Gao Cheng District, Shijiazhuang, China. The gene editing lines were identified from T0 generation to obtain pure lines with each partial homozygous allele, or pure lines and negative control lines were isolated from heterozygous lines in T0 generation. The tobacco material used for transient expression cotransformation experiments was Nicotiana benthamiana.

### Hydrophobic culture condition

The common wheat culture was carried out as previously described with slight modifications (*40*). Seeds were germinated in 0.05 % hydrogen peroxide overnight. The seeds were spread evenly to germinating in moist condition. Seven-day-old seedlings were transferred to plastic pots containing one-liter nutrient solution (0.2 mM KH_2_PO_4_, 1.5 mM KCl, 1.0 mM MgSO_4_.7H_2_O, 1.5 mM CaCl_2_, 1.0 μM H_3_BO_3_, 0.05μM (NH_4_)_6_Mo_7_O_24_.4H_2_O, 0.05μM CuSO_4_.5H_2_O, 1.0 μM ZnSO_4_.7H_2_O, 1.0 μM MnSO_4_.H_2_O, 0.1 mM FeEDTA(Na), 1.0 mM NH_4_NO_3_). The growth condition was in in 16 hours day - 8 nigh hours cycle at 23°C. The different nitrogen supply levels were 0N (0.0 mM NH4NO_3_), 1N (0.5 mM NH_4_NO_3_), 2N (1.0 mM NH_4_NO_3_), 4N (2.0 mM NH_4_NO_3_). All the nutrition solution were refreshed every 4 days and pH was adjusted to 6.0.

### RNA-seq and gene mapping by sequencing

To isolated genes that were responsible for coordination for growth and environmental response, we constructed a BC5F6 population by crossing between a stressed condition sensitive cultivar Jing411(a traditional Chinese winter wheat cultivar) and stresses tolerant cultivar XiaoYan54 (XY54, a widely used cultivar in breeding). The RNA from three paired-pools (one was with more tiller number and the other was less tiller number) were collected. There paired inbreed line from BC4F5-10 and BC5F1-5 (Line27duo-line27shao, line28duo-line28shao, and 2020duo-2020shao) seedlings at tillering stage were collected and RNA was isolated by GeneMarkbio Plant Total RNA Purification Kit (TR02). The cDNA from reverse transcription was sent to library construction. Hiseq-PE150 was used to sequence each library.

The data from sequenced was filter by Trimmomatic (*41*). The resulting sequence was aligned to common wheat genome reference 2.0 by tophat2 (*42*) (--mate-std-dev 50 -- min-coverage-intron 5 --min-segment-intron 5) and sorted, marked and indexed by samtools (*43*). SNP were called with samtools/bcftools (*44*)pipeline. The SNP were filtered by samtools mpileup (*43*) (-Bugp -R -t DP,AD,ADF,ADR,SP,INFO/AD,INFO/ADF,INFO/ADR -I -q 50 -Q 14 -s) and bcftools and further filter by SWEEP (*45*) when necessary. The SNP depth ratios in different paired pools were calculated according to methods as previously described (*46*). In addition, we confirmed our results by software Triti-Map as previous described (*47*).

### Plasmid construction

Wheat overexpression transgenic vector: *TabZIP45-4A pro::TabZIP45-4A* was constructed on *163-JIT* backbone (kindly donated by Caixia Gao, IGDB). The above constructed vectors were entrusted to our transformation platform to transform the wheat variety KN199WT using the gene gun transformation method as previously described (*48*). Independent transgenic lines were obtained and primers (one end on the vector sequence and the other end on the exogenous sequence) were set at the 5’ and 3’ ends of the respective transgene exogenous sequences, respectively. Positive plants were identified when the size of the sequence fragments amplified at both ends matched that of the plasmid positive control (verified by auxiliary sequencing if necessary) and verified by sequencing. Expression was also identified in field or hydroponic conditions to confirm the success of the respective transgenic events.

Mutant lines were created by gene editing technology (clustered regularly interspaced short palindromic repeats/associated nuclease Cas9, CRISPR/Cas9) (*48*). Selection and design of gene target sites that are conserved in the A, B, and D genomes of each gene and specific to other homologous sequences, and the target site sequences are as follows. *TabZIP45*: TGACAGGTCCGACAGGCCTATGG. *TaDwarf4*: CCTCCTGGCCCTGCTCACCTTC.

The pTaU6-sgRNA plasmids of the respective genes were constructed separately (kindly donated by Caixia Gao). The pTaU6-sgRNA plasmids containing the target sequences and pJIT163-UBI-2NLS-rCas9-wheat (kindly donated by Caixia Gao) were commissioned to the transformation platform of Genova (Tianjin) to transform the wheat variety KN199WT using the gene gun transformation method as described previously (*48*). Transgenic positive plants were obtained and firstly amplified by PCR with primers conserved in the A, B and D genomes of the respective genes. Enzyme digestion was used for identification. In case of positive mutations, the plants were then identified by PCR amplification with primers specific to the A, B and D genomes, enzymatic digestion and sequencing. Primers used were provided in tableS1.

### RNA-seq

The gene editing mutant of TabZIP45-4B tabzip45-bb and wild type KN199WT were used as study materials, and root, leaf, and root-stem joint parts (shoot basal parts) samples were collected after two weeks under 0.2N and 2N culture, respectively. High-quality RNA was extracted by referring to the instructions of GeneMark’s Plant Total RNA Purification Kit (TR02/TR02-150).

The library type was eukaryotic transcriptome class with fragment size between 250-300 bp. The sequencing platform was Hiseq-PE150, and the sequencing data were obtained by removing adapter sequences and low-quality sequences to obtain a data volume of >6 Gb. The data were aligned to the reference genome database using stringtie. All data were homogenized and fragments per kilobase per million mapped reads (FPKM) were calculated by RESM (Li et al., 2011) software. Data with an error rate less than 0.01 and a difference ratio greater than log2 ≥ 2 were used to calculate the differentially expressed genes for comparison. Comparisons were made between KN199WT and the mutant tabzip45-bb in the same treatment. comparisons were made between different treatments of KN199WT or tabzip45-bb.

### ChIP-PCR and ChIP-Seq

The procedures were done as precious described (*40*, *49*). Briefly, seedling under 0N condition for two weeks and whole seedlings was collected and fixed. The fixed samples were powered and lysis. The supernatant from lysis solution after centrifugated and fragmented by ultrasonic crusher was added with antibody to TabZIP45. The beads were collected after washes and then DNA was isolated. The specific primers (supplied in **tableS1**) were used to evaluated DNA enrichment by quantitative PCR.

The library type was eukaryotic ChIP-seq DNA library with a fragment peak size of 100 bp. Sequencing platform was Hiseq-PE150. Sequencing data were removed from splice sequences and low-quality sequences, and the sequence length was >50 bp, resulting in clean data volume >10 Gb. the database was aligned to the reference genome using BWA (*43*) and enriched for target genes. The database was enriched for target genes using MACS (*50*). 63.06% were matched to the reference genome, of which 31.48% were matched to one gene and 21.35% were matched to two genes. Screening conditions: within 2 kb upstream and downstream of the gene, and there was a signal strength difference of 32-fold or more, significant P < 0.00001. primers were designed to verify target site binding ability based on ChIP-seq data.

### Protein prokaryotic purification experiments

*TabZIP45* and *FTL43* were cloned into *pGEX-4T-1* vectors and expressed in *E.coli* BL21(DE3) strain. GST Protein were purified with GE sepharose 4B beads (GE, Boston, MA, USA) according to manufacturer’s instructions.

The full length of *TabZIP45* was clone in to the *Pet28a* vector. The recombinant TabZIP45-HIS tag protein was expressed in *E.coli* BL21(DE3) strain and purified with Ni-NTA magnetic beads (genscript). Primers used were provided in tableS2.

### DNA-binding assays

Assays were done according to methods previously described (*51*). The promoter fragments of *TaGS2, TaNAC2, TaDwarf4, TaCKX2, TaTab1* and *TaLax1* were amplified with biotin tagged primers. The purified TabZIP45-His and biotin tagged promoter fragment were incubated in IP 100 buffer (100 mM potassium glutamate, 50 mM Tris-HCl pH 7.6, 2 mM MgCl_2_, 0.05% Nonidet P40) for 3 hours. Then the streptavidin-bound agarose beads (Sigma) were added and incubated in 4°C for another 4 hours. The resulting solution were set in magnetic stand and washed for three times with IP100 in rotating shaker. Finally, the complex was collected and boiled under 95°C for 10 min and spread on PAGE gel, and detected with anti his antibody (abcam) with ECL SuperSignal West Dura (Thermo Scientific™, USA). Primers used were provided in tableS2.

### Yeast one-hybrid

The *TaGS2, TaNAC2, TaDwarf4, TaCKX2, TaTab1* and *TaLax1* promoter were constructed into the pLacZi plasmid and *TabZIP45* was constructed into the pB42AD plasmid. Both plasmids were simultaneously transferred into yeast (saccharomyces cerevisiae). Apply to plates containing X-gal for color development. Primers used were provided in tableS3.

### GST-pull down assay

Prokaryotic purified proteins bZIP45-4B, 14-3-3 and FTL-43. approximately 10 nmol of magnetic beads-GST-protein or no target protein magnetic beads-GST and 1 nmol of HIS-tagged protein were incubated for 15 min at room temperature and eluted with 1 × phosphate buffer (PBS). The bound proteins were eluted with 20 μl of PBS (50 mM NaCl, 50 mM glutathione and 1 mM DTT, pH 6.8). The eluate was boiled followed by SDS PAGE gel for Western detection. Primers used were provided in tableS2.

### Bimolecular fluorescence complementation (BiFC) assays

CDS sequences of the corresponding genes were amplified by designing adapter primers with attB1 and attB2 (TableS4), and tobacco transient expression vectors were constructed according to the instructions of Gateway®Technology (Invitrogen™, USA). *TabZIP45* and *TaFTL43* was constructed into the vectors for the nitrogen-terminus (N-YFC) and carbon-terminus (C-YFC) of yellow fluorescent protein, respectively. Sequence verified vector was transformed into *Agrobacterium tumefaciens* GV3101, which was injected into tobacco leaves for 36-48 hours. The leaves were cut and assayed under a laser confocal microscope (Zeiss LSM710) for fluorescence signals.

### Subcellular localization

The sequences of the coding regions of *TabZIP45* and *TaFTL43* gene were cloned into *UBI-163-GFP* (kindly presented by Caixia Gao). The fluorescence signal was observed between 12 and 24 h after cotransformation of *TabZIP45-4B-GFP* into wheat protoplasts, and tobacco protoplasts were isolated after 24 h of injection into *Agrobacterium*. The protoplasts were isolated as previously described (28). The protoplasts were observed under a laser confocal microscope (Zeiss LSM710) for fluorescence signals. GFP excitation light range: 500 ~ 550 nm, RFP/mCherry excitation light range: 565 ~ 615 nm. Primers used were provided in tableS5.

### Transcription activation assay

The promoter of *TaTab1* was cloned into *pBD Gal4 F. GUS* was also constructed into *p763-UBI* vector (kindly given by Caixia Gao). The *pBD-TaTab1-Gal4 F* (firefly luciferase), *p163-UBI-GUS* (internal reference 1) and pTRL-Renilla luciferase (internal reference 2) were co-transferred into wheat protoplasts. GUS intensity assay was performed according to the experimental method previous described (*52*). Luciferase fluorescence values are based on the instructions of the Dual-Luciferase®Reporter Assay System kit (Promega, USA). Primers used were provided in tableS6.

### Dot-blot assay

Experiment was done according to methods as previously described (*23*). Berifly, Lipid spotted membrane strips (Echelon Biosciences, USA) was incubated with purified TabZIP45 for four hours. The strip was washed for three times and labeled with antibody of TabZIP45. Finally, the interaction signal between lipid and TabZIP45 was detected by western blot. Primers used were provided in tableS2.

### Western blotting assay

pull off the comb from 10% NuPAGE Bis-Tris gel (Thermo Scientific™, USA) and put the gel into the electrophoresis instrument, add protein electrophoresis buffer to the electrophoresis tank. Set the voltage at 110 V. Stop electrophoresis once the target protein is expected to have been properly separated according to the protein molecular weight, and remove the gel. The membranes were placed sequentially in the pre-chilled transfer solution as blackboard-sponge-filter paper-gel-PVDF membrane-filter paper-sponge-white board (membranes were soaked in methanol for 5 min before use), and the voltage was set at 200 mA, and the membranes were transferred in an ice bath for 90 min. The PVDF membranes were removed and washed five times with TBST (50 mM Tris-HCl (pH 7.5), 8 g/L NaCl, 0.2 g/L KCl, 0.5% Tween-20) five times for 10 min each. Pour off TBST, add primary antibody in the appropriate proportion in TBST containing 0.5% skimmed milk powder and incubate slowly overnight at 4°C or for 2 h at 37°C. Wash the membrane 3 ~ 5 times with TBST for 10 min each time. pour off the TBST, add secondary antibody in 0.5% skimmed milk powder in TBST in appropriate proportions and incubate for 1.5 h at 37°C with slow shaking. The membranes were washed 3 ~ 5 times with TBST for 10 min each time. The membranes were laid flat, ECL reagent was added dropwise to the membranes and photographed with Image Quant LAS 4000.

### quantitative real-time PCR (qRT-PCR)

Total RNAs were isolated from various tissues according to TRIzol™ Plus RNA Purification Kit (Invitrogen™, USA). The first strand cDNA was synthesized according to cDNA RevertAid First Strand cDNA Synthesis Kit (Thermo Scientific™, USA). The obtained cDNA samples were diluted 20-fold and quantitative PCR experiments were performed according to the instructions of the LightCycler®480 SYBR Green I Master (Roche Molecular Systems, Inc., CHE). The specificity of the qPCR reaction was confirmed by confirming that the melting curve of the qPCR was a single peak at the end of the reaction. Calculate the mean Ct value. Determine the relative expression of the target gene based on the expression of the corresponding internal reference gene *TaACTIN* (TraesCS1A02G274400). Primers used were provided in tableS7.

### Phospholipid determination

Materials were collected from wheat seedlings in liquid nitrogen after two weeks of incubation under nitrogen supply level of 1.0 mM NH_4_NO_3_ or low nitrogen supply level of 0.1 mM NH4NO3 in the greenhouse. Tiller nodes of 100 ± 5 mg were isolated under dry ice and transferred to the Lipidome Platform of the Institute of Genetics and Developmental Biology, Chinese Academy of Sciences for determination of phospholipids concentrations. The determine of phospholipids concentration was done as previously described (*53*).

#### SNPs in different haplotypes determination assay

The different haplotypes of *TabZIP45-4B* seedlings were under 0N (0.0 mM NH_4_NO_3_) and 2N (1.0 mM NH_4_NO_3_) conditions for two weeks. The isolated DNA and RNA from these whole seedlings were used to determined SNPs and splicing variant. Related primers were listed in **table S5.**

#### Screen for developments regulators assay

The different haplotypes of *TabZIP45-4B* seedlings at different developmental stages from field experiment sowed at autumn grew under natural conditions were collected. The RNAs from these samples were isolated and the different genes expression at transcriptional level were determined by qRT-PCR. Primers used were provided in tableS7.

### Evolutionary tree building

The homologous protein sequences of TabZIP45 were downloaded from the database (http://plants.ensembl.org and https://www.rcsb.org). Evolutionary distances were calculated and evolutionary trees were constructed using MEGA 6.0 software for neighbor-joining.

### Statistical analysis

All data were counted and plotted using GraphPad Prism 8 or Microsoft office 2016. p-values less than 0.05 were considered to be significantly different. Appropriate statistics were selected according to the significance of the orthogonal distribution of the data and the conditions of processing.

## Supporting information

All suplementary

## ACKNOWLEDGMENTS

We thank Prof. Caixia Gao’s laboratory (Institute of Genetics and Developmental Biology, Chinese Academy of Sciences) for developing the transgenic wheat lines. We thank Prof. Guanghou Shui’s laboratory (Institute of Genetics and Developmental Biology, Chinese Academy of Sciences) for performing lipids determination and analysis.

## Funding

This research was supported by the Strategic Priority Research Program of Chinese Academy of Sciences, Grant No. XDA24010202

## Author contributions

Z.H. carried out most of experiments. Y.T. initiated and supervised the project. Z.H. conceived the project. Z.H. and Y.T. designed the experiments. Z.H. performed field experiments. Z.H. and M.H. collected data for traits. Z.H. performed in cellular, molecular and biochemistry experiments. X.H. provided technique support in molecular and biochemistry experiments. X.Z., W.T., M.H. and H.L. provided technique support in agriculture experiments. Z.H. and Y.Z. analyzed the data and computation. Z.H. wrote the manuscript with input of all other authors. Y.T. revised the manuscript.

## Competing interests

all authors declare no competing interests.

## Data and materials availability

all data and material are available on line. Requests for materials should be sent to Y.T.

## List of supplementary materials

Figs. S1 to S22

Tables S1 to S7

---

**Fig.S1.**
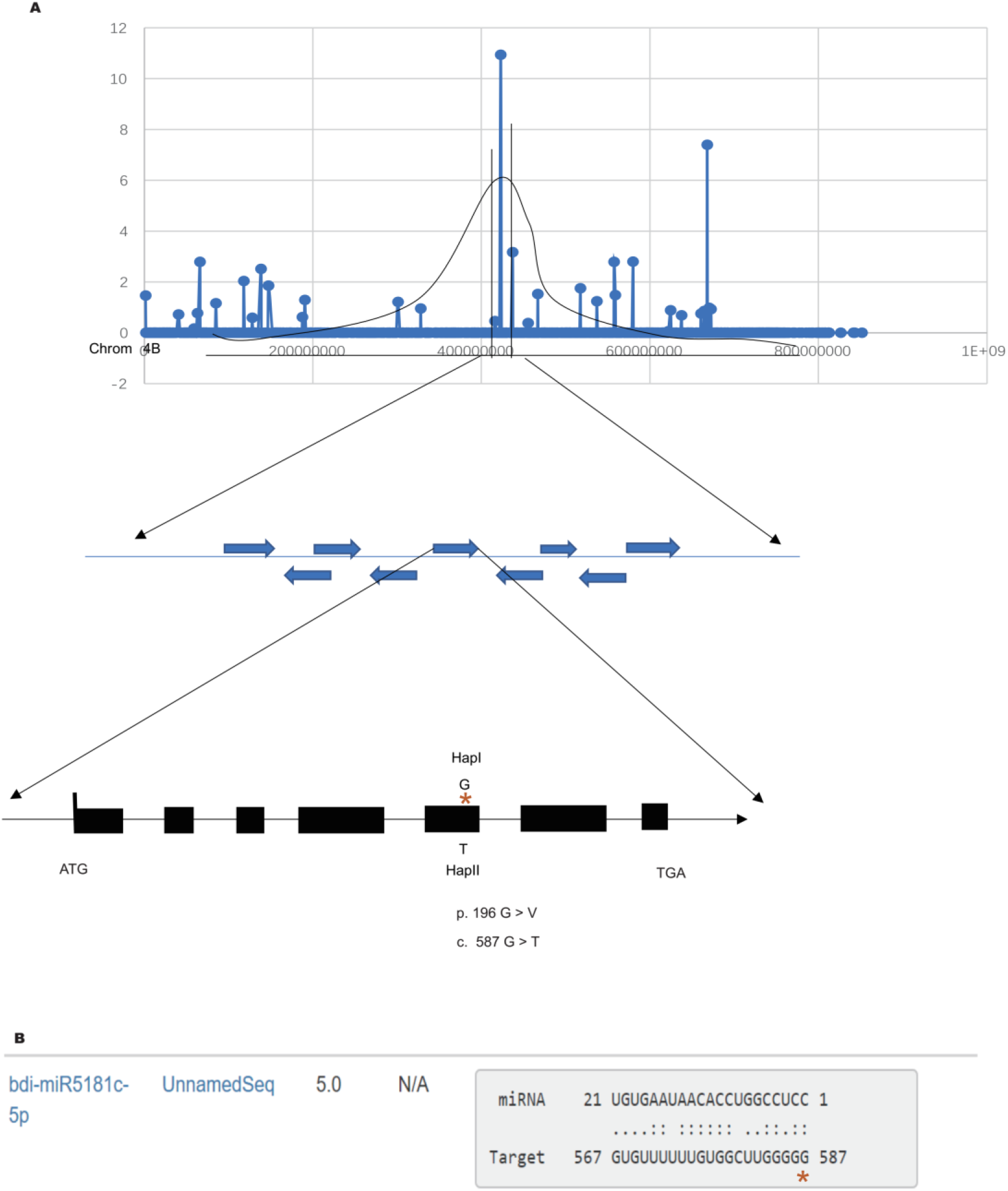
*TabZIP45* isolation and miRNA targeted to *TabZIP45-4B*. (**A**) the isolation of *TabZIP45-4B* by mapping of sequencing. (**B**) the mutation site is predicted to disrupt the binding sites of microRNA (http://plantgrn.noble.org/psRNATarget/).

**Fig.S2.**
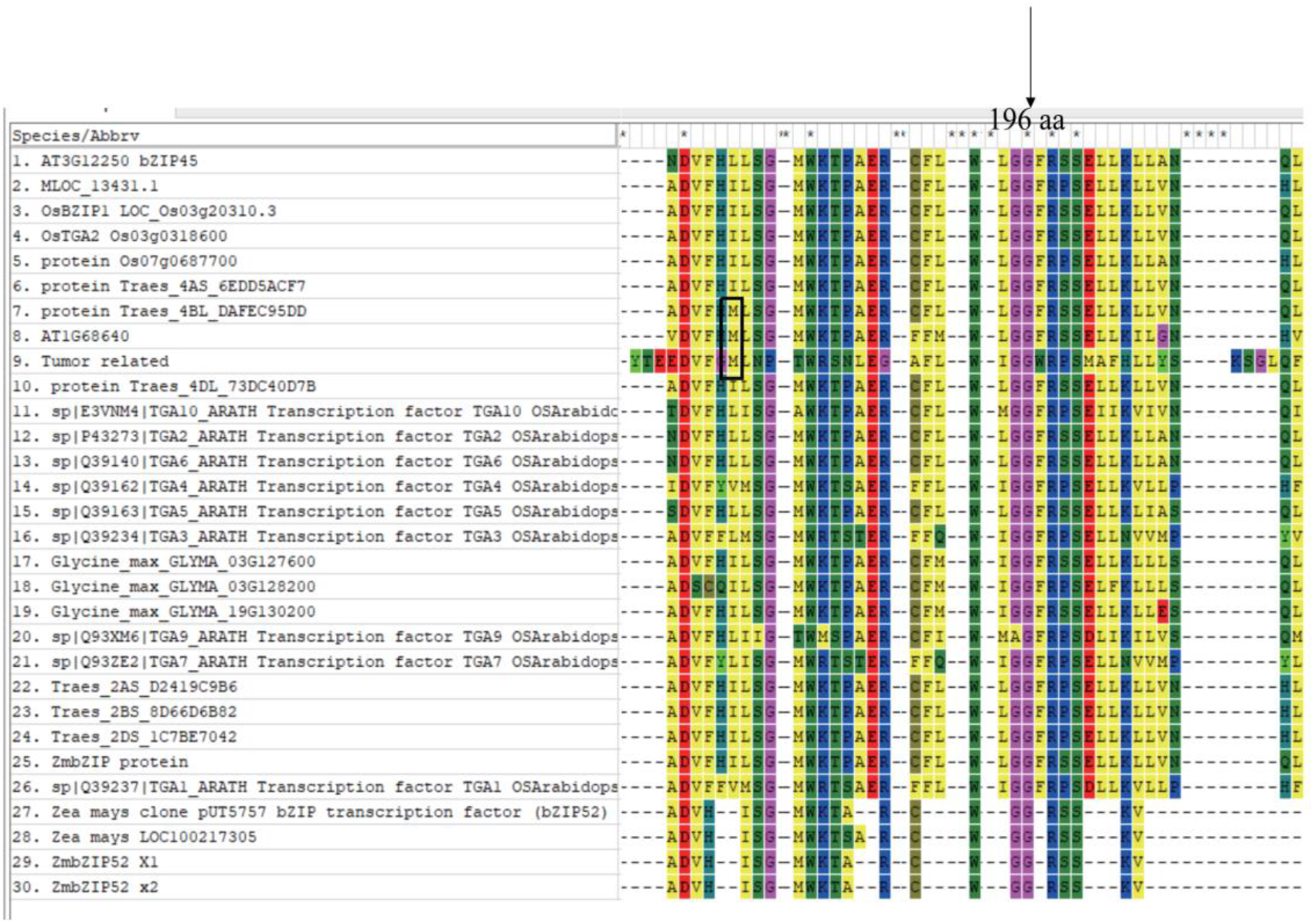
The TabZIP45 sequence conservation. 196 aa in DOG domain of TabZIP45-4B protein is conserved in angiosperms. The right above arrow indicates the conserved 196 aa (Glysine, G) in DOG domain of bZIP45 proteins in plants. The left above arrow indicates 178 aa difference among TabZIP45-4A, −4B and −4D. The Arrow pointed at dashed line box at middle left side of panel indicates the position of Methioine. The alignment is constructed by MEGA 6.0 by ClustalW method. Pairwise alignment and multiple alignment penalty is 8. Gap extension penalty is 0.025. Protein Weight Matrix is identity. Residue-specific penalties and hydrophilic penalties is on. Gap separation distance is 5. End Gap Separation is on. Use Negative Matrix, delay divergent cutoff is 15%. Keep predefined gaps.

**Fig.S3.**
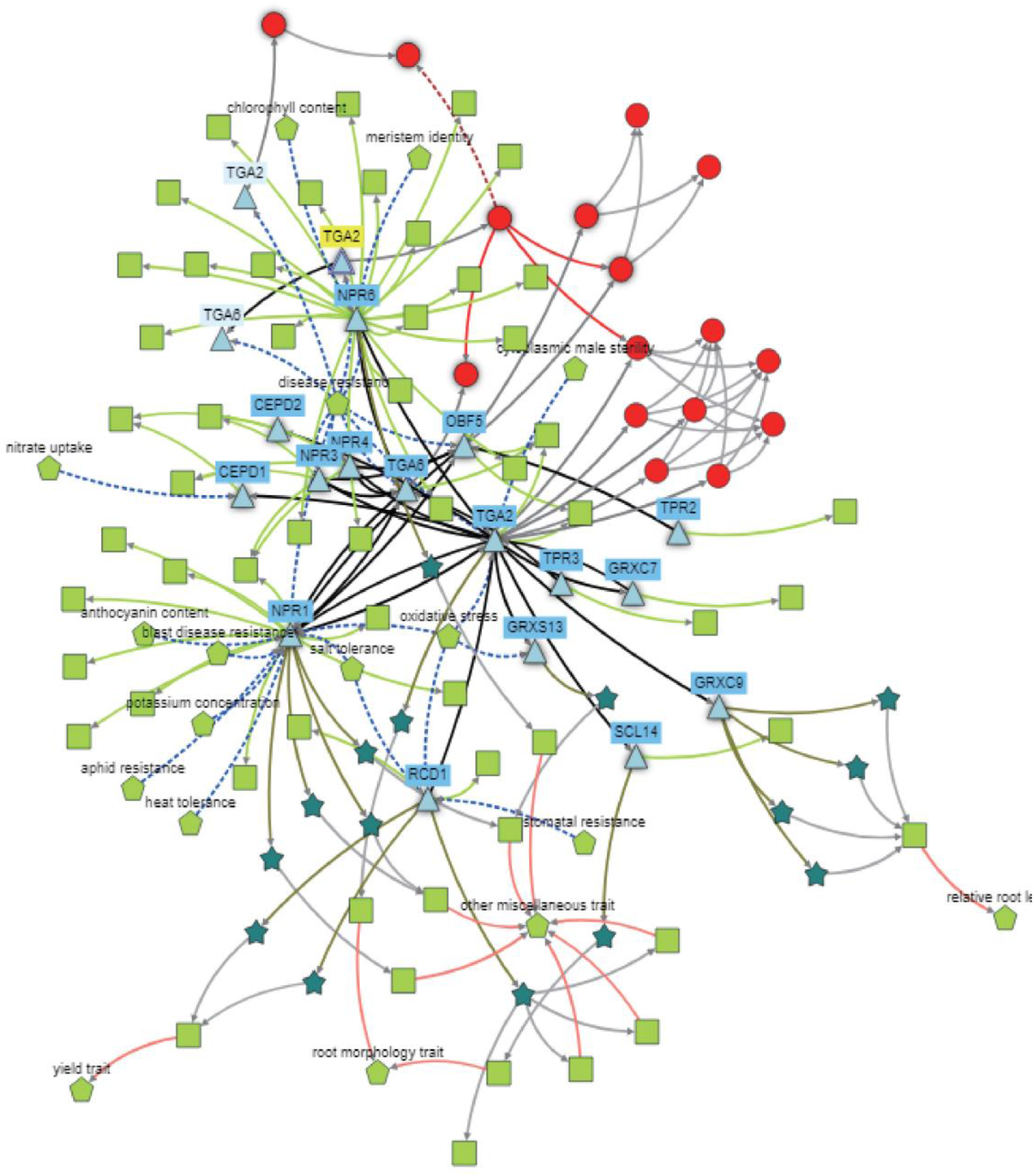
*TabZIP45* genetic network. The network of *TabZIP45-4B* in nutrients sensation, growth regulation and stresses adaptation. The network was constructed in knetminer (https://knetminer.com/Triticum_aestivum/).

**Fig.S4.**
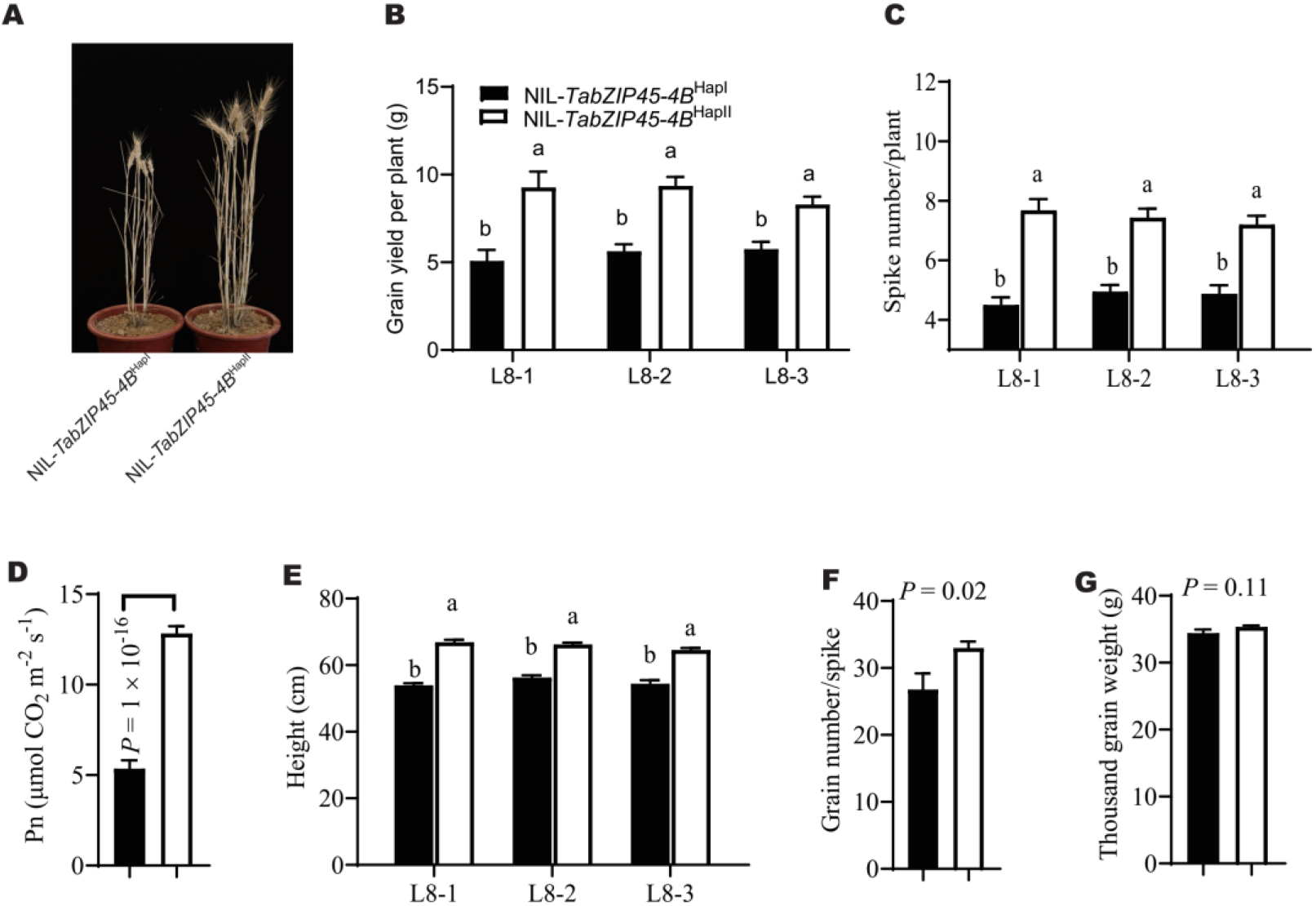
Agronomic traits of NIL-*TabZIP45-4B^HapI^* and NIL-*TabZIP45-4B^HapII^* under nitrogen deficient conditions. **(A)** Representative of more than four independent replicates. Scale bar = 10 cm. **(B)** Grain yield per plant. **(C)** Spike number per plant. **(D)** photosynthetic rates. (**E**) Height. (**F**) Grain number per spike. **(G)** 1000-grain weight. Data in B-G are mean ± S. E. (n = 40) and different letters denote statistically significant differences (*P* < 0.05) from two-way ANOVA multiple comparisons. L8-1, L8-2 and L8-3: different isogenic lines with NIL-*TabZIP45-4B*^HapII^or *NIL-TabZIP45-4B*^HapI^. Nitrogen supply level: 30 kg N/ha. Isogenic lines *NTL-TabZIP45-4B^HapI^* and *NIL-TabZIP45-4B^HapII^* are generated by introducing *TabZIP45-4B^HapI^* from J411 into XY54 (BC5F5).

**Fig.S5.**
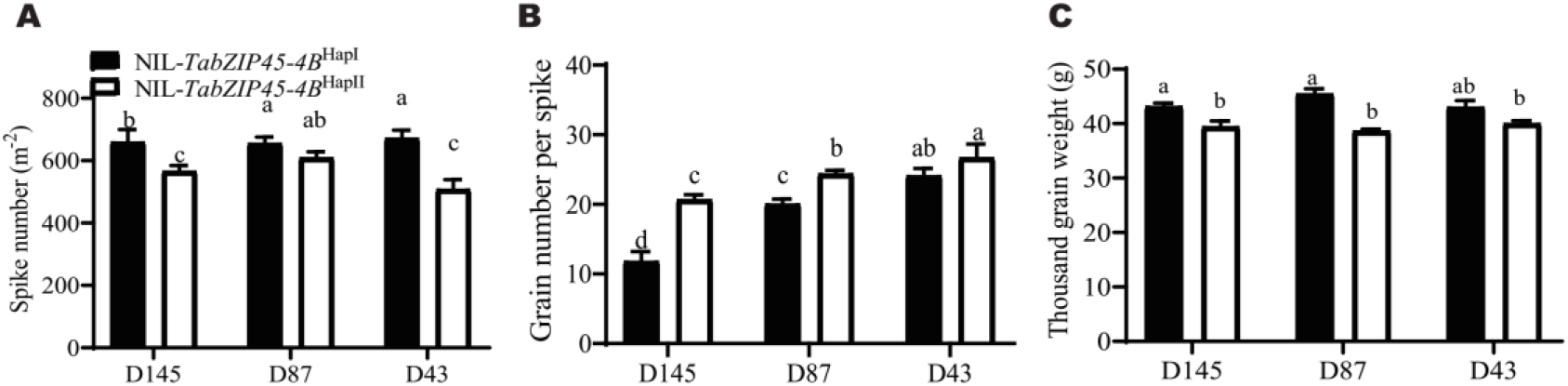
Agronomic traits of NIL-*TabZIP45-4B^HapI^* and NIL-*TabZIP45-4B*^HapII^ under different planting densities. (**A**) Spike number per plant (**B**) Grain number per spike and (**C**) Thousand grain weight under different nitrogen supply levels and sowing densities. Data in A and B are mean ± S. E. (n ≥ 34) Isogenic lines NIL-*TabIP45-4B*^HapI^ and NIL-*TabZIP45-4B*^HapII^ are generated by introducing NIL-*TabZIP45-4B*^HapII^ from J411 into XY54 (BC5F6). In (A-C), D43 (43 seeds/m^2^), D87 (87 seeds/m^2^), D145 (145 seeds/m^2^).

**Fig.S6.**
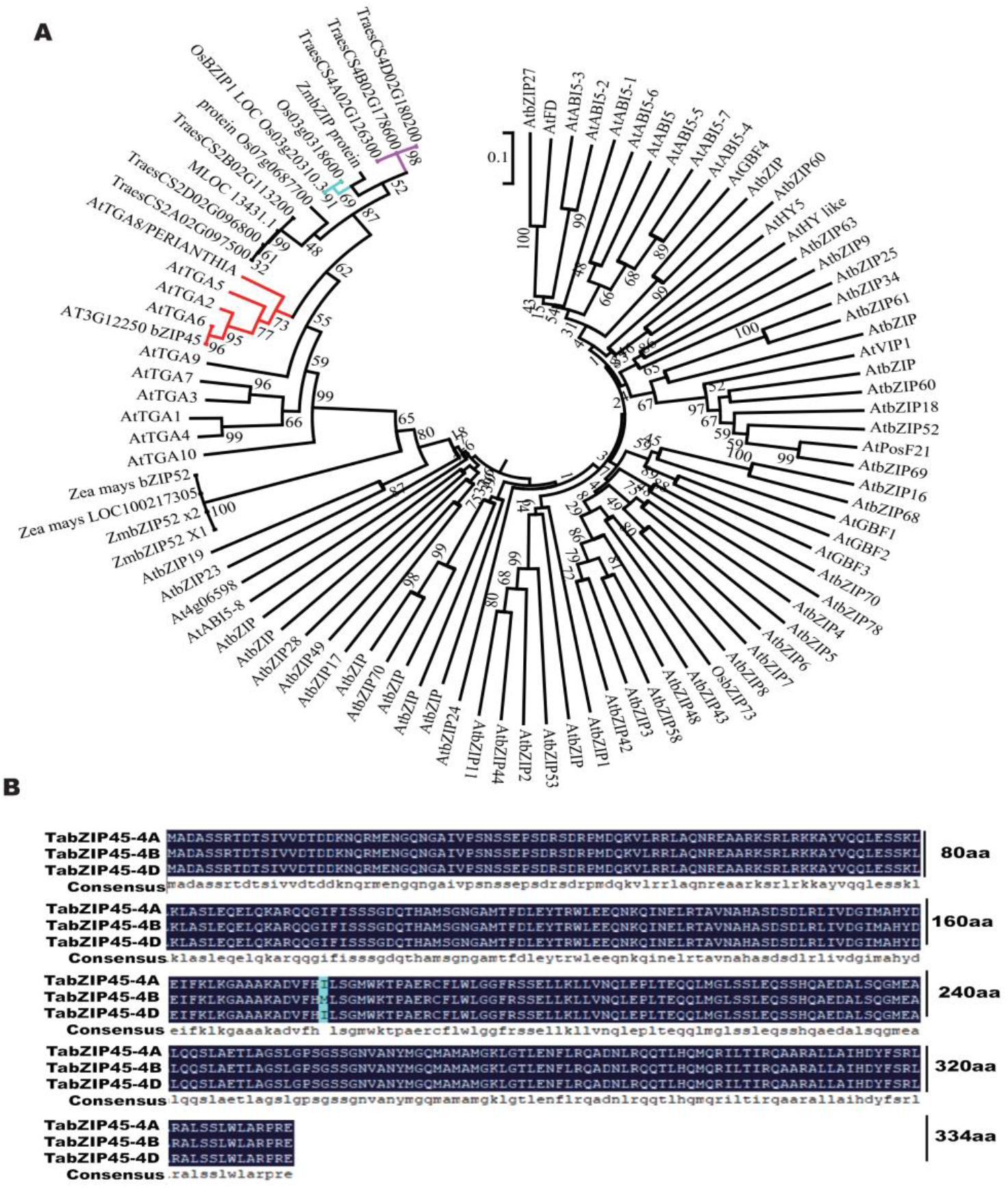
TabZIP45 phylogeny tree and sequence. (**A**) The phylogeny tree is draw by neighbor-joining method of MEGA 6.0. (**B**) Coding sequences of *TabZIP45-4A, −4B* and −*4D* cloned in KN199, which was generated by DNAMAN (LynnonBiosoft, USA).

**Fig.S7.**
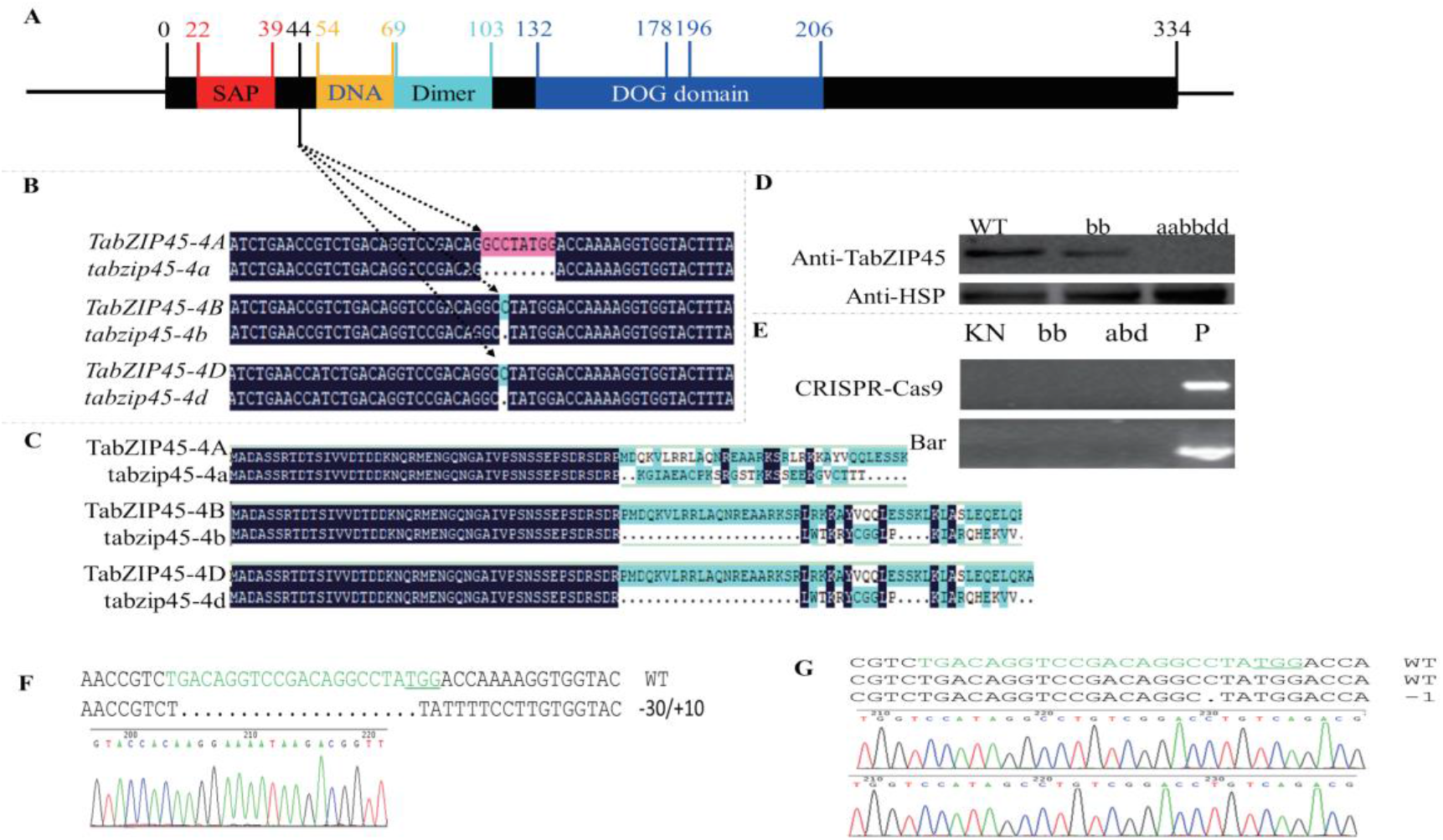
TabZIP45 protein structure domain and *TabZIP45* gene editing sites. (**A**) Structure domain of TabZIP45, number stands for aa sites. Red region, Ser-Ala-Pro (SAP) motif. Yellow region, DNA binding sites domain of TabZIP45. Cyan region, Dimmer interface domain. Blue region, Delay of Germination (DOG) domain. Original amino acids site (aa) in 178 between TabZIP45-4A, 4D and 4B is different. Two alleles of different haplotypes encode different aa in 196. (**B**) Genome editing leads to 8 bp insertion in *TabZIP45-4A*, one bp deletion in *TabZIP45-4B* and *TabZIP45-4D* respectively. (**C**) Premature stop in *TabZIP45-4A, 4B and4D* caused by genome editing produce 73 aa, 66 aa and 66 aa mutant protein respectively, without original structure domain. (**D**) The mutants are confirmed by TabZIP45 antibody. (**E**) Determine whether the plants have CRISPR-Ca9 and anti-Basta (Bar) genes. In (D) and (E), WT, homozygous wild type offsprings separated from last generation; *bb*, homozygous mutant offsprings separated from last generation; *aabbdd, TabZIP45-4A, 4B* and *4D* all homozygous mutant; P, plasmid positive control. (**F** and **G**) the sequence results of two independent *TabZIP45-4B* mutants.

**Fig.S8.**
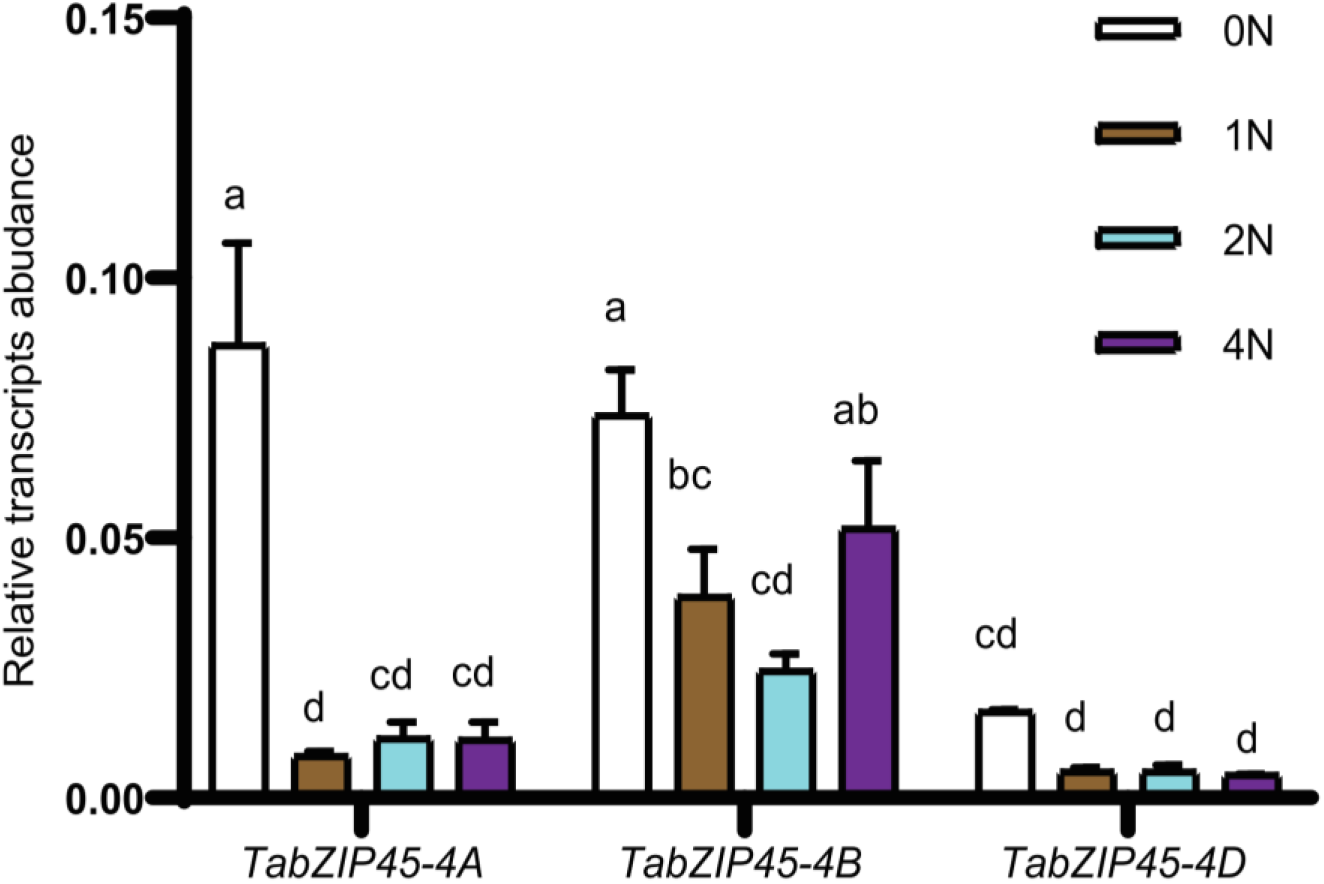
The response of *TabZIP45* to nitrogen supply. The seedlings of KN199WT were grown for 4 weeks in the nutrient solutions that contained 0.0 mM NH_4_NO_3_ (0N), 0.5 NH_4_NO_3_ (1N), 1.0 NH_4_NO_3_ (2N) and 2.0 NH_4_NO_3_ (4N). The shoots were collected for gene expression analysis. Data are mean ± S. E. (n = 4). Different letters denote statistically significant differences (*P* < 0.05) from Fisher’s LSD ANOVA multiple comparisons. The genes expression level was normalized to that of *Taactin* as internal control.

**Fig.S9.**
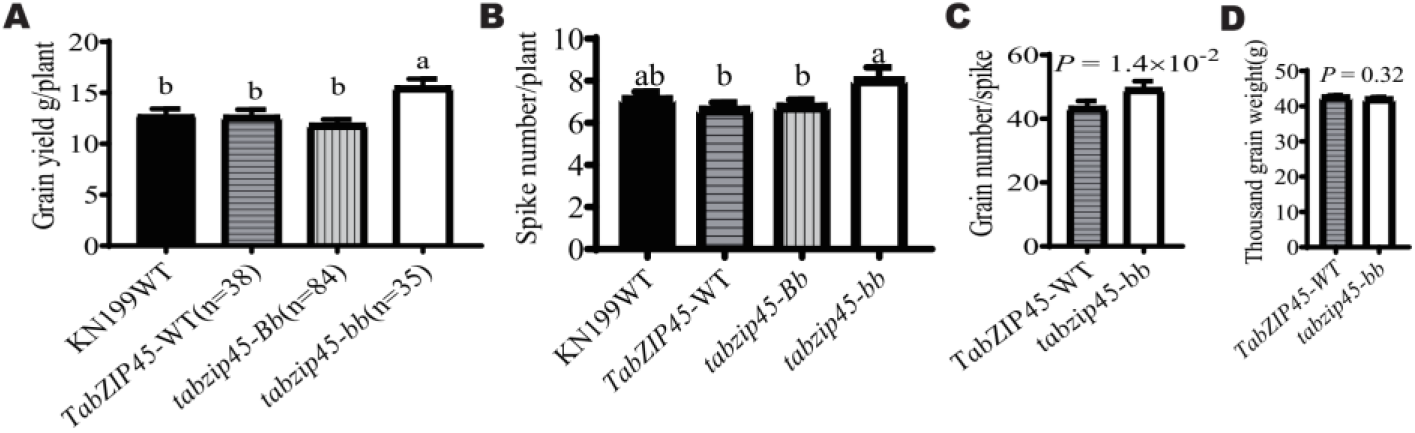
Knockout of *TabZIP45-4B* increased grain yield. **(A)** Grain yield per plant. **(B)** Spike number per plant. Data in A and B are mean ± S. E. (n ≥ 35). Different letters in A-B denote statistically significant differences (*P* < 0.05) from Fisher’s LSD ANOVA multiple comparisons. **(C)** Grain number per spike. *P* value in C is from a Mann-Whitney *t* test. **(D)**1000-grain weight. *P* value in C-D is from a two tailed unpaired test. In A D: KN199WT, KN199 wild type; *TabZIP45-WT*, homozygous wild type offsprings separated from last generation; *tabzip45-Bb*, heterozygous mutant offsprings separated from last generation; *tabzip45-bb*, homozygous mutant offsprings separated from last generation. Nitrogen supply level (270 kg N/ha).

**Fig.S10.**
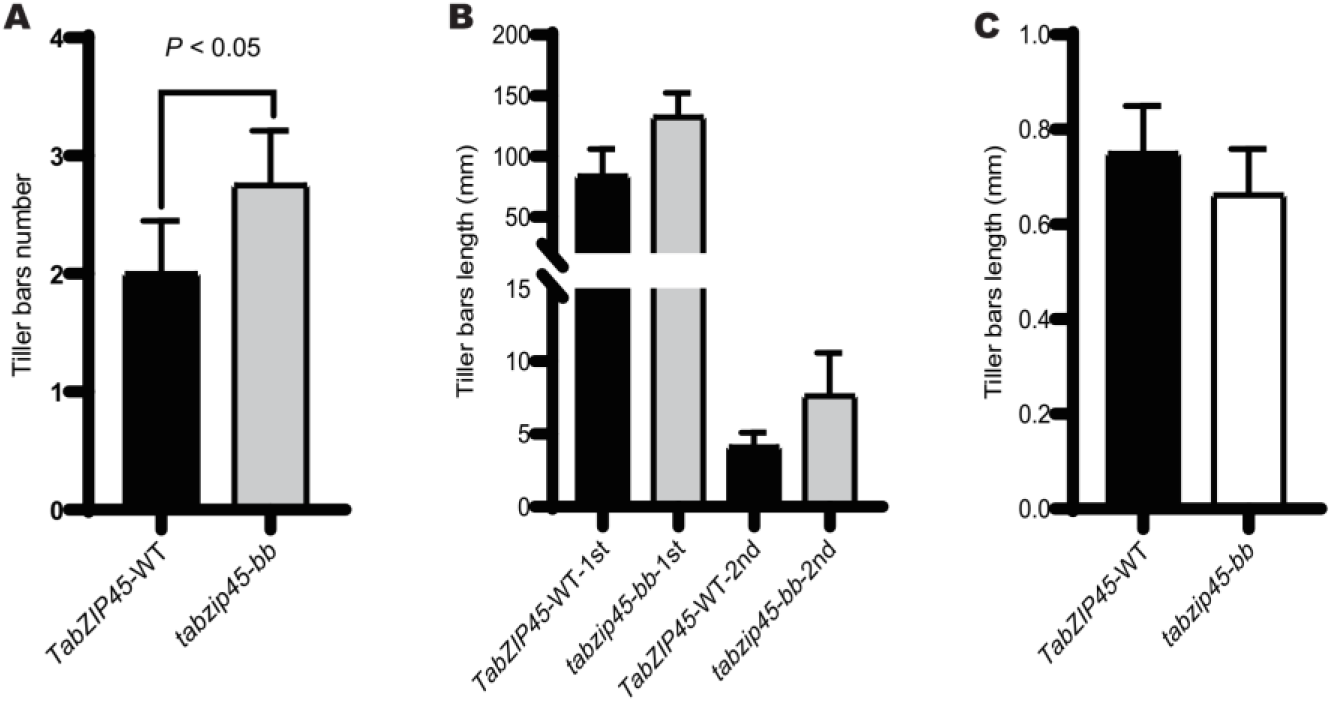
*TabZIP45-4B* inhibited tiller initiation and tiller elongation. (**A**) Tiller buds number in 2N. (**B**) Tiller buds length in 2N. (**C**) Tiller buds number in 0.2N. Seedlings were grown under high N conditions (1.0 mM NH_4_NO_3_, 2N) or (0.1 mM NH_4_NO_3_, 0.2N) for two weeks and samples were collected. Data are mean ± S. E. (n = 8) and *P* values in A-C were from paired Student’s *t*-test. 1st is first tiller, 2nd is second tiller. *TabZIP45-WT*,homozygous wild type offsprings. *tabzip45-bb*, homozygous mutant offsprings.

**Fig.S11.**
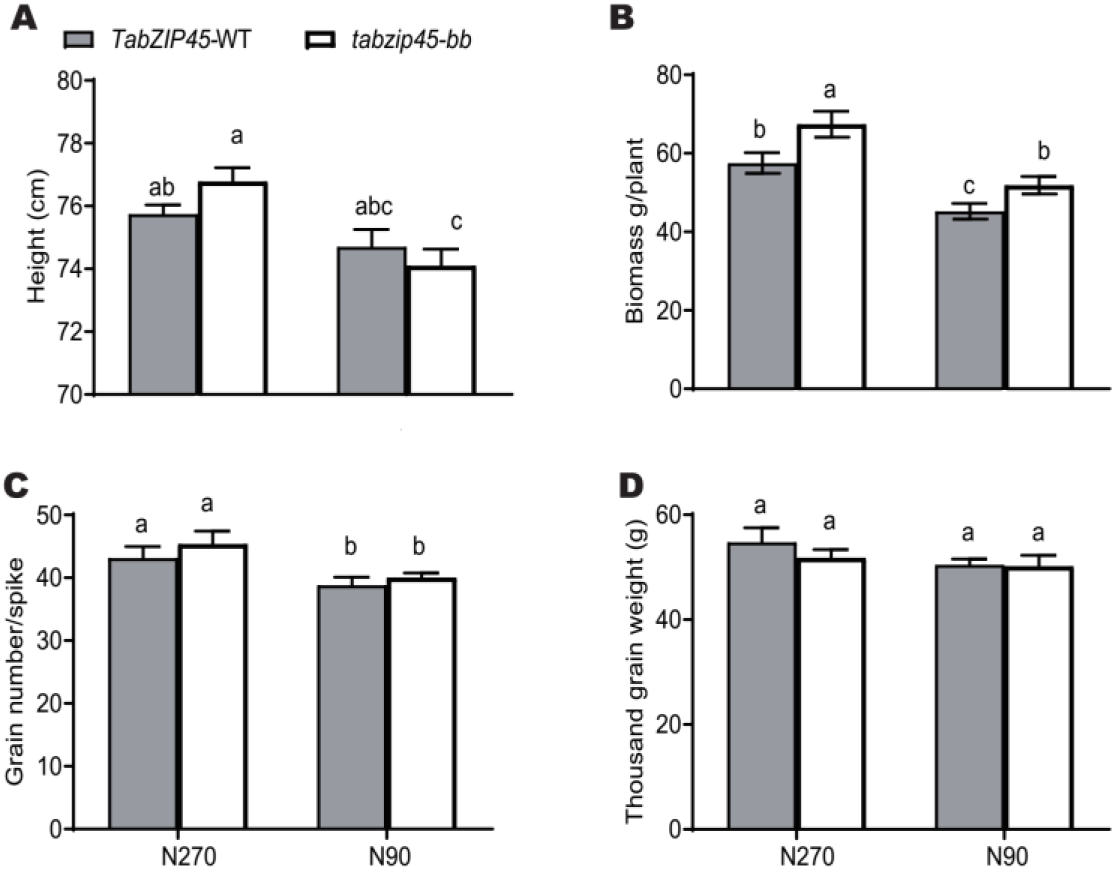
Knockout of *TabZIP45-4B* increased grain yield. **(A)** Height. **(B)** Biomass per plant. **(C)** Grain yield per plant. (**D**) Grain number per spike. (**E**) 1000-grain weight. Data in A-E are mean ± S. E. (n = 40) and different letters denote statistically significant differences (*P* < 0.05) from two-way ANOVA multiple comparisons. *TabZIP45-WT*, homozygous wild type offsprings. *tabzip45-bb*, homozygous mutant offsprings. In (A-D), (90 kg N/ha), N270 (270 kg N/ha).

**Fig.S12.**
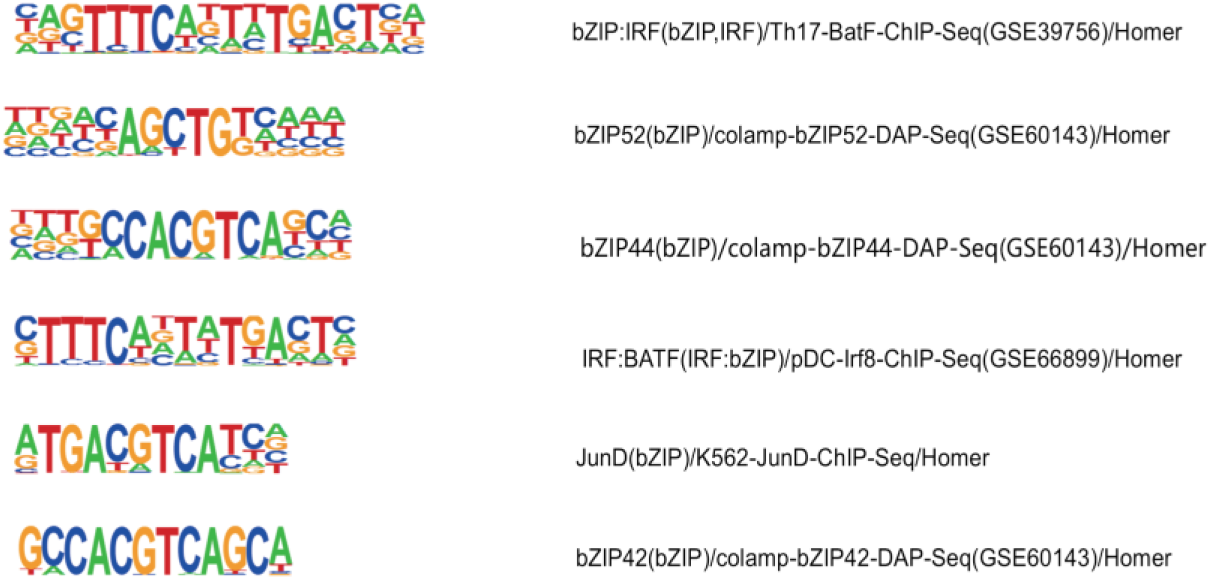
TabZIP45 binding sequence motif. The sequence motif of bZIP family enriched by ChIP-seq.

**Fig.S13.**
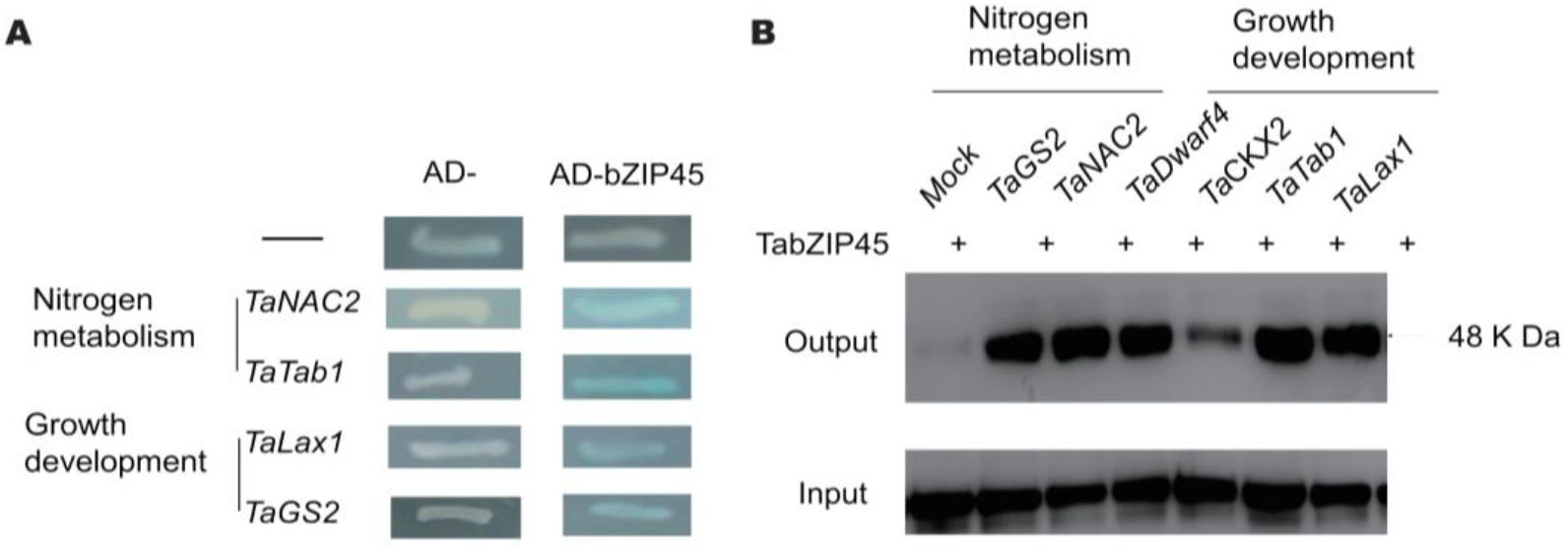
The binding ability of TabZIP45-4B to the promoters of downstream genes. **(A)** Yeast one hybrid using TabZIP45-4B and the promoters of candidate downstream genes; **(B)** Prokaryotic purified TabZIP45-4B was incubated with different genes’ promoters with biotin marked at 5’end of both DNA strands and probed with biotin antibody.

**Fig.S14.**
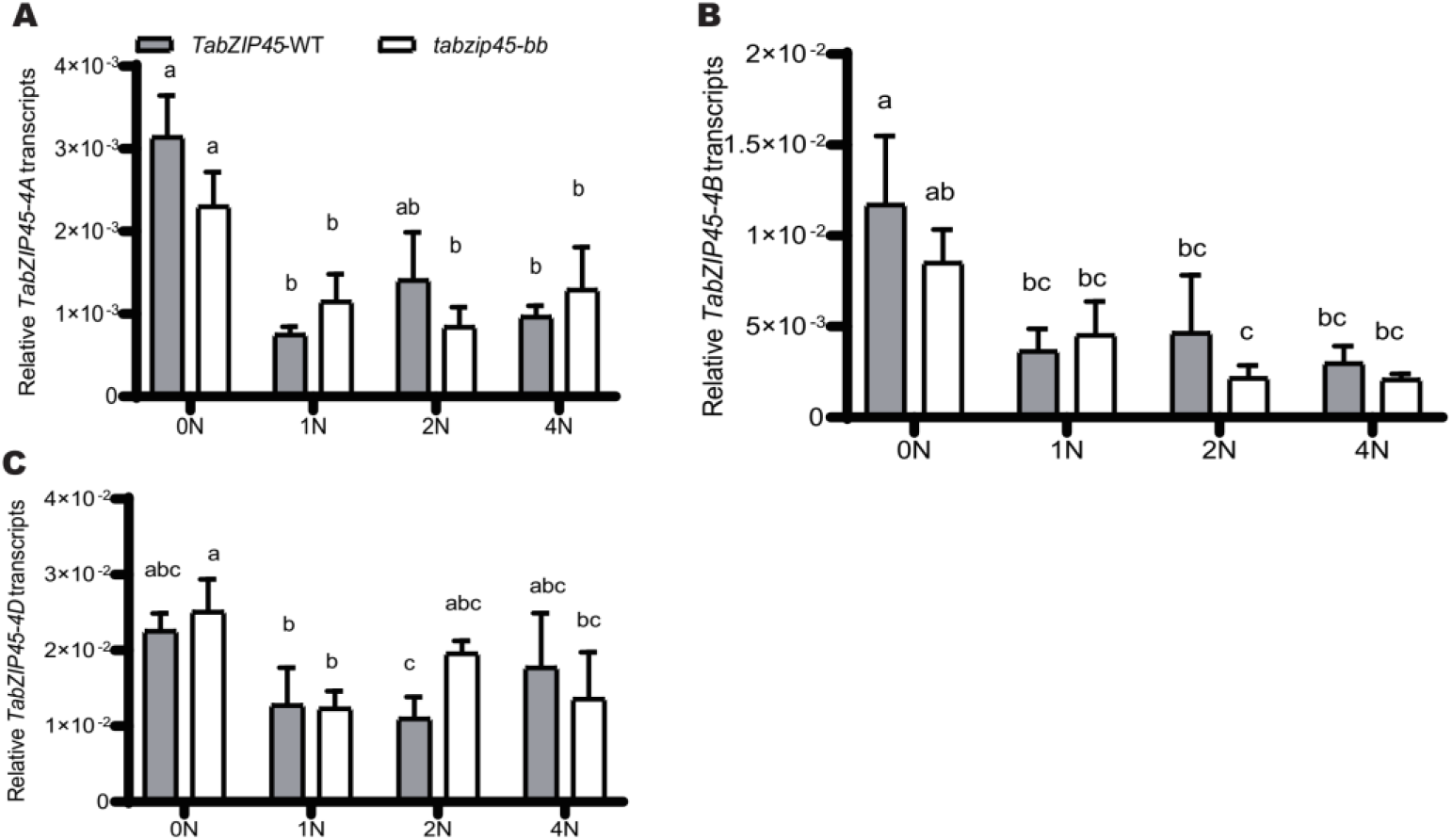
Knockout of *TabZIP45-4B* did not affect *TabZIP45-4A, −4B, −4D* expression. (**A**) *TabZIP45-4A*, (**B**) *TabZIP45-4B*, (**C**) *TabZIP45-4C*. The seedlings of *TabZIP45-4B-WT* and *tabzip45-bb* mutant were grown for four weeks in the nutrient solutions which contain 0.00 mM NH_4_NO_3_ (0N), 0.50 mM NH_4_NO_3_ (1N), 1.00 mM NH_4_NO_3_ (2N) and 2.00 mM NH_4_NO_3_ (4N). Then the shoots and roots were collected for gene expression analysis. *TabZIP45-WT*, homozygous wild type offsprings. *tabzip45-bb*,homozygous mutant offsprings. Data in A-D were from shoots, and data in E and F were from roots. The genes expression level was normalized to that of *Tatublin* as internal control. Data are mean ± S. E. (n ≥ 3). Different letters denote statistically significant differences (*P* < 0.05) from Fisher’s LSD ANOVA multiple comparisons.

**Fig.S15.**
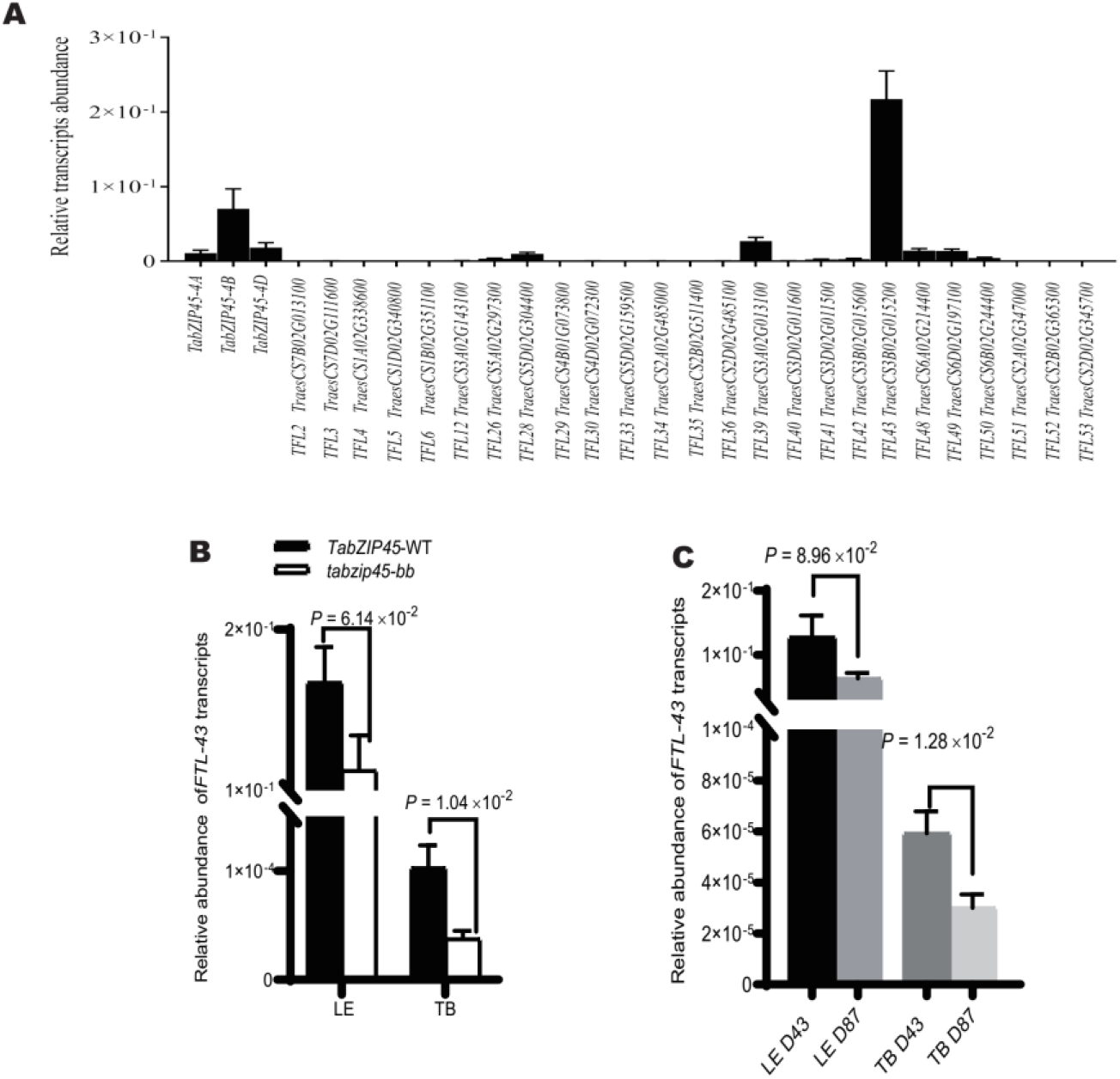
*TaFTL43* was essential in wheat development and interaction with environment. **(A**) Relative expression of *TaFTL* genes in above ground parts collected 27 days after sowing in Hebei province under 45 kg N/ha supply. **(B-C)** Relative expression of *TaFTL43* (in leaves (LE) and tiller basal parts (TB) collected 27 days after sowing. The genes expression level was normalized to that of *Tatublin* as internal control. Data are mean ± S. E. (n = 4). *P* values in B-C are from a two tailed unpaired Mann-Whitney or Welch’s *t* test depends on data normality significance. In (D), D43 (43 seeds/m^2^), D87 (87seeds/m^2^).

**Fig.S16.**
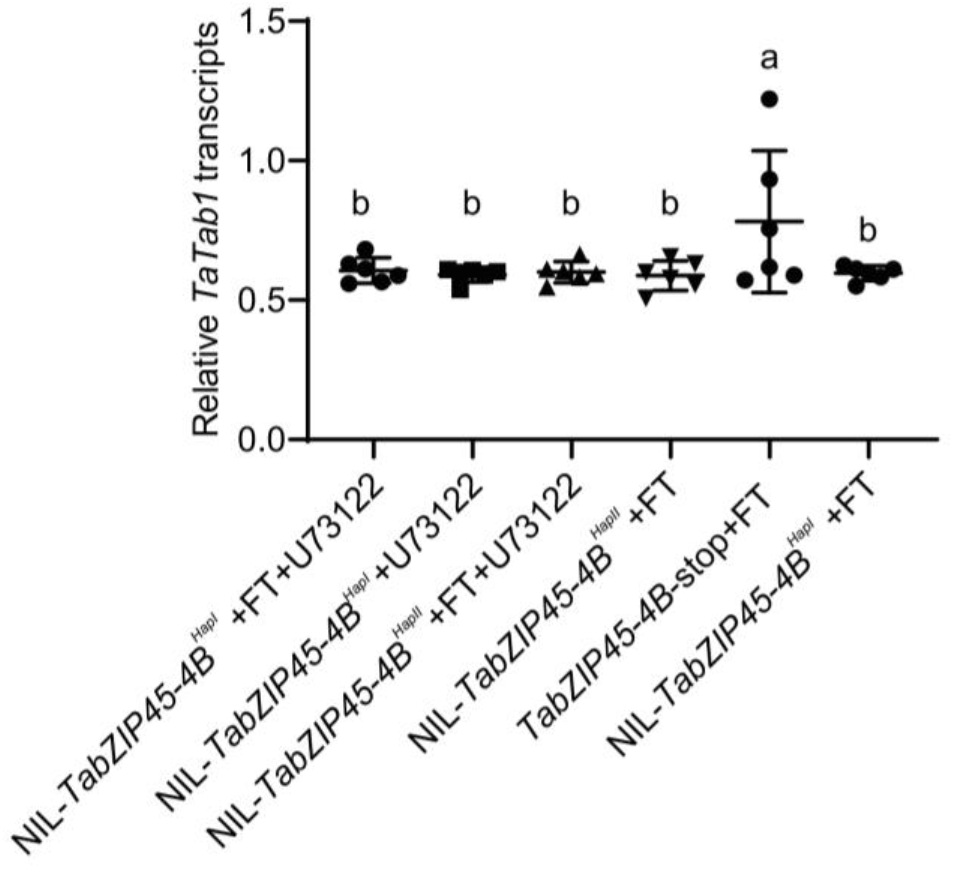
*TabZIP45-4B* inhibited *Tatab1* expression at presence of TaFTL-43. The knock out *TabZIP45-4B* promoted *Tatab1* promoter transcription. The different haplotype of *TabZIP45-4B* did not affect transcriptional activation on *Tatab1*. FT, TaFTL-43; U73122, Phospholipase C (PLC) inhibitor. Relative luciferase was normalized to internal luciferase reference. Data were presented as mean ± S.E. (n=6)

**Fig.S17.**
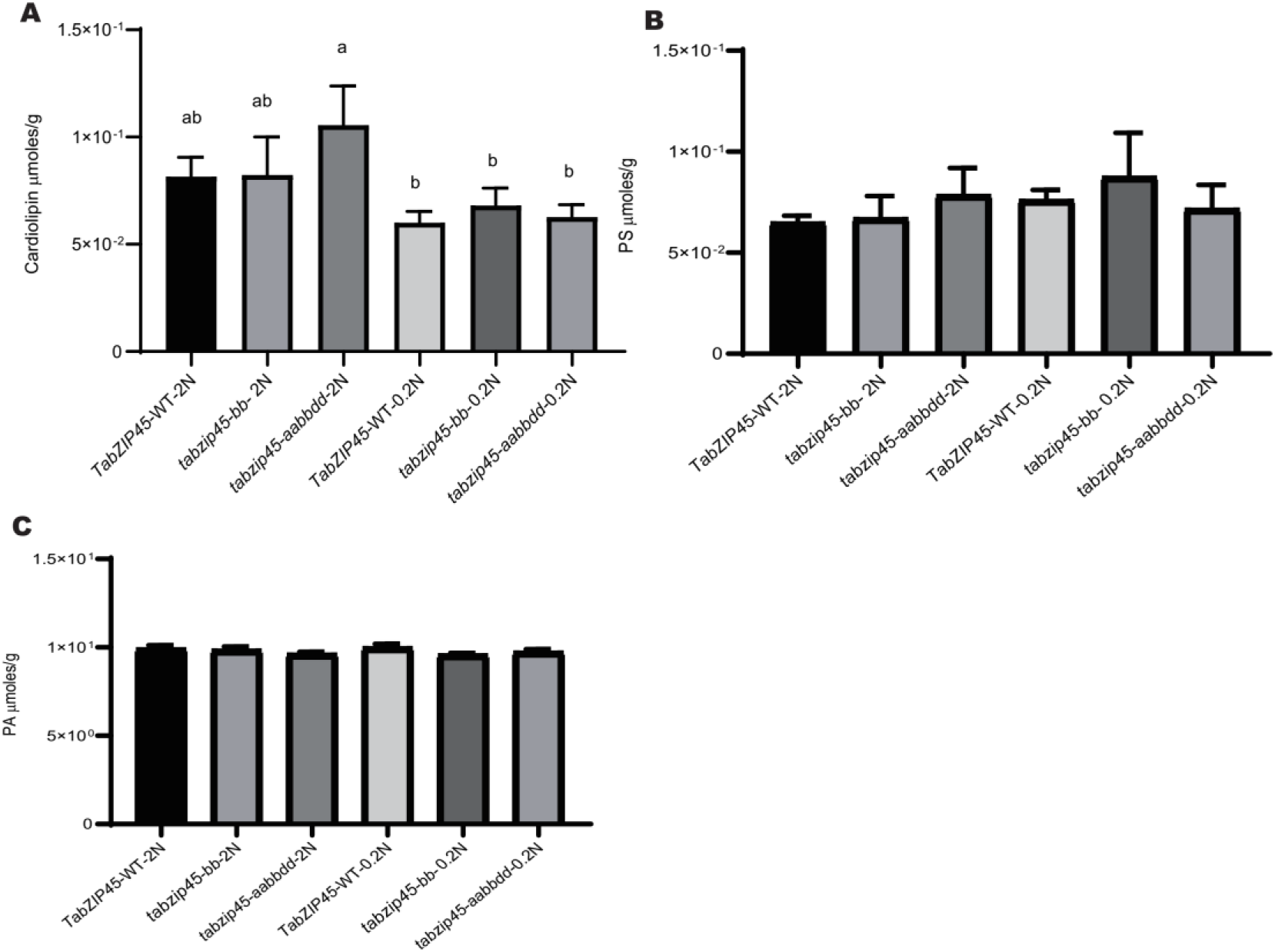
*TabZIP45-4B* did not affected cardiolipin, phosphatidic acid and phosphatidylserine. Phospholipids (**A**) Cardiolipin (**B**) Phosphatidylserine (**C**) Phosphatidic acid in tiller basal parts. Seedlings were grown under high N conditions (1.0 mM NH_4_NO_3_, 2N) or (0.1 mM NH_4_NO_3_, 0.2N) for two weeks and samples were collected. Data are mean ± S. E. (n ≥ 3). Different letters denote statistically significant differences (P < 0.05) from Fisher’s LSD ANOVA multiple comparisons. *tabzip45-aabbdd*, homozygous *TabZIP45-4A, 4B and 4D* triple mutant.

**Fig.S18.**
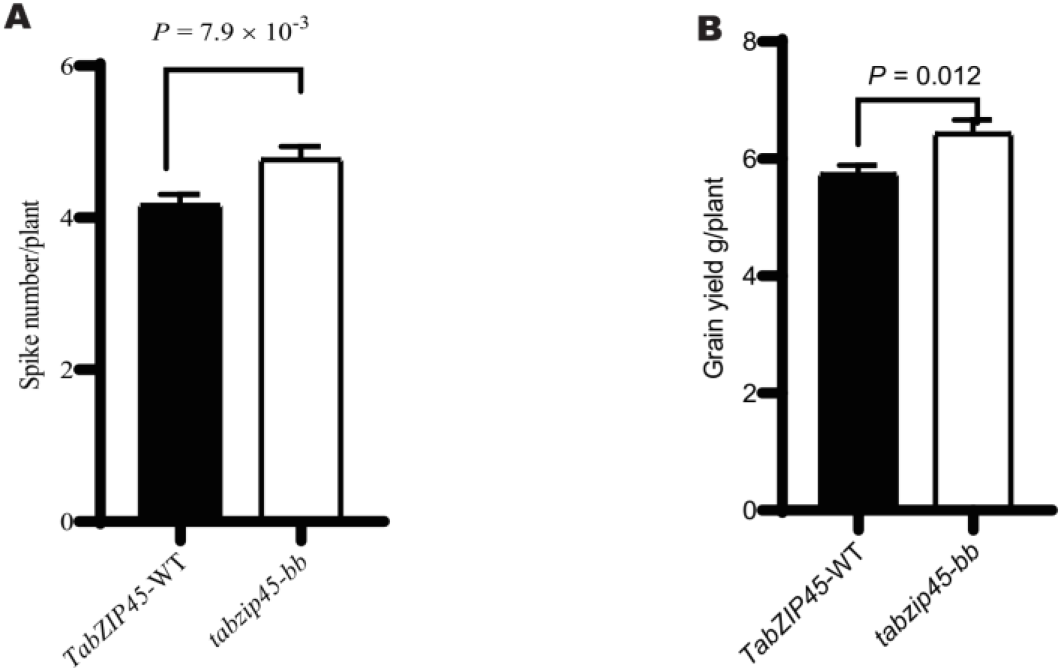
Knockout of *TabZIP45-4B* enhanced grain yield in drought condition. (**A**) Grain yield per plant. (**B**) Spike number per plant. Data in A and B are mean ± S. E. (n ≥ 40). *P* value in D is from a two tailed unpaired test. *TabZIP45-WT*, homozygous wild type offsprings separated from last generation; *tabzip45-bb*, homozygous mutant offsprings separated from last generation.

**Fig.S19.**
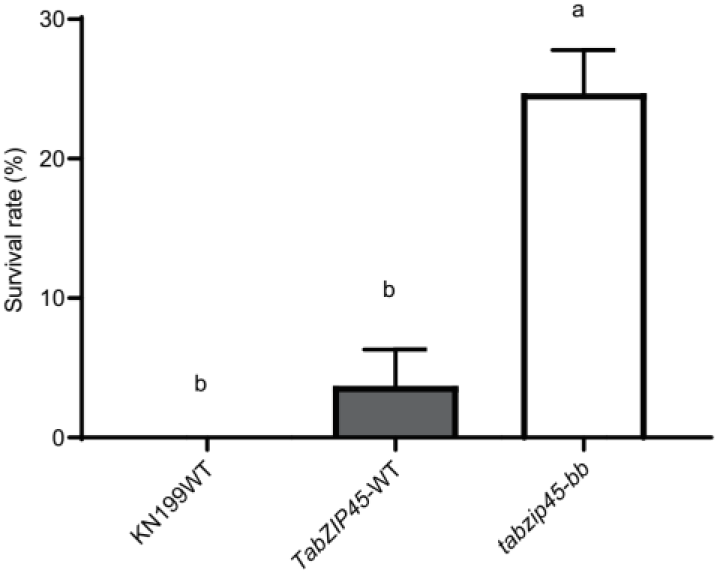
*TabZIP45-4B* was crucial for wheat freezing resistance. Statistical analysis of freezing survival rate in soil experiments. KN199WT, KN199 wild type. *TabZIP45-WT*, homozygous wild type offsprings. *tabzip45-bb*, homozygous mutant offsprings. Data are mean ± S. E. of representative plants (n = 9) from more than three independent experiments. Different letters denote statistically significant differences (*P* < 0.05) from one-way ANOVA multiple comparisons

**Fig.S20.**
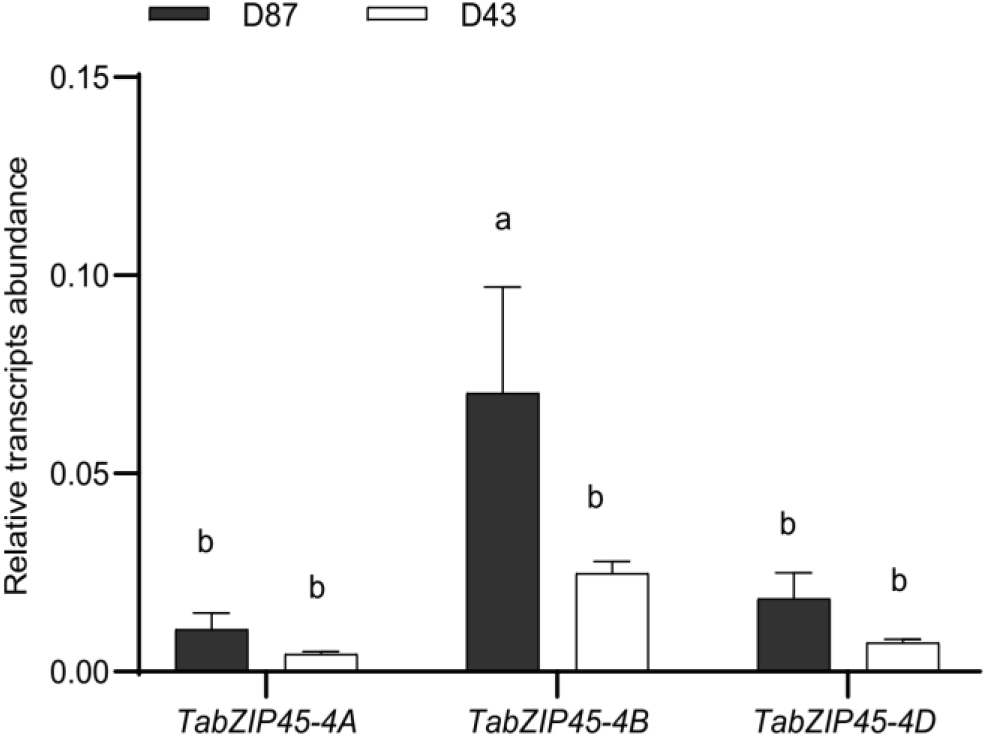
Dense planting promoted expression of *TabZIP45*. The above-ground parts of the seedlings were collected 20 days after sowing in 2019-2020 growing season in Beijing. The genes expression level was normalized to that of *Tatublin* as internal control. Data are mean ±S.E. (n = 3). Different letters denote statistically significant differences (*P* < 0.05) from Fisher’s LSD ANOVA multiple comparisons. D43 (43 seeds/m^2^) , D87 (87seeds/m^2^).

**Fig.S21.**
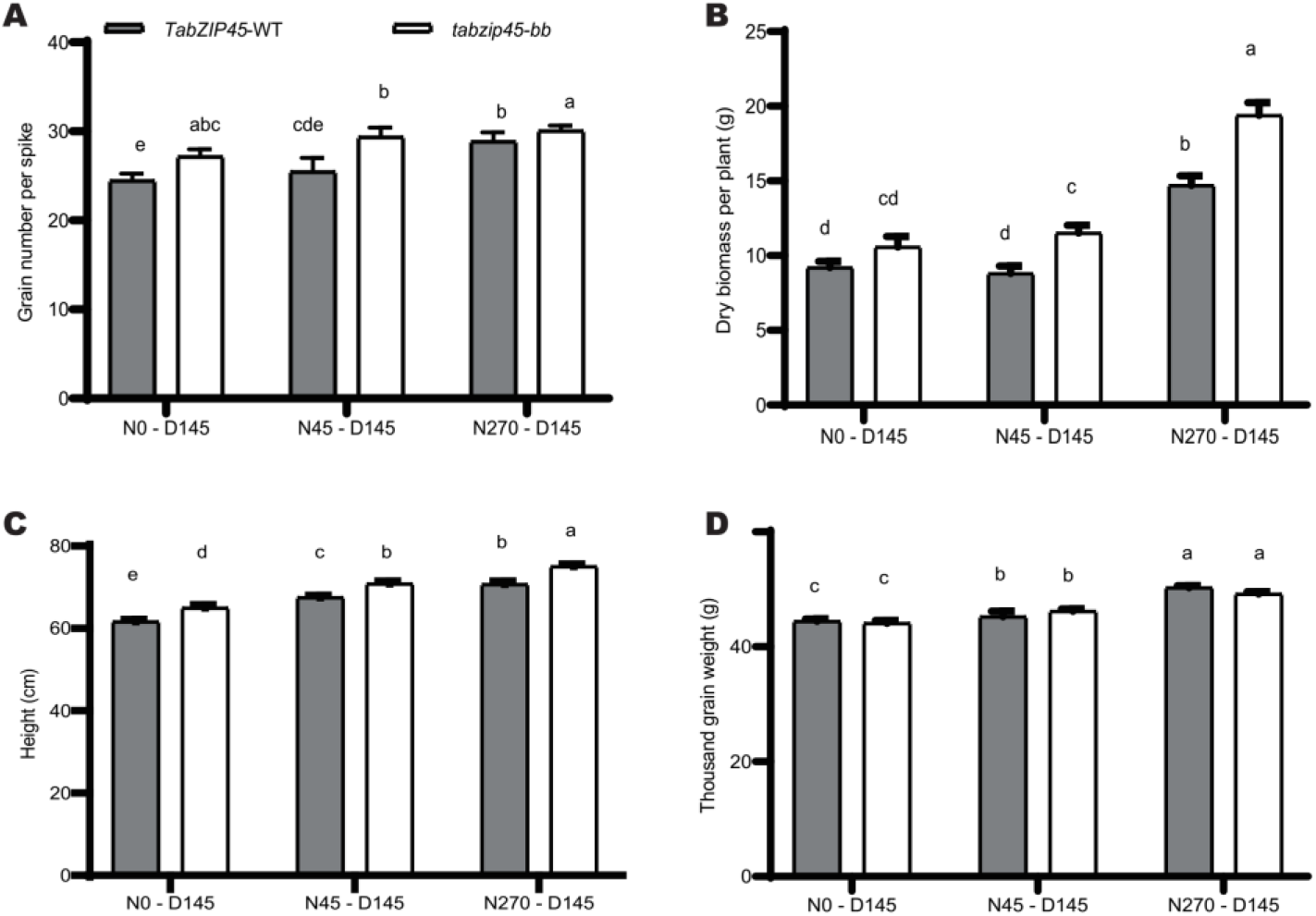
Knockout of *TabZIP45-4B* boosted grain yield under high planting density. **(A)** Grain number per spike. **(B)** Biomass per plant. **(C)** Height. (**D**) 1000-grain weight. *TabZIP45-WT*, homozygous wild type offsprings. *tabzíp45-bb*, homozygous mutant offsprings. Nitrogen supply level is 0 kg N/ha, 45 N kg/ha and 270 kg N/ha. Data are mean ± S. E. of representative plants (n = 40) from more than three independent experiments. Different letters denote statistically significant differences (*P* < 0.05) from two-way ANOVA multiple comparisons. In (A-D), N0 (0 kg N/ha), N45 (45 kg N/ha), N270 (270 kg N/ha). D145 (145 seeds/m^2^).

## Notes

### Competing Interest Statement

The authors have declared no competing interest.

