## Supplementary material for "Adaptation to climate change through dense planting for sustainable agriculture": All suplementary

**Supplementary materials**

Figs. S1 to S21

Tables S1 to S7

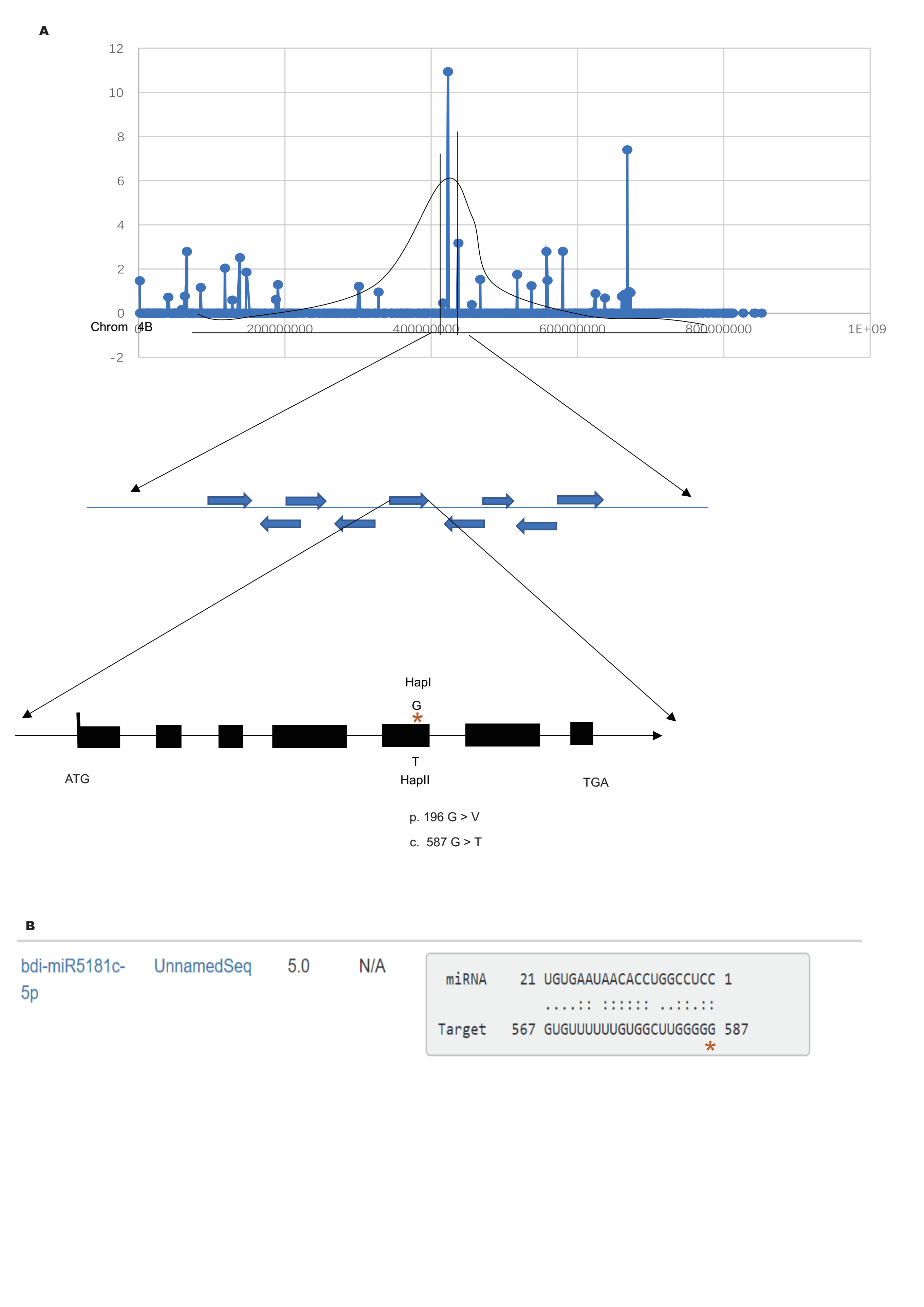

**Fig.S1. *TabZIP45* isolation and miRNA targeted to *TabZIP45-4B***

(**A**) the isolation of *TabZIP45-4B* by mapping of sequencing. (**B**) the mutation site is predicted to disrupt the binding sites of microRNA (http://plantgrn.noble.org/psRNATarget/).

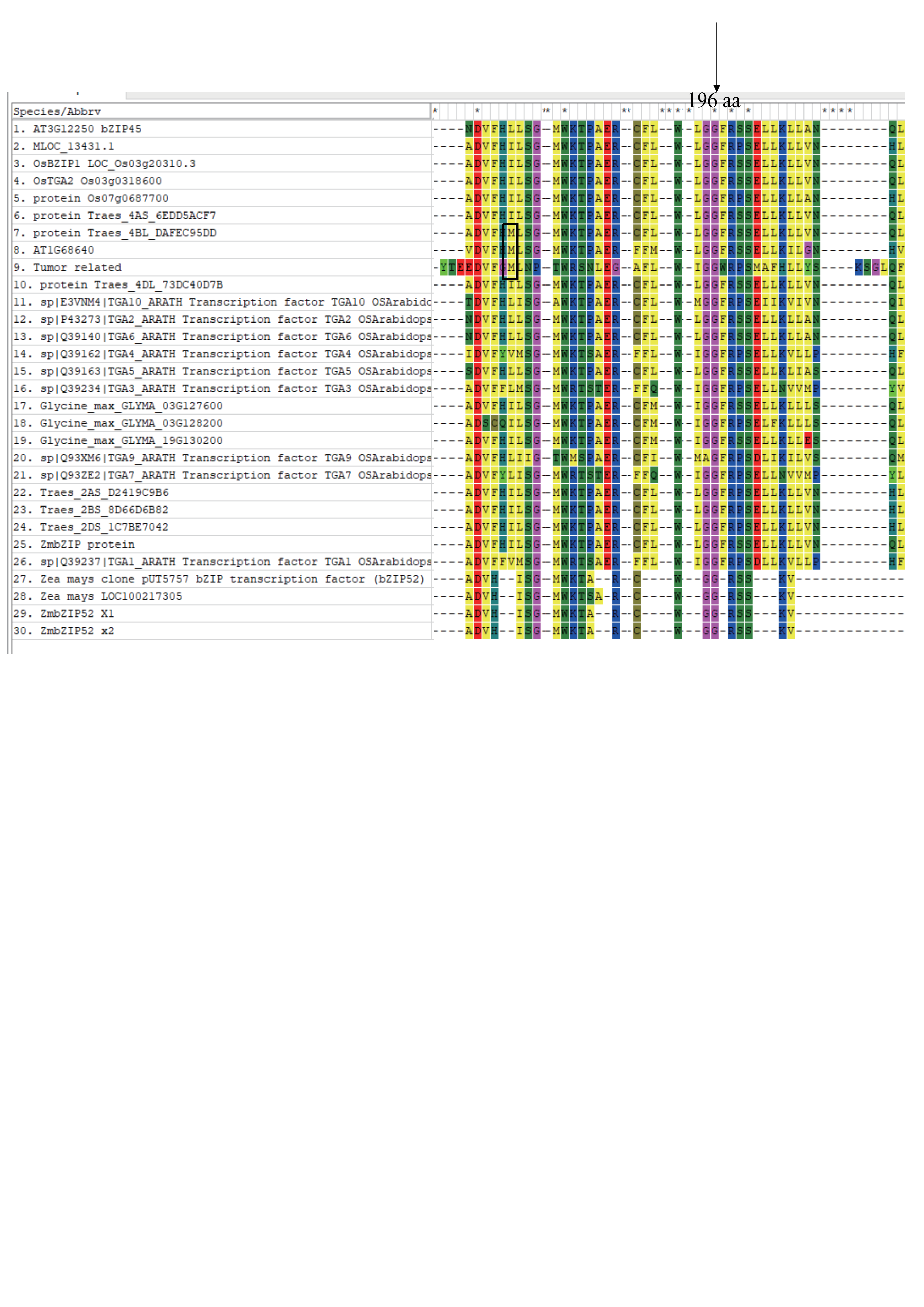

**Fig.S2. The TabZIP45 sequence conservation**

196 aa in DOG domain of TabZIP45-4B protein is conserved in angiosperms. The right above arrow indicates the conserved 196 aa (Glysine, G) in DOG domain of bZIP45 proteins in plants. The left above arrow indicates 178 aa difference among TabZIP45-4A, -4B and -4D. The Arrow pointed at dashed line box at middle left side of panel indicates the position of Methioine. The alignment is constructed by MEGA 6.0 by ClustalW method. Pairwise alignment and multiple alignment penalty is 8. Gap extension penalty is 0.025. Protein Weight Matrix is identity. Residue-specific penalties and hydrophilic penalties is on. Gap separation distance is 5. End Gap Separation is on. Use Negative Matrix, delay divergent cutoff is 15%. Keep predefined gaps.

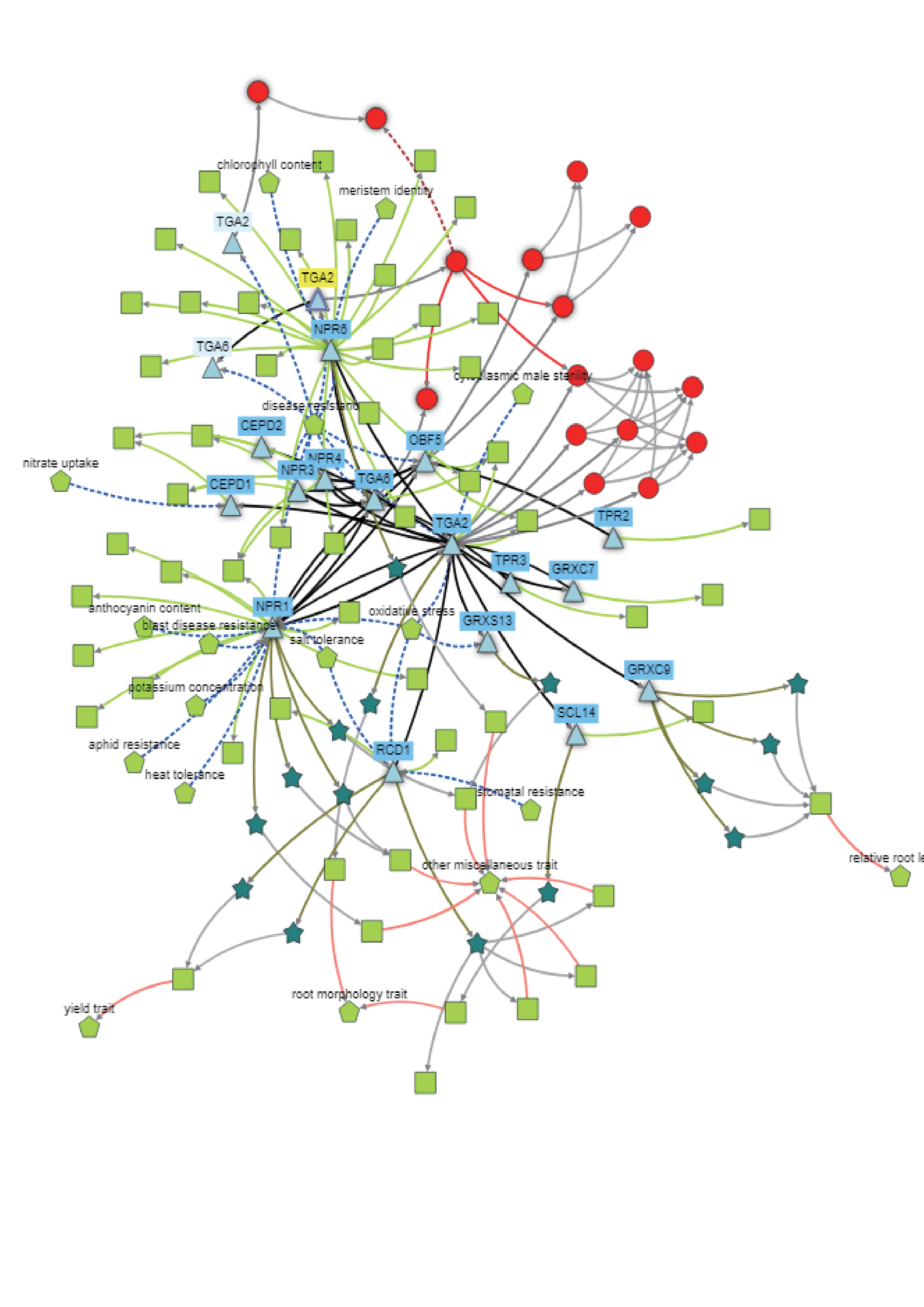

**Fig.S3. *TabZIP45* genetic network**

The network of *TabZIP45-4B* in nutrients sensation, growth regulation and stresses adaptation. The network was constructed in knetminer (<https://knetminer.com/Triticum_aestivum/>).

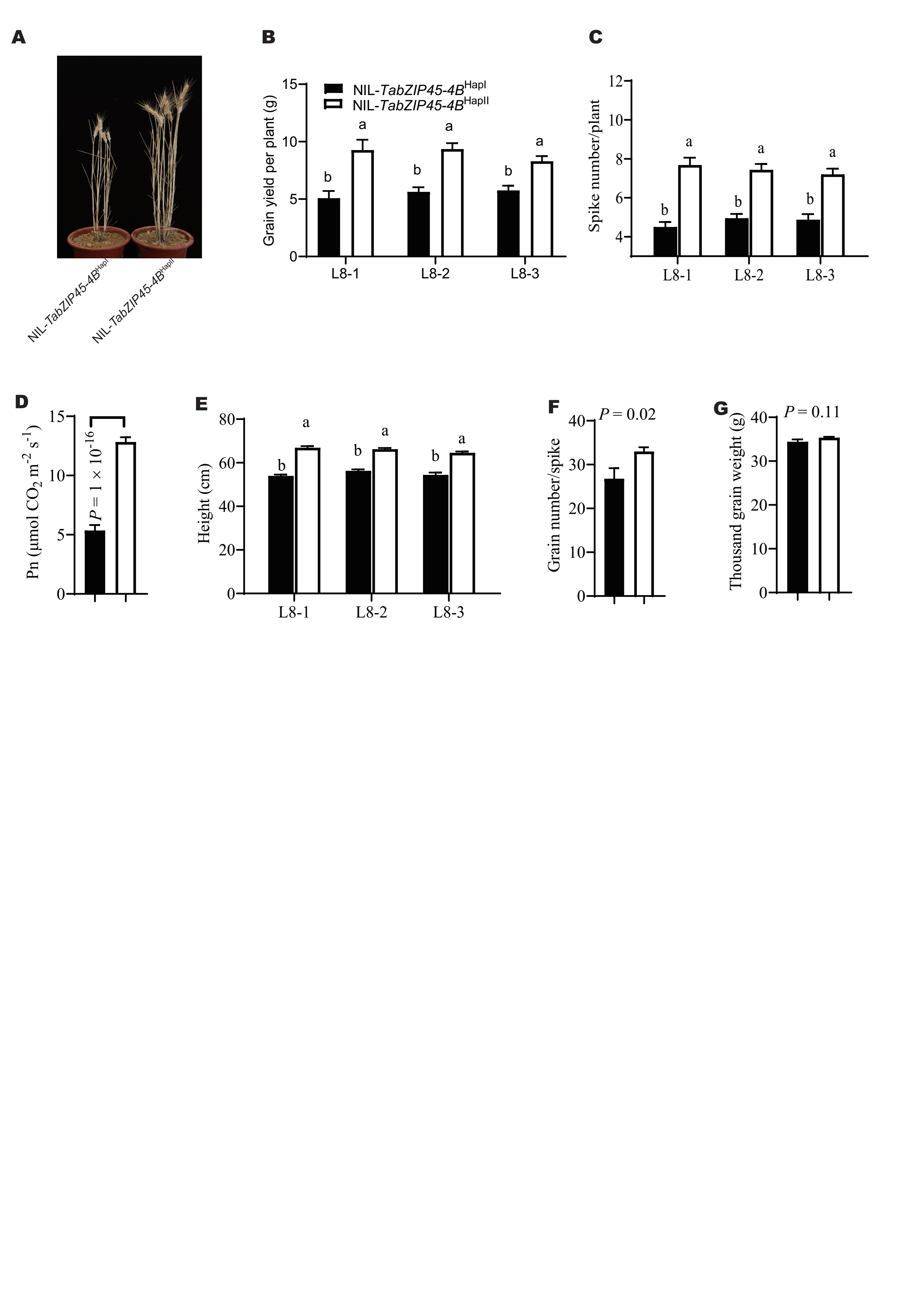

**Fig.S4. Agronomic traits of NIL-*TabZIP45-4B***^HapI^ **and NIL-*TabZIP45-4B***^HapII^ **under nitrogen limited conditions**

**(A)** Representative of more than four independent replicates. Scale bar = 10 cm. **(B)** Grain yield per plant. **(C)** Spike number per plant. **(D)** photosynthetic rates. (**E**) Height. (**F**) Grain number per spike. **(G)** 1000-grain weight. Data in B-G are mean ± S. E. (n = 40) and different letters denote statistically significant differences (*P* < 0.05) from two-way ANOVA multiple comparisons. L8-1, L8-2 and L8-3: different isogenic lines with NIL-*TabZIP45-4B*^HapII^ or NIL-*TabZIP45-4B*^HapI^*.* Nitrogen supply level: 30 kg N/ha. Isogenic lines NIL-*TabZIP45-4B*^HapI^ and NIL-*TabZIP45-4B*^HapII^ are generated by introducing *TabZIP45-4B*^HapI^ from J411 into XY54 (BC5F5).

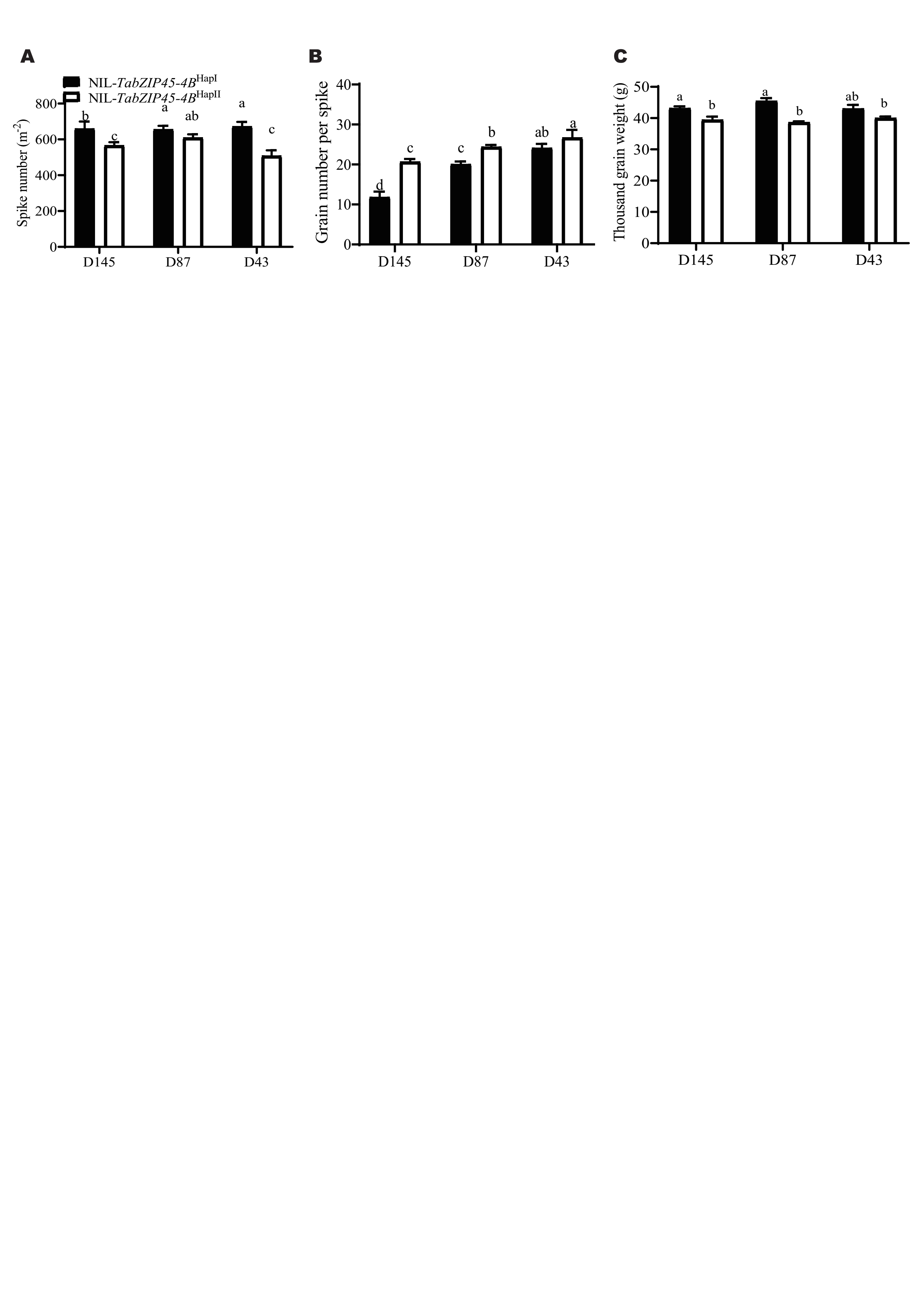

**Fig.S5. Agronomic traits of NIL-*TabZIP45-4B***^HapI^ **and NIL-*TabZIP45-4B***^HapII^ **under different planting densities.**

(**A**) Spike number per plant (**B**) Grain number per spike and (**C**) Thousand grain weight under different nitrogen supply levels and sowing densities. Data in A and B are mean ± S. E. (n ≥ 34) Isogenic lines NIL-*TabIP45-4B*^HapI^ and NIL-*TabZIP45-4B*^HapII^ are generated by introducing NIL-*TabZIP45-4B*^HapII^ from J411 into XY54 (BC5F6). In (A-C), D43 (43 seeds/m^2^), D87 (87 seeds/m^2^), D145 (145 seeds/m^2^).

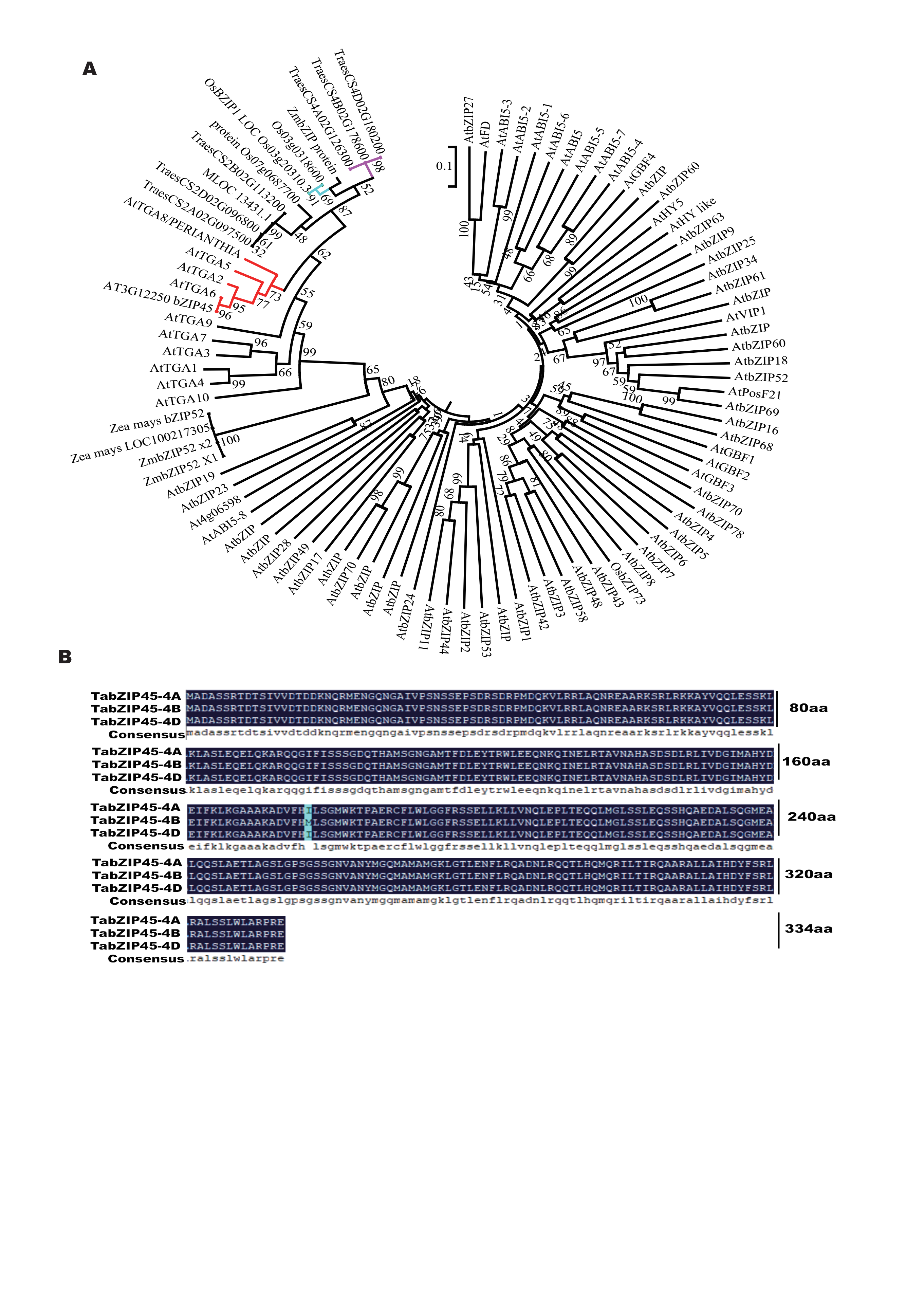

**Fig.S6. TabZIP45 phylogeny tree and sequence**

(**A**) The phylogeny tree is draw by neighbor-joining method of MEGA 6.0. (**B**) Coding sequences of *TabZIP45-4A, -4B* and -*4D* cloned in KN199, which was generated by DNAMAN (LynnonBiosoft, USA).

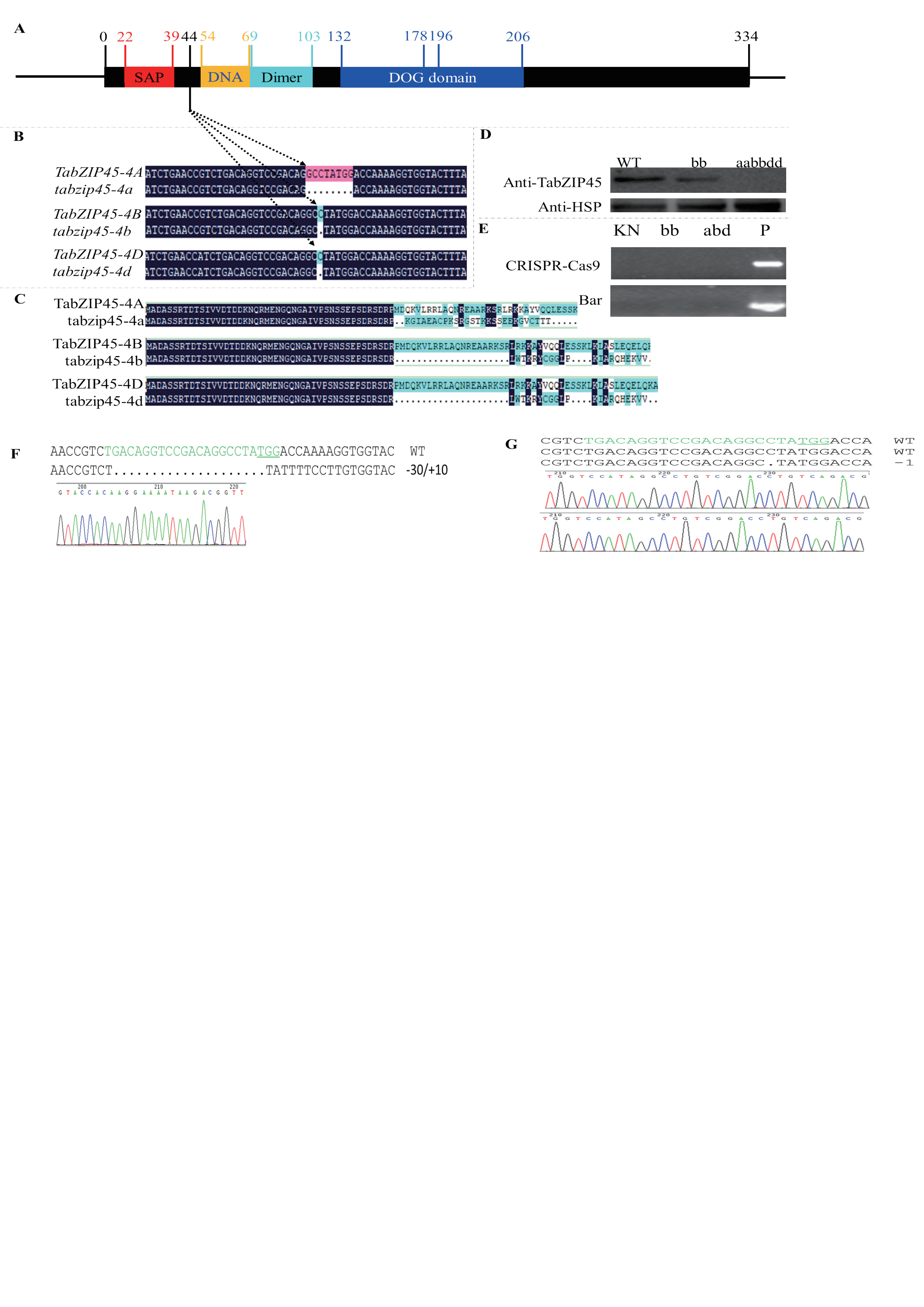

**Fig.S7. TabZIP45 protein structure domain and *TabZIP45* gene editing sites**

(**A**) Structure domain of TabZIP45, number stands for aa sites. Red region, Ser-Ala-Pro (SAP) motif. Yellow region, DNA binding sites domain of TabZIP45. Cyan region, Dimmer interface domain. Blue region, Delay of Germination (DOG) domain. Original amino acids site (aa) in 178 between TabZIP45-4A, 4D and 4B is different. Two alleles of different haplotypes encode different aa in 196. (**B**) Genome editing leads to 8 bp insertion in *TabZIP45-4A,* one bp deletion in *TabZIP45-4B* and *TabZIP45-4D* respectively*.* (**C**) Premature stop in *TabZIP45-4A, 4B* and*4D* caused by genome editing produce 73 aa, 66 aa and 66 aa mutant protein respectively, without original structure domain. (**D**) The mutants are confirmed by TabZIP45 antibody. (**E**) Determine whether the plants have CRISPR-Ca9 and anti-Basta (Bar) genes. In (D) and (E), WT, homozygous wild type offsprings separated from last generation; *bb*, homozygous mutant offsprings separated from last generation; *aabbdd*, *TabZIP45-4A*, *4B* and *4D* all homozygous mutant; P, plasmid positive control. (**F** and **G**) the sequence results of two independent *TabZIP45-4B* mutants.

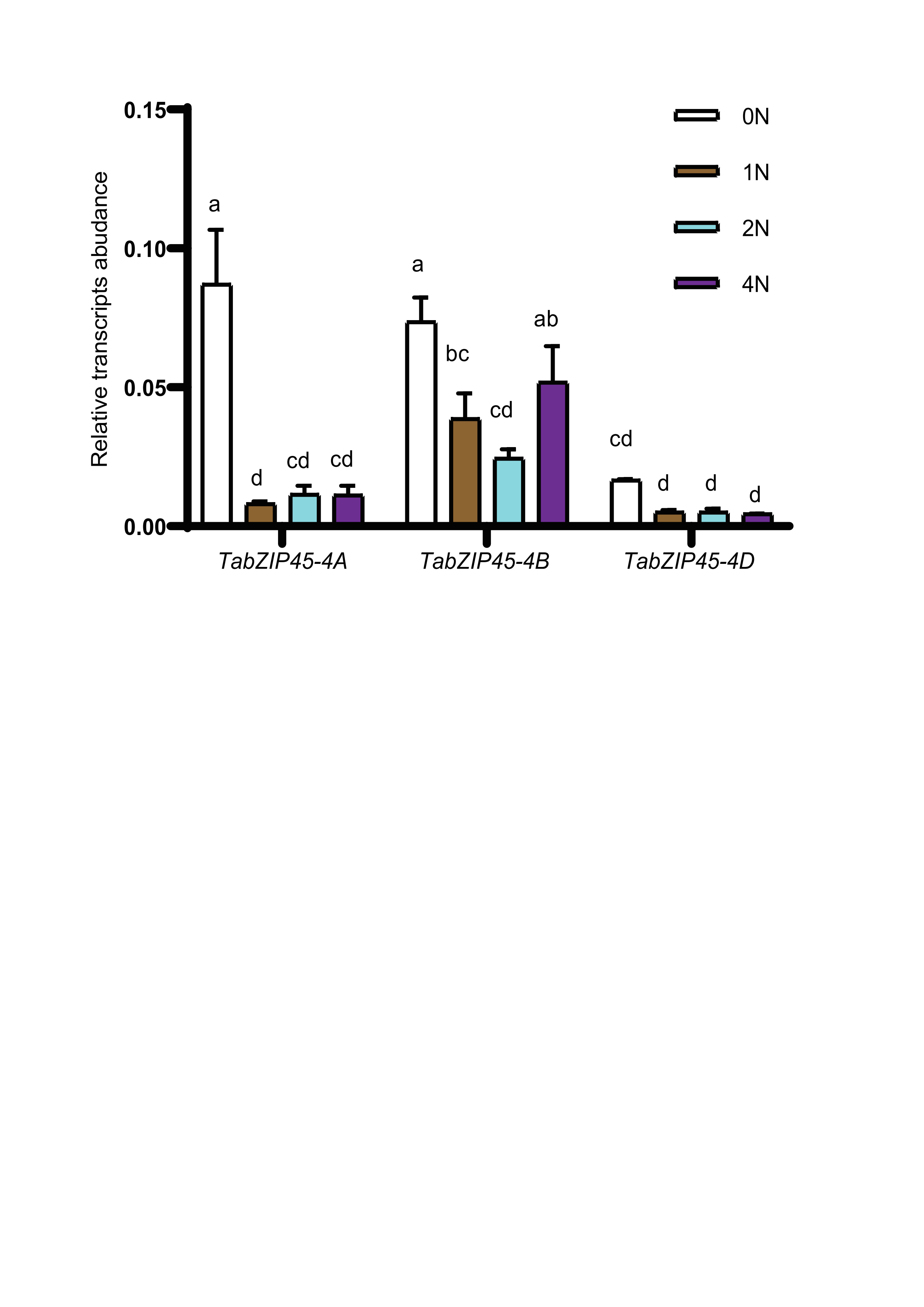

**Fig.S8. The response of *TabZIP45* to nitrogen supply**

The seedlings of KN199WT were grown for 4 weeks in the nutrient solutions that contained 0.0 mM NH_4_NO_3_ (0N), 0.5 NH_4_NO_3_ (1N), 1.0 NH_4_NO_3_ (2N) and 2.0 NH_4_NO_3_ (4N). The shoots were collected for gene expression analysis. Data are mean ± S. E. (n = 4). Different letters denote statistically significant differences (*P* < 0.05) from Fisher’s LSD ANOVA multiple comparisons. The genes expression level was normalized to that of *Taactin* as internal control.

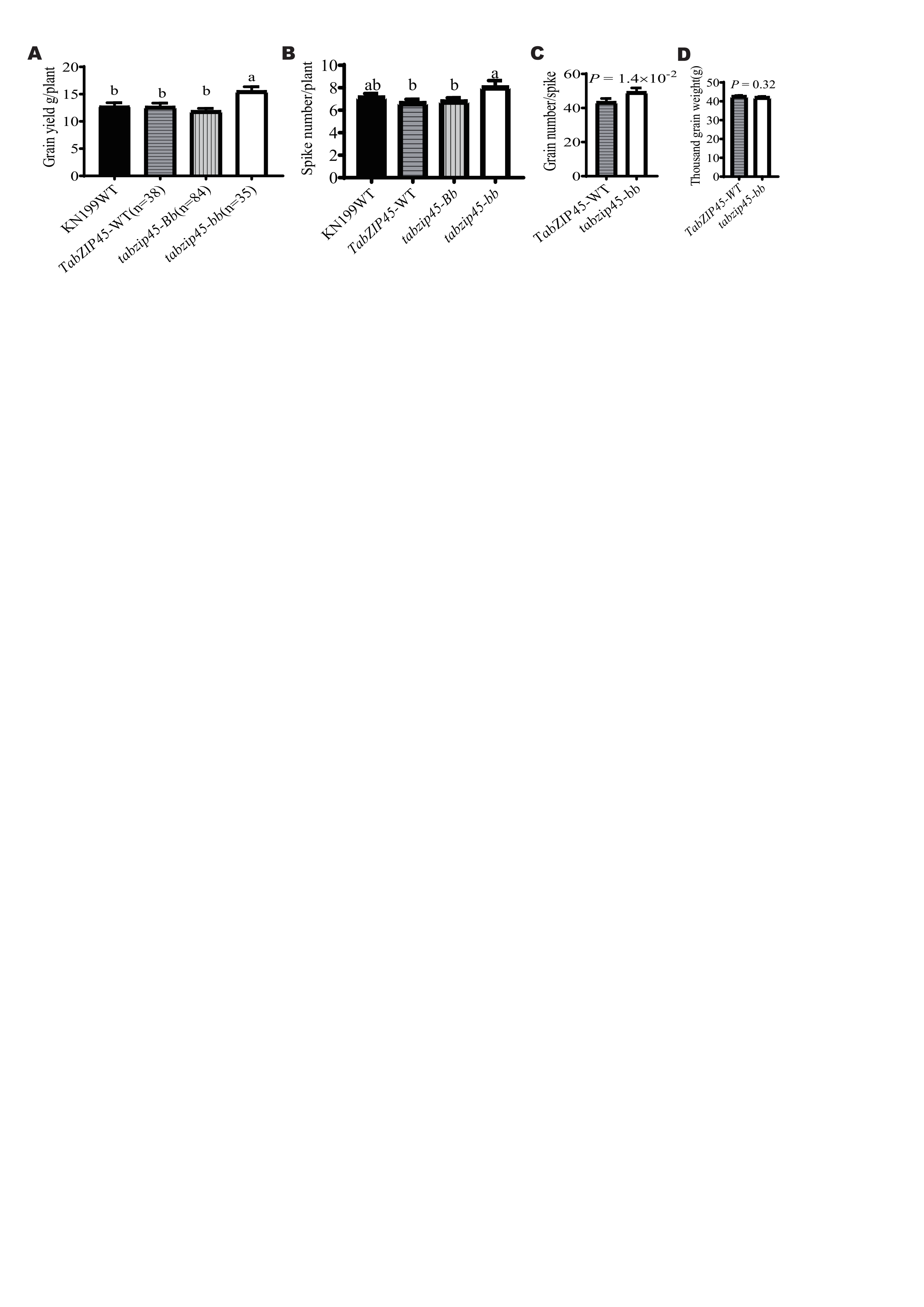

**Fig.S9. Knockout of *TabZIP45-4B* increased grain yield**

**(A)** Grain yield per plant. **(B)** Spike number per plant. Data in A and B are mean ± S. E. (n ≥ 35). Different letters in A-B denote statistically significant differences (*P* < 0.05) from Fisher’s LSD ANOVA multiple comparisons. **(C)** Grain number per spike. *P* value in C is from a Mann-Whitney *t* test. **(D)** 1000-grain weight. *P* value in C-D is from a two tailed unpaired test. In A-D: KN199WT, KN199 wild type; *TabZIP45*-WT, homozygous wild type offsprings separated from last generation; *tabzip45-Bb*, heterozygous mutant offsprings separated from last generation; *tabzip45-bb*, homozygous mutant offsprings separated from last generation. Nitrogen supply level (270 kg N/ha).

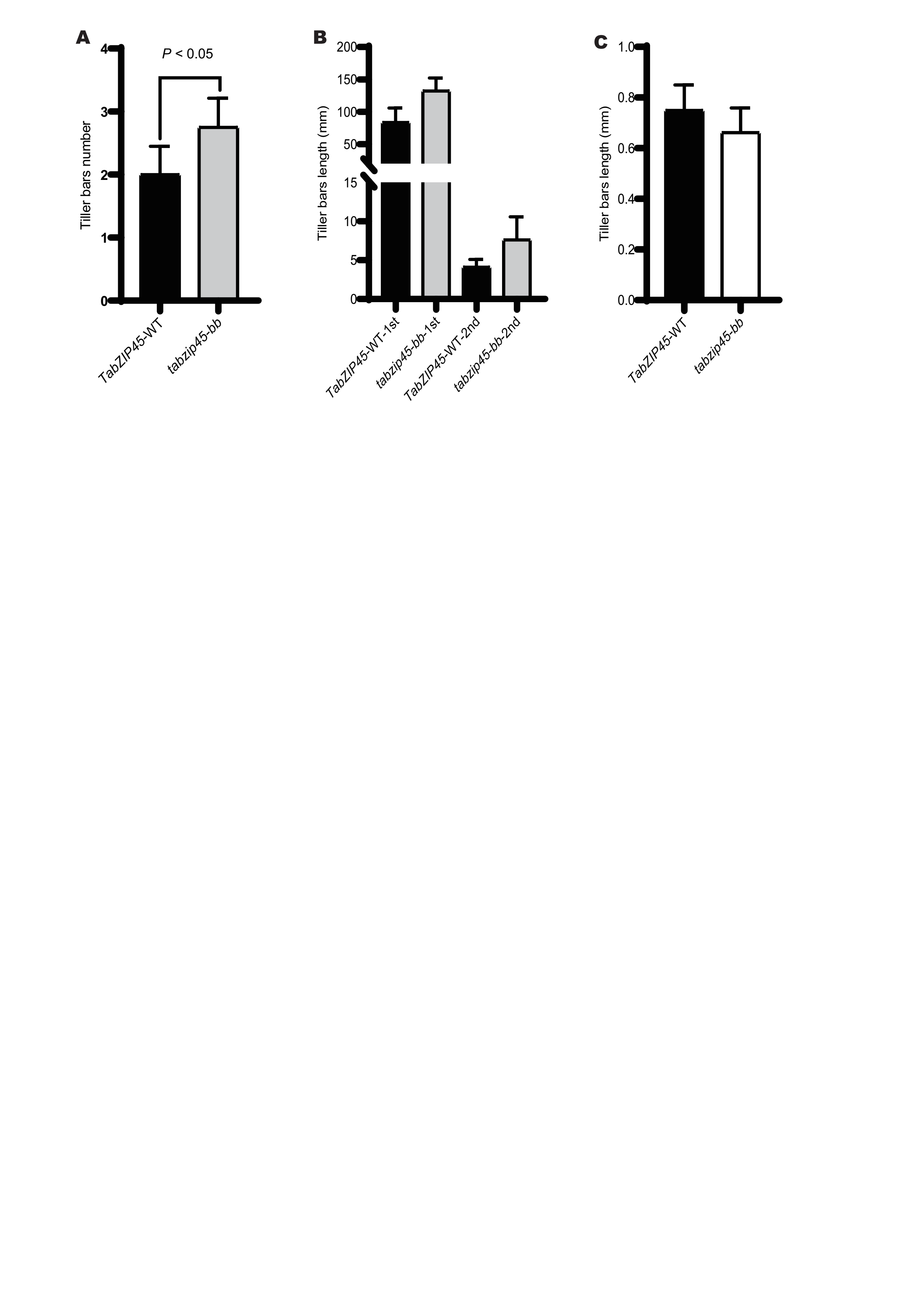

**Fig.S10. *TabZIP45-4B* inhibited tiller initiation and tiller elongation**

(**A**) Tiller buds number in 2N. (**B**) Tiller buds length in 2N. (**C**) Tiller buds number in 0.2N. Seedlings were grown under high N conditions (1.0 mM NH4NO3, 2N) or (0.1 mM NH4NO3, 0.2N) for two weeks and samples were collected. Data are mean ± S. E. (n = 8) and *P* values in A-C were from paired Student’s *t*-test. 1st is first tiller, 2nd is second tiller. *TabZIP45*-WT, homozygous wild type offsprings. *tabzip45-bb*, homozygous mutant offsprings.
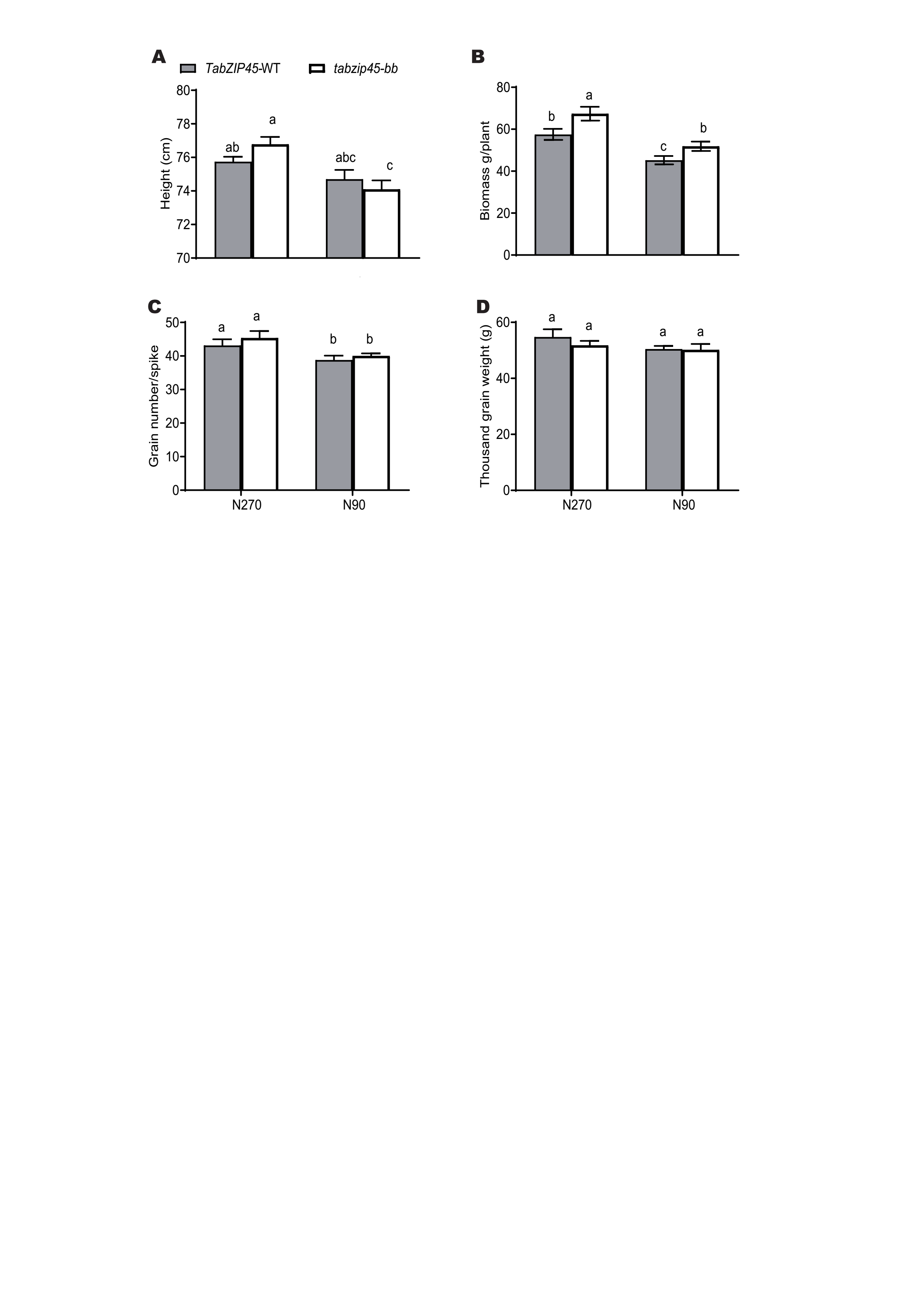

**Fig.S11. Knockout of *TabZIP45-4B* increased grain yield**

**(A)** Height. **(B)** Biomass per plant. **(C)** Grain yield per plant. (**D**) Grain number per spike. (**E**) 1000-grain weight. Data in A-E are mean ± S. E. (n = 40) and different letters denote statistically significant differences (*P* < 0.05) from two-way ANOVA multiple comparisons. *TabZIP45*-WT, homozygous wild type offsprings. *tabzip45-bb*, homozygous mutant offsprings. In (A-D), (90 kg N/ha), N270 (270 kg N/ha).

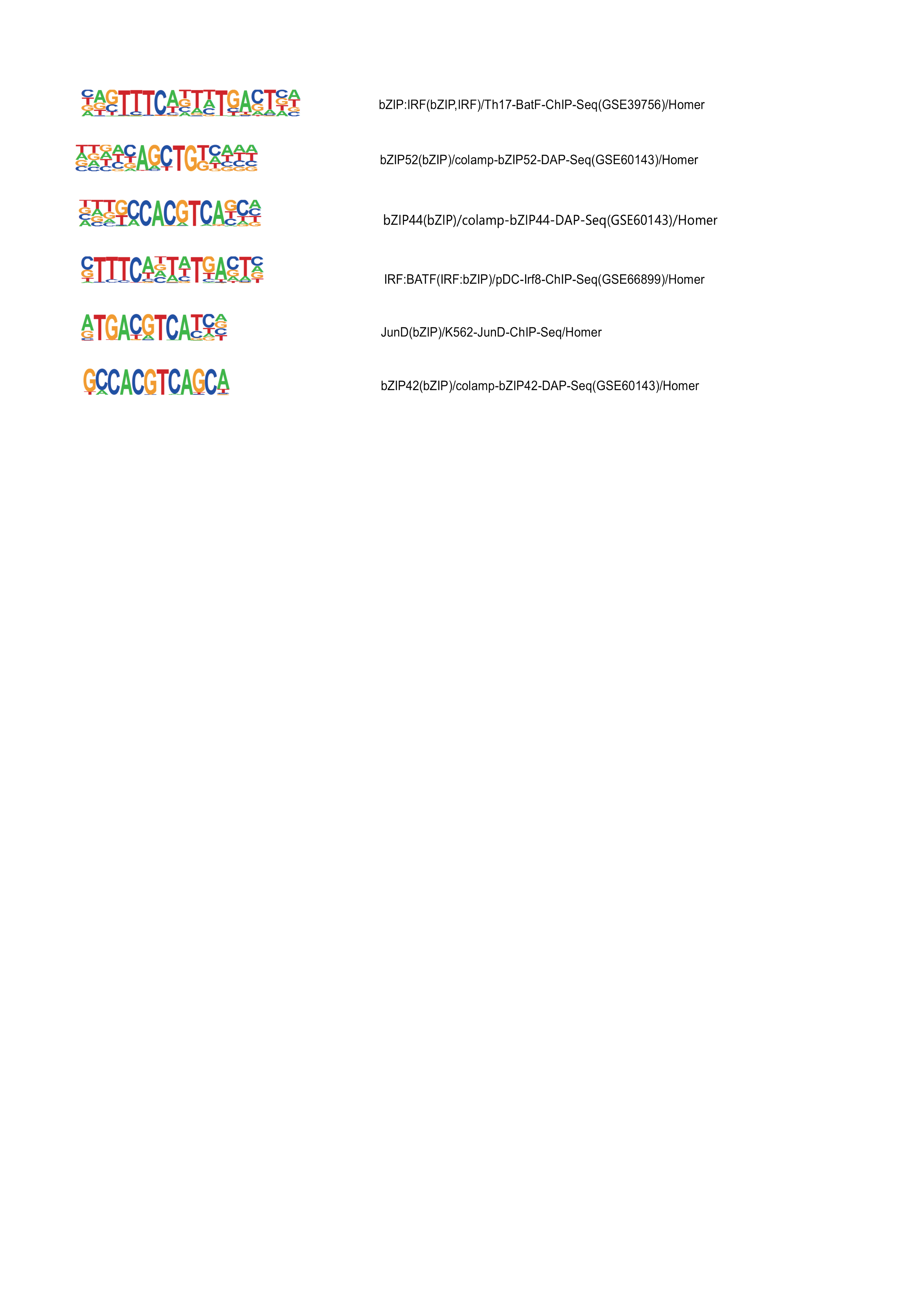

**Fig.S12. TabZIP45 binding sequence motif**

The sequence motif of bZIP family enriched by ChIP-seq.

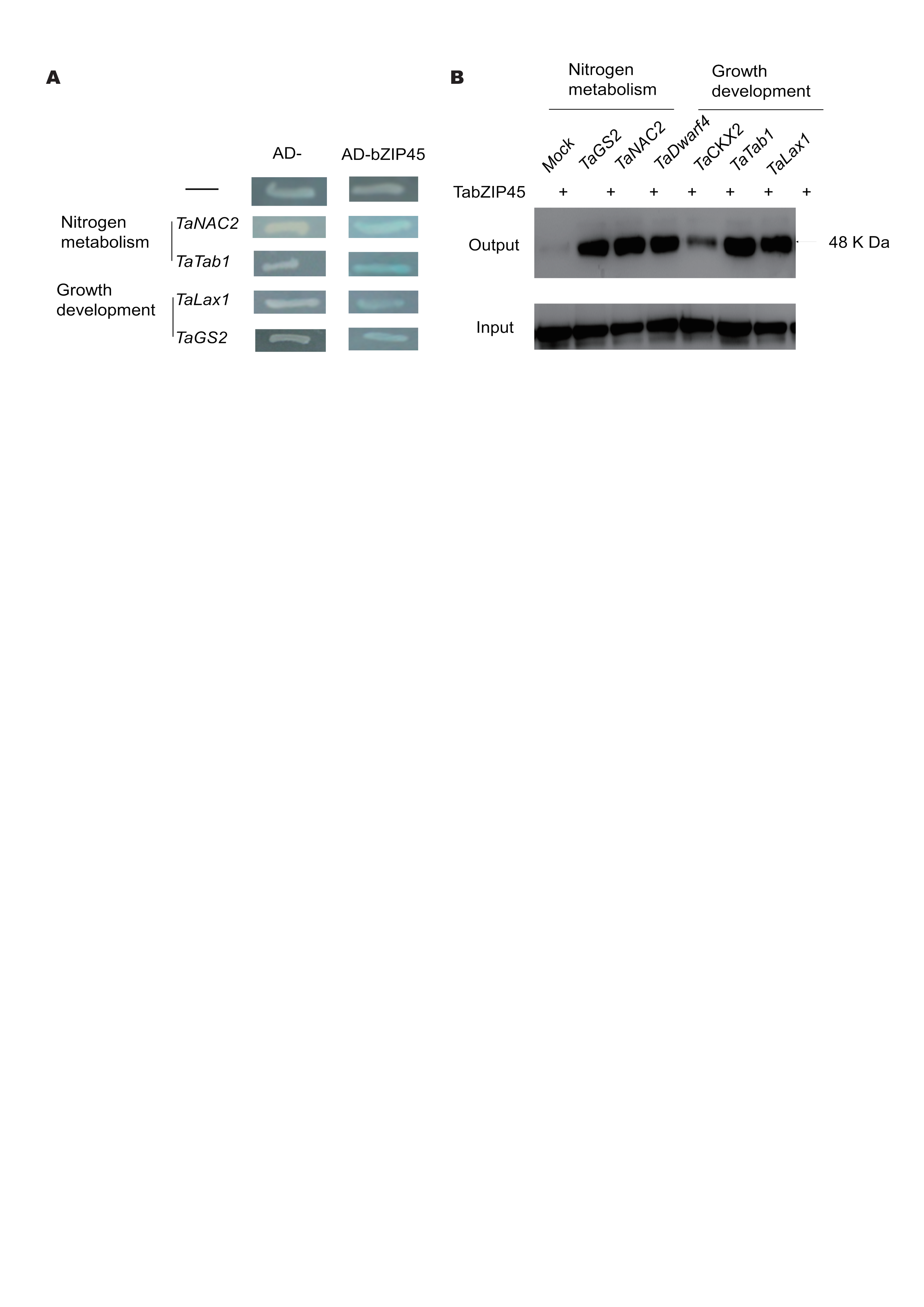

**Fig.S13. The binding ability of TabZIP45-4B to the promoters of downstream genes**

**(A)** Yeast one hybrid using TabZIP45-4B and the promoters of candidate downstream genes; **(B)** Prokaryotic purified TabZIP45-4B was incubated with different genes’ promoters with biotin marked at 5′ end of both DNA strands and probed with biotin antibody.

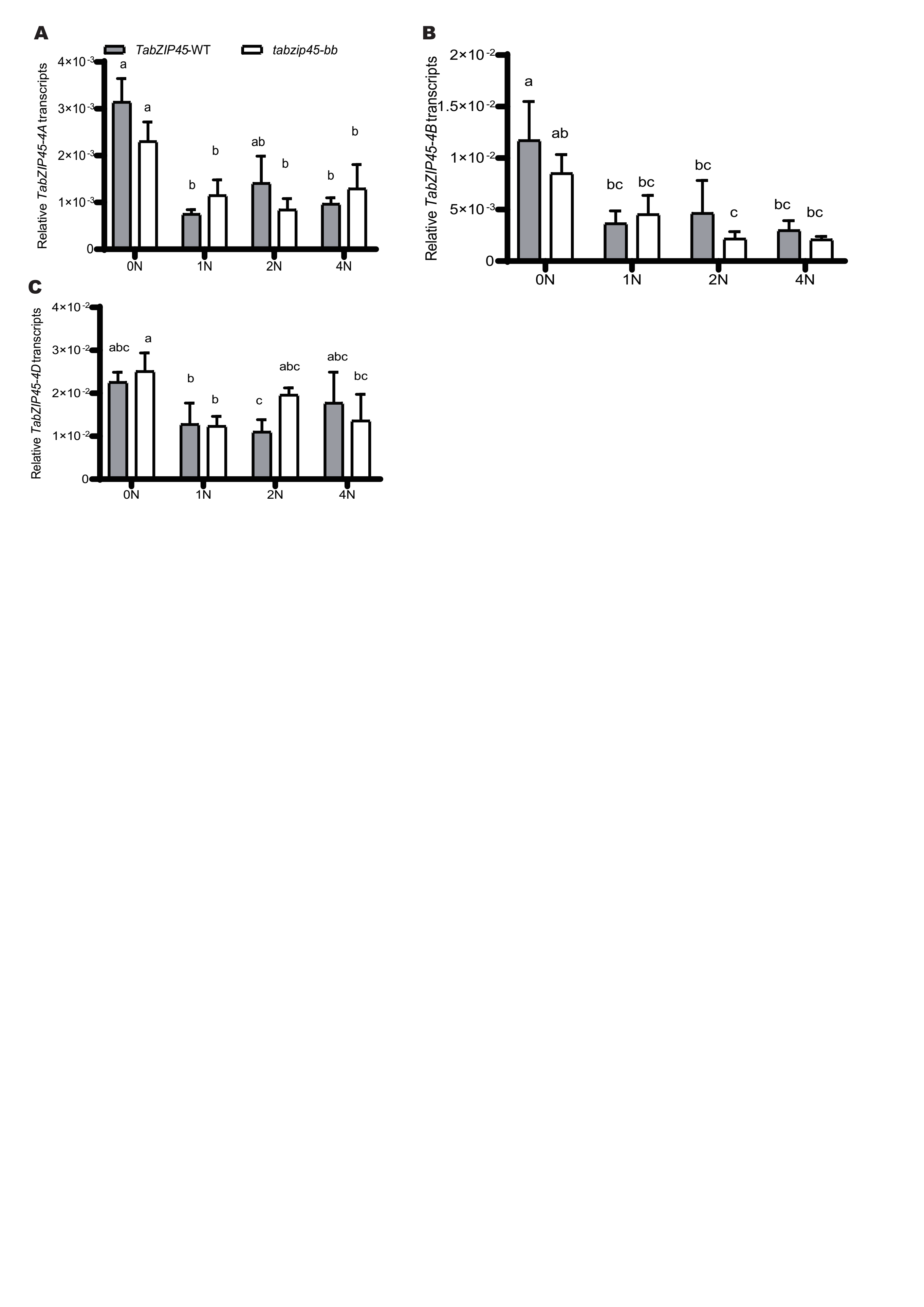

**Fig.S14. Knockout of *TabZIP45*-*4B* did not affect *TabZIP45-4A*, *-4B*, *-4D* expression.**

(**A**) *TabZIP45-4A,* (**B**) *TabZIP45-4B*, (**C**) *TabZIP45-4C*. The seedlings of *TabZIP45-4B-WT* and *tabzip45-bb* mutant were grown for four weeks in the nutrient solutions which contain 0.00 mM NH_4_NO_3_ (0N), 0.50 mM NH_4_NO_3_ (1N), 1.00 mM NH_4_NO_3_ (2N) and 2.00 mM NH_4_NO_3_ (4N). Then the shoots and roots were collected for gene expression analysis. *TabZIP45*-WT, homozygous wild type offsprings. *tabzip45-bb*, homozygous mutant offsprings. Data in A-D were from shoots, and data in E and F were from roots. The genes expression level was normalized to that of *Tatublin* as internal control. Data are mean ± S. E. (n ≥ 3). Different letters denote statistically significant differences (*P* < 0.05) from Fisher’s LSD ANOVA multiple comparisons.

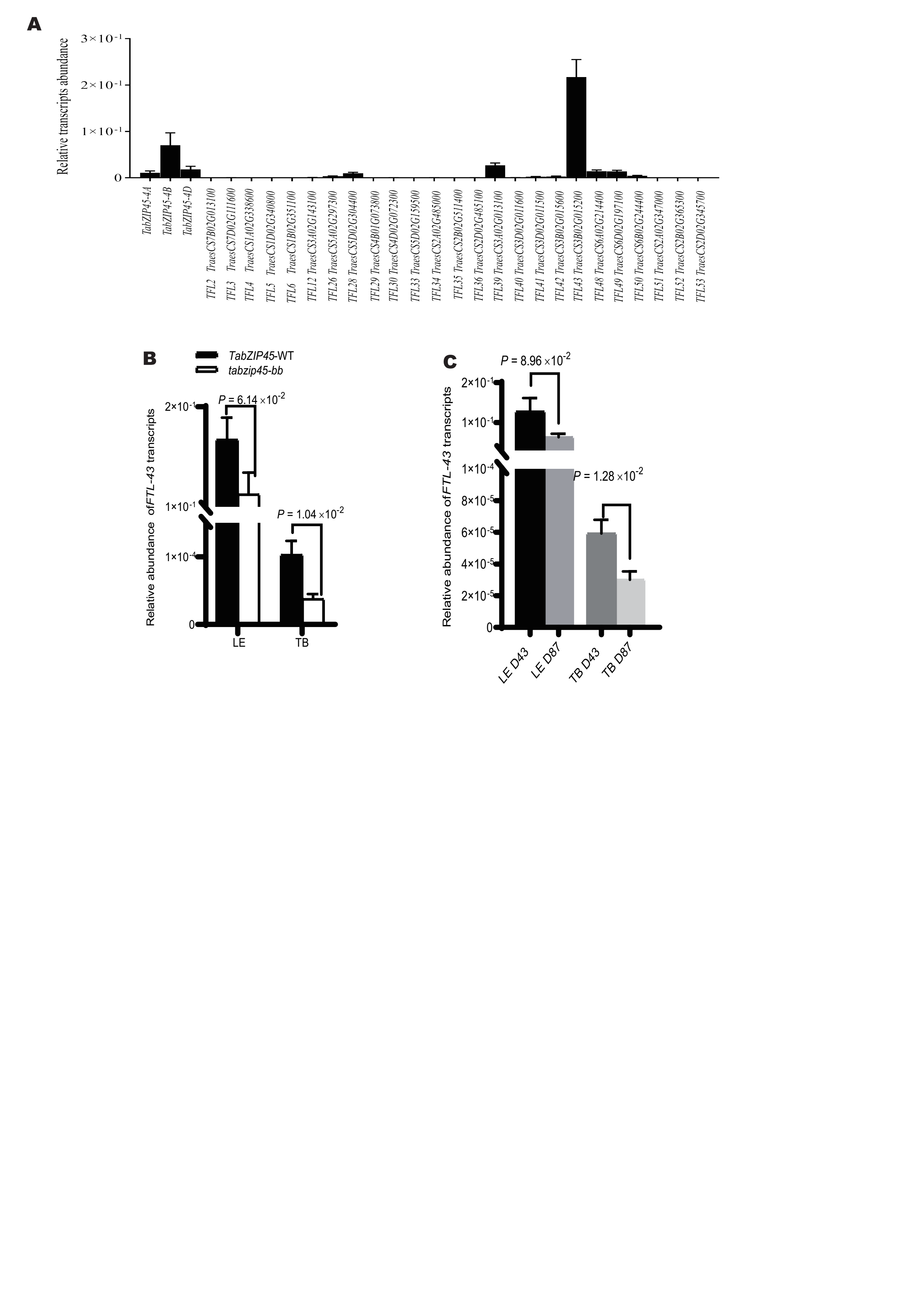

**Fig.S15. *TaFTL43* was essential in wheat development and interaction with environment**

**(A**) Relative expression of *TaFTL* genes in above ground parts collected 27 days after sowing in Hebei province under 45 kg N/ha supply. **(B-C)** Relative expression of *TaFTL43* (in leaves (LE) and tiller basal parts (TB) collected 27 days after sowing. The genes expression level was normalized to that of *Tatublin* as internal control. Data are mean ± S. E. (n = 4). *P* values in B-C are from a two tailed unpaired Mann-Whitney or Welch’s *t* test depends on data normality significance. In (D), D43 (43 seeds/m^2^), D87 (87seeds/m^2^).

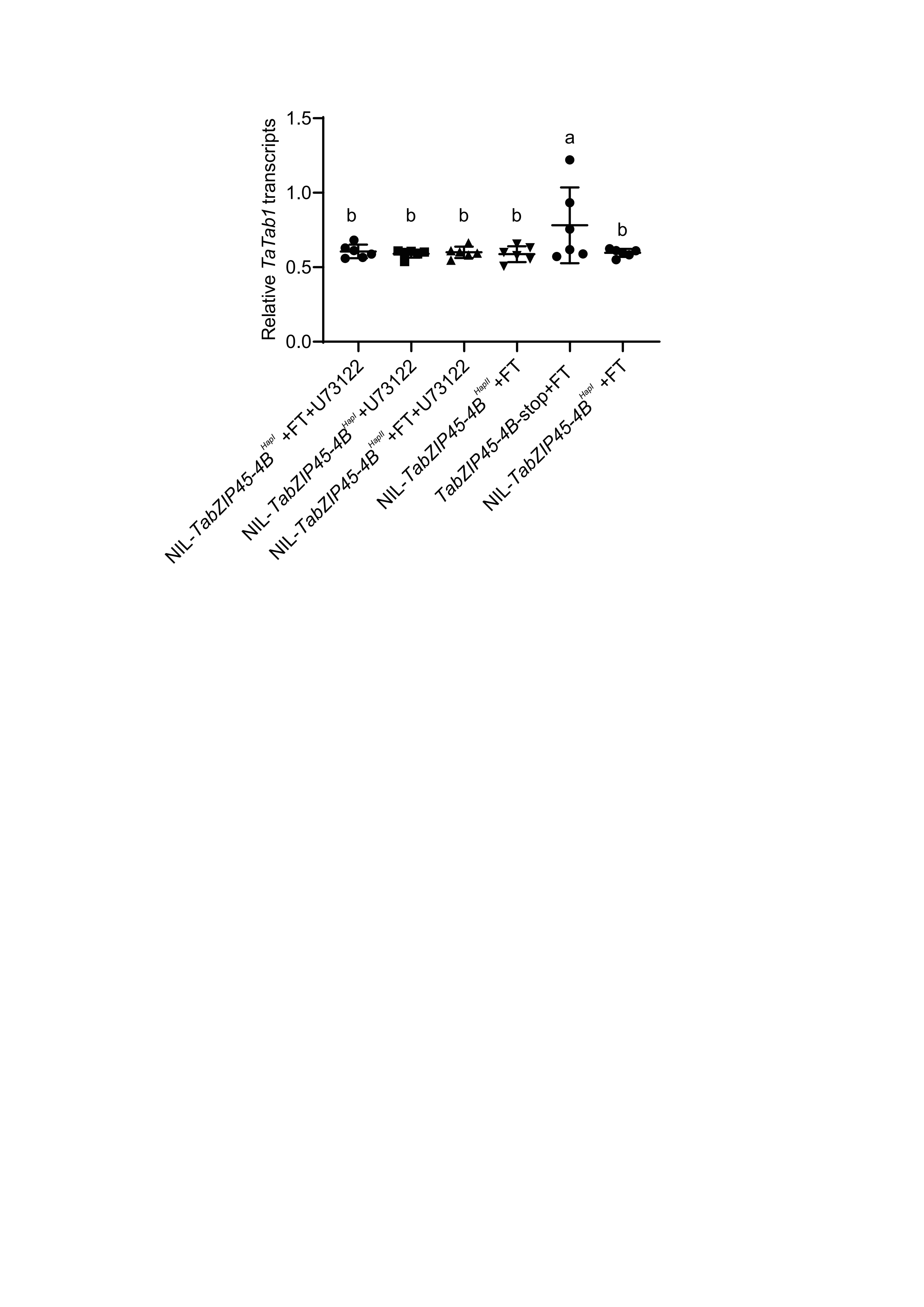

**Fig.S16. *TabZIP45*-*4B* inhibited *Tatab1* expression at presence of TaFTL-43**

The knock out *TabZIP45*-*4B* promoted *Tatab1* promoter transcription. The different haplotype of *TabZIP45*-*4B* did not affect transcriptional activation on *Tatab1*. FT, TaFTL-43; U73122, Phospholipase C (PLC) inhibitor. Relative luciferase was normalized to internal luciferase reference. Data were presented as mean ± S.E. (n = 6)

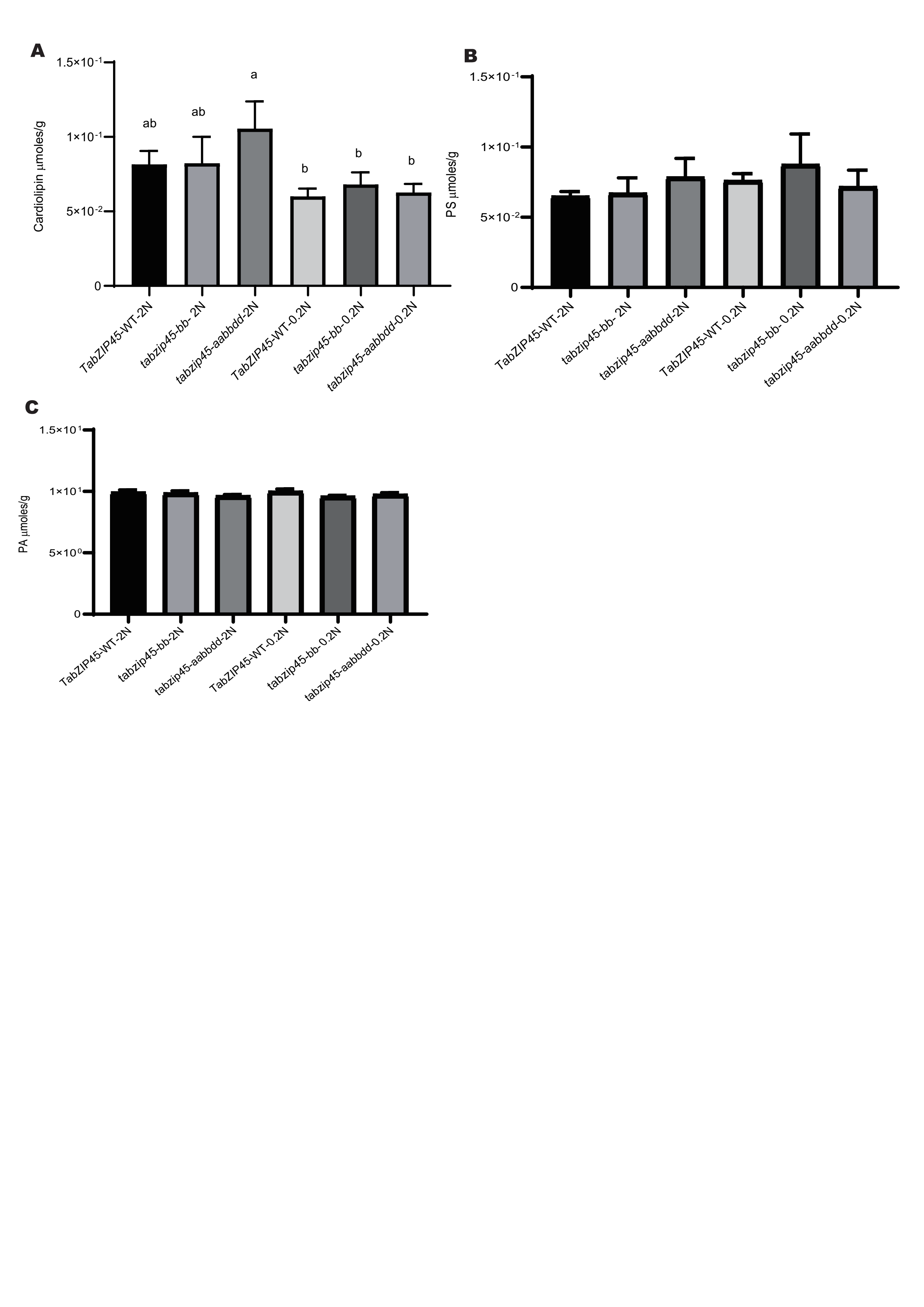

**Fig.S17. *TabZIP45*-*4B* did not affected cardiolipin, phosphatidic acid and phosphatidylserine**

Phospholipids (**A**) Cardiolipin (**B**) Phosphatidylserine (**C**) Phosphatidic acid in tiller basal parts. Seedlings were grown under high N conditions (1.0 mM NH_4_NO_3_, 2N) or (0.1 mM NH_4_NO_3_, 0.2N) for two weeks and samples were collected. Data are mean ± S. E. (n ≥ 3). Different letters denote statistically significant differences (P < 0.05) from Fisher’s LSD ANOVA multiple comparisons. *tabzip45-aabbdd*, homozygous *TabZIP45-4A, 4B and 4D* triple mutant.

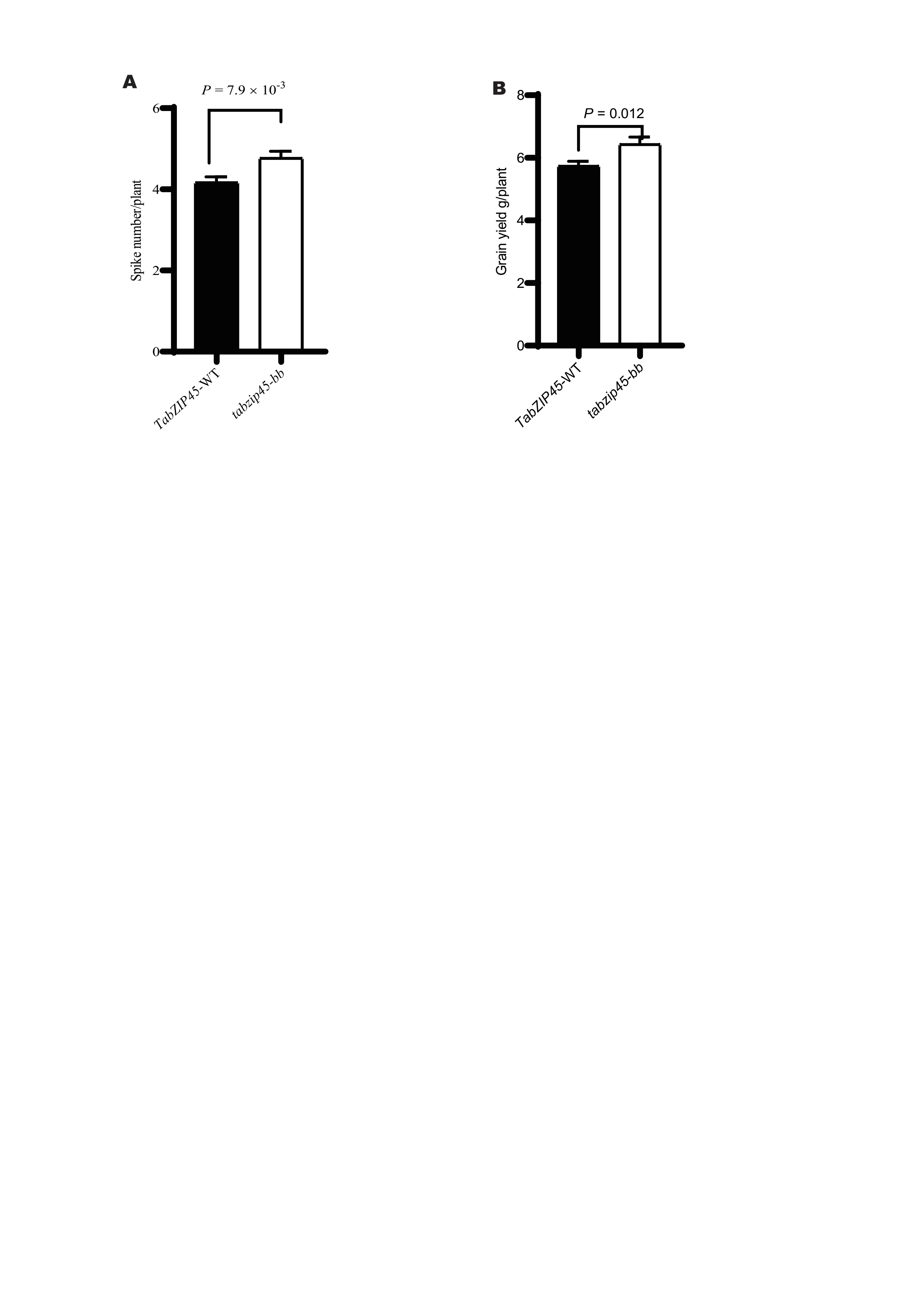

**Fig.S18. Knockout of *TabZIP45-4B* enhanced grain yield in drought condition**

(**A**) Grain yield per plant. (**B**) Spike number per plant. Data in A and B are mean ± S. E. (n ≥ 40). *P* value in D is from a two tailed unpaired test. *TabZIP45*-WT, homozygous wild type offsprings separated from last generation; *tabzip45-bb*, homozygous mutant offsprings separated from last generation.

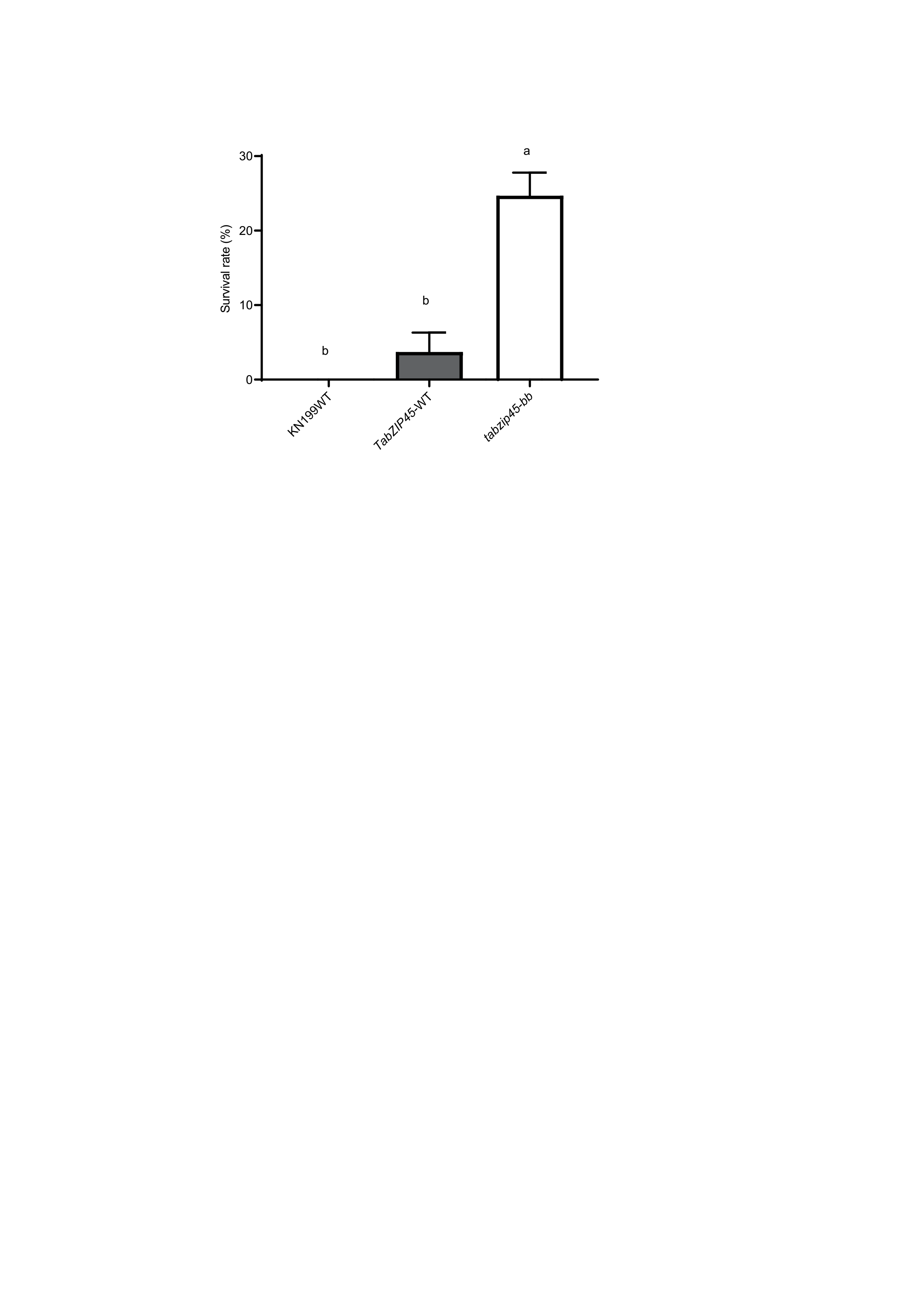

**Fig.S19. *TabZIP45-4B* was crucial for wheat freezing resistance**

Statistical analysis of freezing survival rate in soil experiments. KN199WT, KN199 wild type. *TabZIP45*-WT, homozygous wild type offsprings. *tabzip45-bb*, homozygous mutant offsprings. Data are mean ± S. E. of representative plants (n = 9) from more than three independent experiments. Different letters denote statistically significant differences (*P* < 0.05) from one-way ANOVA multiple comparisons。

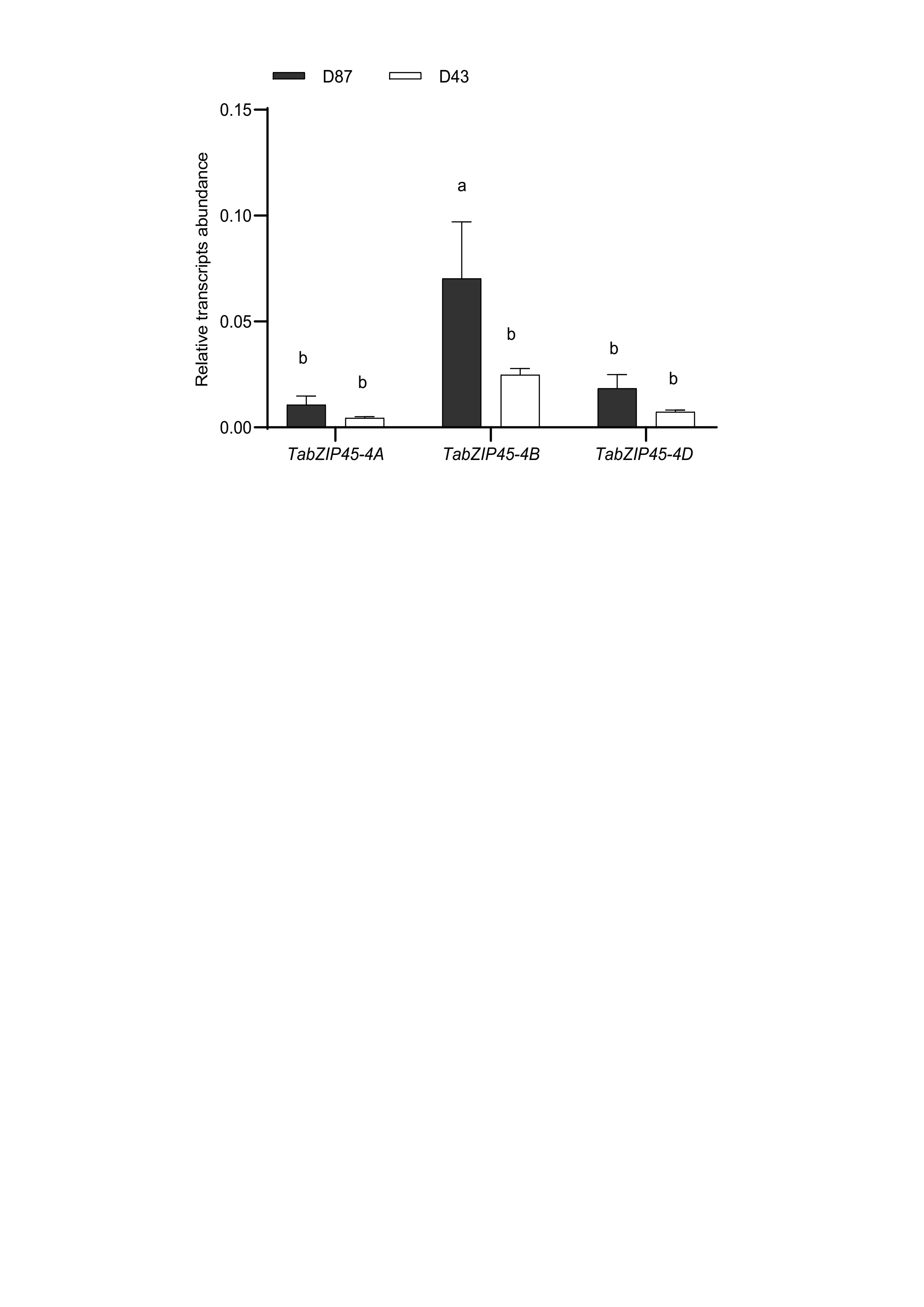

**Fig.S20. Dense planting promoted expression of *TabZIP45***

The above-ground parts of the seedlings were collected 20 days after sowing in 2019-2020 growing season in Beijing. The genes expression level was normalized to that of *Tatublin* as internal control. Data are mean ± S.E. (n = 3). Different letters denote statistically significant differences (*P* < 0.05) from Fisher’s LSD ANOVA multiple comparisons. D43 (43 seeds/m^2^) , D87 (87seeds/m^2^).

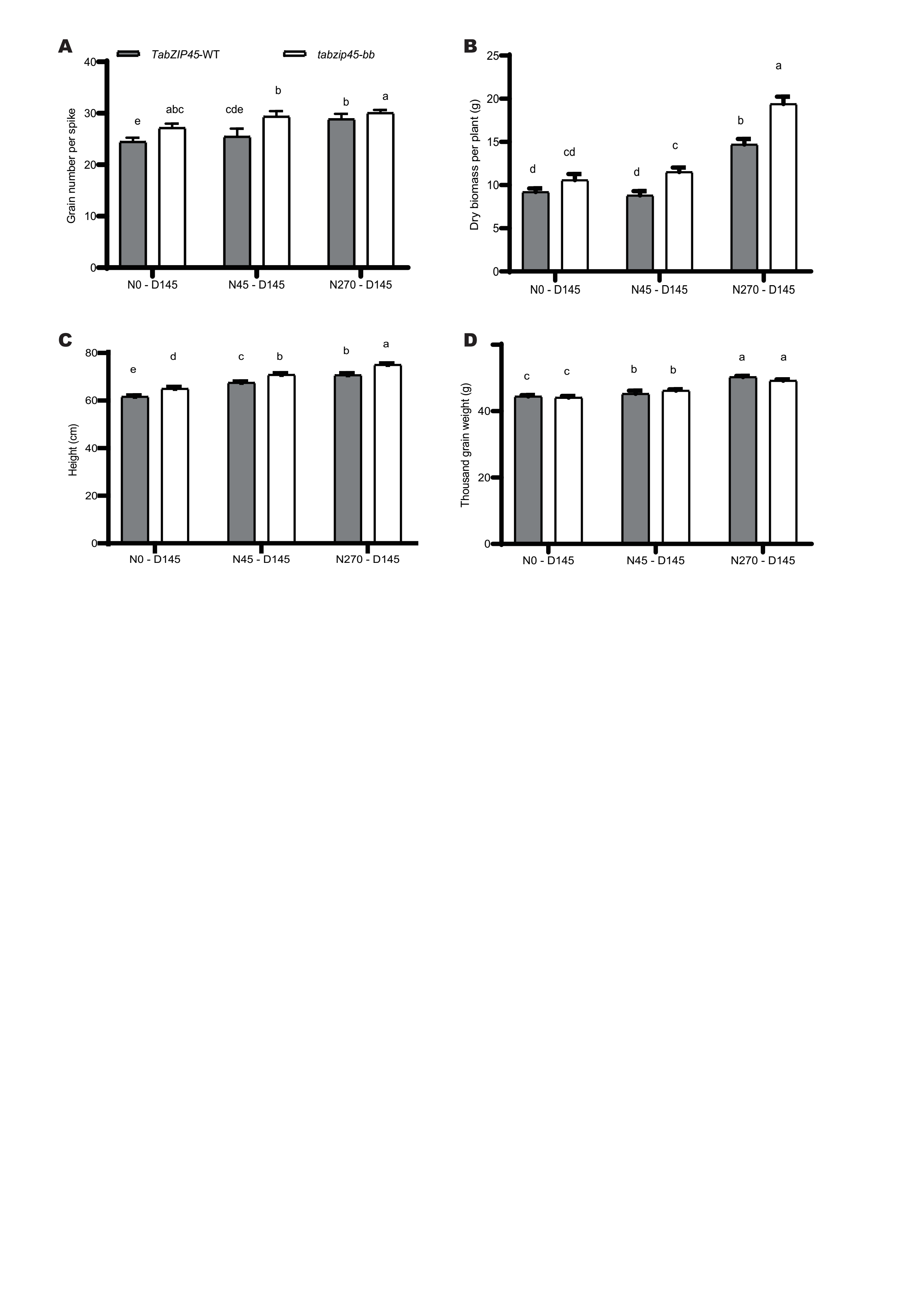

**Fig.S21. Knockout of *TabZIP45-4B* boosted grain yield under high planting density**

**(A)** Grain number per spike. **(B)** Biomass per plant. **(C)** Height. (**D**) 1000-grain weight. *TabZIP45*-WT, homozygous wild type offsprings. *tabzip45-bb*, homozygous mutant offsprings. Nitrogen supply level is 0 kg N/ha, 45 N kg/ha and 270 kg N/ha. Data are mean ± S. E. of representative plants (n = 40) from more than three independent experiments. Different letters denote statistically significant differences (*P* < 0.05) from two-way ANOVA multiple comparisons. In (A-D), N0 (0 kg N/ha), N45 (45 kg N/ha), N270 (270 kg N/ha). D145 (145 seeds/m^2^).
